# Repurposing drugs to treat Amoebic Gill Disease in Atlantic Salmon

**DOI:** 10.64898/2026.06.05.730056

**Authors:** Yee Wan Liu, Abby Lou Elizabeth Bryce, Bachar Cheiab, Brendan Robertson, Kyrie Dickson, Sophie Mouginot, Lindsay Covington, Emma Ohalloran, Philip McGininity, Julie Maguire, Dave O’Neill, Simona Paolacci, Fiona Henriquez, Ralph Bickerdike, Kyrie Dickson, Fintan Egan, Stephanie Linehan, Samantha White, Neil Ruane, Michael Barrett, Martin Llewellyn

## Abstract

*Neoparameoba perurans* causes Amoebic Gill Disease (AGD), a major parasitic disease of marine-phase Atlantic salmon and rainbow trout worldwide. Treatment options are limited to freshwater baths, which are costly at scale and exhibit only limited long-term efficacy. *N. perurans* contains an obligate eukaryotic symbiont, Perkinsela-like organism (PLO). PLO belongs to the class Kinetoplastida, which includes medically and veterinary important parasites such as *Trypanosoma* and *Leishmania*. As such, we hypothesised that trypanocidal drugs developed against other kinetoplastids might also affect *N. perurans*, potentially through disruption of its PLO symbiont, and used this hypothesis as a rationale for prioritising a focused panel of candidate compounds for screening.

A holographic motility-based cytotoxicity assay was established to identify promising candidates *in vitro*, followed by controlled host tolerance testing and finally a field efficacy sea trial using naturally AGD-exposed site in the west of Ireland. Several compounds showed activity *in vitro*, especially miltefosine (EC_50_ 1.84 uM, amoebicidal) and isometamidum (EC_50_ 4.63 uM, amoebostatic). *In vivo* (two intramuscular injections, two weeks apart), miltefosine (Odds Ratio (OR) 0.62), isometamidum (OR 0.61) and benznidazole (OR 0.64) significantly improved gill score over four weeks, with miltefosine showing the largest effect size. Gill parasitaemia, measured via qPCR, was not reduced. Instead, two compounds increased apparent amoeba loads. This work support trypanocidal as potential AGD treatments in the field, although optimisation of dosing, delivery and mode of action requires further study.

## 1. Introduction

Amoebic gill disease (AGD), caused by the marine amphizoic *amoeba Neoparamoeba perurans*, is among the most economically significant diseases affecting farmed Atlantic salmon (*Salmo salar*) worldwide [1]. Outbreaks lead to substantial losses through direct mortality, reduced growth performance, compromised welfare, and the labour and infrastructure costs associated with repeated treatments [2], [3], [4]. In severe cases, mortality rates can approach 80% [5], [6]. Current management relies largely on freshwater bathing and hydrogen peroxide exposure, which provide only transient relief, require repeated handling, and impose significant operational and welfare burdens [7], [8], [9]. As salmon aquaculture continues to expand there is a pressing need for more effective, scalable, and welfare-compatible therapeutic strategies.

A defining feature of *N. perurans* is its complex symbiotic biology, which may offer novel opportunities for targeted intervention. Endosymbiosis is recognised as a powerful driver of rapid evolutionary innovation, enabling the emergence of new cellular functions through the integration of phylogenetically distinct organisms [10], [11]. Symbiotic dependencies are widespread among protists and can generate exploitable vulnerabilities. *N. perurans* is consistently associated with bacterial consortia, particularly *Vibrio* spp., and attempts to establish axenic cultures have been unsuccessful, with surface-disinfected isolates repeatedly reintroducing bacteria into culture [12], [13], [14]. Experimental work on related amoebae indicates that depletion of associated bacterial communities can reduce growth and virulence, highlighting the functional importance of amoeba–bacteria interactions for physiology and pathogenicity [15], [16].

In addition to bacterial associates, most *Neoparamoeba* spp. harbour a eukaryotic endosymbiont, the Perkinsela-like organism (PLO), a kinetoplastid that resides adjacent to the host nucleus [17], [18], [19]. Phylogenetic analyses place the PLO within Kinetoplastida, with basal Perkinsela lineages forming a sister group to Ichthyobodo, indicating early divergence from trypanosomatid ancestors [18], [20], [21]. Strong congruence between host and endosymbiont phylogenies supports an ancient origin followed by co-speciation [19], [20], [22]. Genomic analyses reveal reductive evolution and metabolic interdependence, with the PLO possessing one of the smallest known kinetoplastid genomes [23], [24]. Although the precise role of the PLO in *N. perurans* biology remains unresolved, this obligate association suggests that disruption of the PLO–amoeba symbiosis could compromise *N. perurans* viability or virulence.

Kinetoplastids include several major human and veterinary pathogens, such as *Trypanosoma cruzi*, *T. brucei*, and *Leishmania* spp., for which multiple licensed drugs already exist. Conventional phenotypic drug discovery pipelines in kinetoplastid disease routinely screen libraries of tens of thousands to millions of compounds to identify active hits [25], [26], a scale of effort that is not feasible for a disease of farmed fish such as AGD. The presence of the PLO in *N. perurans* provides a biologically motivated rationale for a much more focused approach, allowing a panel of fewer than 20 trypanocidal candidates to be prioritised for screening against the amoeba. Repurposing such compounds offers an attractive strategy for AGD control by exploiting shared biology while avoiding the cost and time required for *de novo* drug development [27], [28], [29]. We emphasise that this rationale is hypothesis-driven; direct evidence that candidate drugs act via the PLO in *N. perurans* remains to be established. Trypanocidal drugs encompass diverse chemical classes and mechanisms, including alkylphosphocholines (e.g. miltefosine), aminoglycosides (e.g. paromomycin), benzoxaboroles targeting mRNA processing (e.g. acoziborole), diamidines targeting kinetoplast DNA, and nitroheterocyclic prodrugs requiring parasite-specific activation [30], [31], [32], [33], [34]. Several of these mechanisms directly affect organelles, nucleic acids, or metabolic processes that are distinctive to kinetoplastids, making them plausible candidates for targeting the PLO. Importantly, aquaculture also represents a translational context in which successful repurposing could support economically viable production at scale, with potential downstream benefits for drug accessibility in other kinetoplastid disease settings.

A key challenge in evaluating candidate compounds against *N. perurans* is the lack of robust quantitative drug assays compatible with non-axenic cultures. The presence of cocultured bacteria and biofilm fragments can confound conventional viability readouts and reduce reproducibility [13], [14], [35]. Moreover, *N. perurans* readily forms pseudocysts under stress, complicating interpretation of assays based solely on morphology or terminal membrane-integrity stains [36]. Traditional approaches using trypan blue or neutral red rely on manual microscopic assessment of dye exclusion and morphological criteria to determine viability misclassification [37], [38]. In these assays, rounded cells have frequently been classified as non-viable; however, pseudocysts may later revert to trophozoites, leading to potential misclassification [36], [37]. To reduce operator-dependent variability inherent to haemocytometer-based counting, fluorescence-based membrane integrity assays such as propidium iodide (PI) provide quantitative readouts that can be acquired using flow cytometry or automated fluorescence imaging platforms [39], [40]. Furthermore, multiplex approaches combining membrane integrity (cellTox Green) and ATP-based metabolic (cellTitre Glo) readouts have been validated for *N. perurans* and are proposed to differentiate amoebostatic from amoebicidal effects [41]. Although these approaches claim to improve quantification and throughput, their reliability in non-axenic *N. perurans* cultures may be constrained by bacterial interference and the biological complexity of pseudocyst formation.

Digital holographic microscopy provides a complementary, time-resolved approach by enabling label-free tracking of individual cells and quantification of motility and morphology. Amoeba motility is closely linked to physiological status, with loss and recovery of movement reflecting stress, survival, or death [42], [43]. The HoloMonitor® platform allows continuous imaging under standard culture conditions and automated single-cell analysis, facilitating discrimination between transient immotility and irreversible damage through recovery assays [44]. When integrated with established fluorescence-based viability assays, this approach offers a robust framework for prioritising candidate compounds for *in vivo* evaluation.

Ultimately, translation of *in vitro* activity into effective AGD control requires testing under biologically realistic conditions. AGD develops within a complex gill disease context shaped by environmental, microbial, and host factors, and therapeutic success must be demonstrated at the level of gill pathology and amoeba burden. Natural exposure systems with predictable AGD pressure therefore provide critical validation. The Lehanagh Pool field site, which experiences recurring seasonal AGD epidemic, has been used to investigate host–microbiome–pathogen interactions under natural challenge conditions [3]. Coupling quantitative *in vitro* screening with such a natural exposure system offers a unique opportunity to assess the efficacy and tolerability of repurposed trypanocidal drugs across a broad translational pathway.

The aim of this study was to evaluate whether existing trypanocidal drugs could be repurposed to treat AGD, using the presence of the kinetoplastid endosymbiont of *N. perurans* as a rationale for prioritising candidate compounds. Specifically, we (i) compared complementary *in vitro* drug-screening approaches suitable for non-axenic *N. perurans* cultures, (ii) screened mechanistically diverse anti-kinetoplastid compounds for amoebicidal or amoebostatic activity, (iii) assessed host tolerance of prioritised candidates in Atlantic salmon, and (iv) evaluated therapeutic efficacy under natural AGD exposure using clinical gill scoring and quantitative PCR.

## 2. Materials and Methods

### 2.1. In vitro drug assays

#### 2.1.1 Cell culturing

*Neoparamoeba perurans* cultures were kindly provided by Dr Roderick Williams and Prof Fiona Henriquez-Mui (University of the West of Scotland) and were originally isolated from CEFAS gill-swab samples. Cultures were maintained at 17 °C in T10 vented tissue-culture flasks containing 10 mL of medium, replaced at least weekly. The medium consisted of artificial seawater (35 ppt) supplemented with 1% (w/v) yeast extract and 1% (w/v) malt extract. Media were autoclaved, and salinity was verified post-sterilisation using a handheld refractometer within a laminar-flow hood.

For subculture, spent media from all flasks were pooled to recover floating amoebae, filtered through a 15 µm cell strainer to remove biofilm fragments, and centrifuged at 1,500 × g for 15 min at 4 °C. Pelleted cells were resuspended in 50 mL fresh medium and evenly distributed into five T75 vented flasks. Pooling and redistribution were used to minimise inter-flask variation arising from flask-specific microbial communities, which is critical for downstream drug screening.

#### 2.1.2. Drug assay set up using Propidium iodide staining

To assess drug efficacy against *N. perurans,* compounds were prepared as ten-fold serial dilutions in culture medium. Tested compounds included: alkylphosphocholine (miltefosine), phenanthridine (isometamidium), aminoglycoside (paromomycin), benzoxaboroles (AN11736, AN5568), diamidines (pentamidine, DB75, DB829, diminazene), nitroheterocyclics (nifurtimox, benznidazole, fexinidazole), polyene (amphotericin B), and suramin.

Amoebae were detached by gentle tapping, collected, centrifuged at 1,500 × g for 15 min, and resuspended to a density of 2.0 × 10⁴ cells mL⁻¹. Aliquots of 2,000 cells were dispensed per well and incubated with compounds for 72 h. Cells were then stained with 20 µM Propidium iodide (**Figure S1**) for 30 min and examined using a Leica DiM8 microscope. Viability was quantified using a disposable haemocytometer (Fast-Read 102®, Biosigma, Italy).

#### 2.1.3. Dual-assay approach using CellTox Green and CellTiter-Glo

A dual-assay approach adapted from Botwright *et al*. (2020) was employed to assess N. perurans viability. CellTiter-Glo® quantified ATP as a proxy for metabolic activity, while CellTox™ Green detected DNA from membrane-compromised cells. As cultures were non-axenic, two sample types were analysed: (i) non-homogenised amoebae with associated bacteria, and (ii) homogenised amoebae releasing intracellular and extracellular bacteria.

Amoebae were disrupted using an Isobiotec Balch homogeniser (4 µm ball, 20 strokes; **Figure S2**). Both sample types were subjected to three treatments: untreated (medium only), 4% formaldehyde, and lysis buffer (positive dead-cell control). To determine assay sensitivity, 3,250 cells per well were seeded in 96-well plates and serially diluted two-fold under each treatment in quadruplicate. CellTox™ Green fluorescence was additionally confirmed by fluorescence microscopy using a Leica DiM8 microscope.

Three treatments were included for validation: no treatment, 4% formaldehyde, and the lysis buffer supplied with CellTox Green designed to act as a negative control. Two sample types were also prepared: non-homogenised samples contained amoebae with intracellular and extracellular bacteria, and homogenised samples contained only bacteria after mechanical disruption of amoebae. These preparations were used to test whether bacterial activity influenced the assay readouts.

Statistical analyses were conducted in R (v. 4.4.2). Linear models and one-way ANOVA with Tukey’s HSD post-hoc tests were used where appropriate. Data were visualised using ggplot2, and model assumptions were verified using residual diagnostics, histograms, and Q-Q plots.Statistical analyses were conducted in R (v. 4.4.2). Linear models and oneway ANOVA with Tukey’s HSD post-hoc tests were used where appropriate. Data were visualised using ggplot2, and model assumptions were verified using residual diagnostics, histograms, and Q-Q plots.

#### 2.1.4 Drug assay setup using holographic microscopy

##### 2.1.4.1 Experimental setup

*N. perurans* were seeded at 10,000 cells per well in Sarstedt Lumox® 24-well plates (cat. #94.6000.014) with 2,000 µL complete medium and incubated overnight for adherence. Floating bacteria and debris were removed by replacing the upper medium layer, followed by an additional overnight incubation. Drug treatments were applied according to the plate layout shown in Figure 1, with serially diluted compounds added to triplicate wells at final concentrations of 0.625, 1.25, 2.5, 5, 10, and 20µM, alongside negative controls. Serially diluted drugs (500 µL) were then added to each well (final volume 2,500 µL). Plates were sealed with HoloLid™ (cat. #71130) to minimise vibration and condensation artefacts. Live-cell imaging commenced immediately using a HoloMonitor M4 system (Phase Holographic Imaging, PHI AB, Sweden). Images were captured at two random positions per well every 2 min over 72 h.

**Figure 1.**
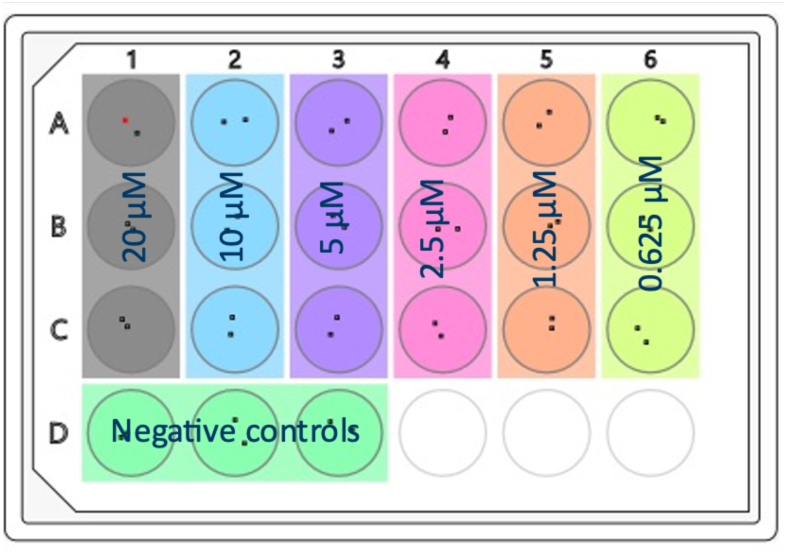
*Neoparamoeba perurans* were treated with drugs at concentrations ranging from 0.625 µM to 20 µM.

##### 2.1.4.2 Single-cell tracking analysis and statistics

Single-cell tracking was performed using HoloStudio 2.4 software. For each well, five randomly selected amoebae were tracked frame-by-frame. When fewer than five amoebae were present, missing values were assigned zero motility. Migration directness (migration/motility) was used as the primary response metric to correct for non-voluntary movement. Representative trajectories are provided in **S3 (Supplementary materials 3)**.

Drug responses were assessed over 72 h using 4-h analysis windows at 0, 24, 48, and 72 h. Effects of drug concentration and time were analysed using generalised linear models with a Gamma distribution and log link. Likelihood-ratio tests assessed time–drug interactions and the independent contribution of time and concentration. Analyses were performed in R (v. 4.4.2) using glmmTMB, with tidyverse and ggplot2 for data processing and visualisation. Dose–response curves were generated using the drc package to estimate EC₅₀ values and percentage inhibition relative to untreated controls.

### 2.2. In vivo drug assays

This section describes pilot tolerance (Trial 1) and field (Trial 2) efficacy trials evaluating candidate drugs active against *N. perurans* in Atlantic salmon.

#### 2.2.1 Trial 1: Experimental organism and rearing

Atlantic salmon (Salmo salar) post-smolts (weight 110 g) were housed in five 1,000-L tanks (50 fish per tank) at Bantry Marine Research Station (BMRS), Ireland, in April 2022. Seawater (75 m³ h⁻¹) was filtered, UV-treated, ozonated, and sand-filtered prior to tank entry (Figure 23.1). Tanks were aerated and oxygenated, and fish were fed ad libitum. All procedures were conducted under HPRA authorisation AE19114.

**Figure 2.**
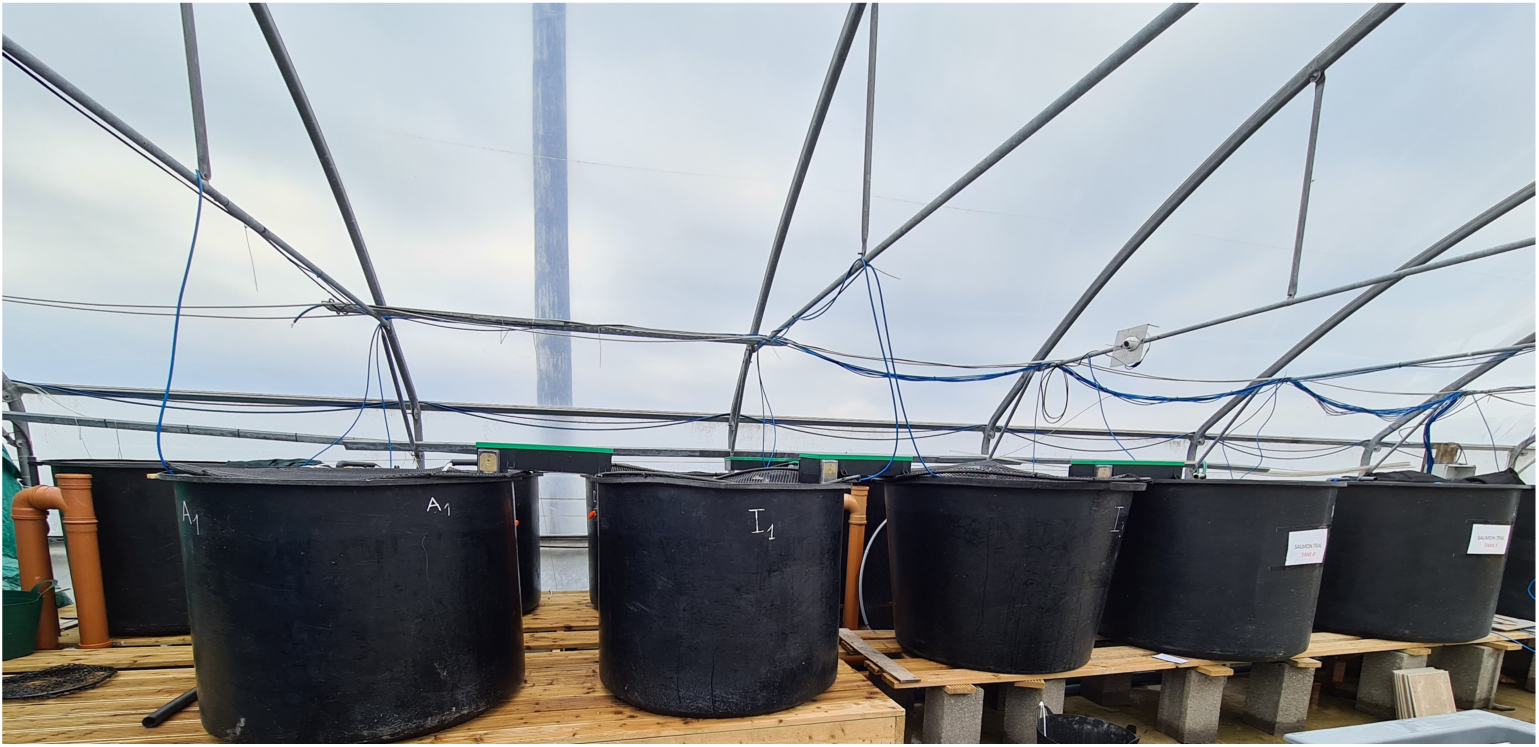
Tank set up for in vivo tolerance assessment in Bantry Marine Research Station (BMRS), Gearhies, Bantry, Co. Cork, Ireland (P75 AX07), within Dr Julie Maguire’s research group. Five tanks were used, each representing a separate treatment group and stocked with 50 healthy Atlantic salmon (Salmo salar) with a mean weight of approximately 100 g.

#### 2.2.2. Trial 1: Compound administration and dosing

Four candidate drugs (miltefosine, isometamidium, benznidazole, nifurtimox) were selected based on prior *in vitro* screening. Treatments were randomly assigned to tanks. Fish received 1.0 mL intramuscular injections on Days 1, 3, and 5. Controls received vehicle only (PBS + 1% DMSO). Doses were: miltefosine 2.5 mg kg⁻¹, isometamidium 1.0 mg kg⁻¹, benznidazole 41.66 mg kg⁻¹, and nifurtimox 5.18 mg kg⁻¹ (reduced from 10 mg kg⁻¹ due to solubility). Dosages were based on established regimens in other species. Fish were anaesthetised using MS-222 prior to injection.

#### 2.2.3. Trial 1: Endpoints and monitoring

Mortality was recorded multiple times daily. Every other day, three fish per tank were euthanised, and blood was collected for serum cortisol analysis. Cortisol concentrations were quantified in duplicate using a competitive ELISA (Enzo Life Sciences) and calculated using a four-parameter logistic model.

#### 2.2.4. Trial 1: Statistical analysis

Mortality was analysed using a binomial GLM and Mantel–Haenszel chi-squared tests stratified by day. Cortisol concentrations were analysed using GLMs with treatment and sampling day as predictors. Model selection used likelihood-ratio tests, and diagnostics were conducted using DHARMa. Analyses were performed in R (v. 4.3.1).

### 2.3. Trial 2: In vivo efficacy trial in AGD-infected S. salar

#### 2.3.1. Trial 2: Experimental setup

Irish-strain Atlantic salmon were reared from eyed eggs (Mowi Ireland) to post-smolts at the Marine Institute Newport Facility and transferred to the Lehanagh Pool Marine Research Site (Galway, Ireland) in April 2023. The site comprises 21.7 ha with water temperatures of 11–17 °C. The trial was conducted under HPRA authorisation AE19121/P006. Fish were fed with commercial pellets (EWOS Cargill) twice daily by hand to apparent satiation.

#### 2.3.2. Trial 2: AGD monitoring and treatment

Natural *N. perurans* infection was monitored weekly by qPCR of gill swabs. Infection was first detected at Week 5 and reached 88% prevalence by Week 8. Fish were distributed into six 4 × 4 m sentinel pens (298 fish per pen), each containing two treatments (n = 149 per treatment) identified by fin clipping (see Figure 3 for sentinel-pen layout). Each set of six sentinel pens were placed within a larger 50 m circumference marine pen, which contained a net to a depth of 10 m for additional security against the escape of experimental fish.

**Figure 3.**
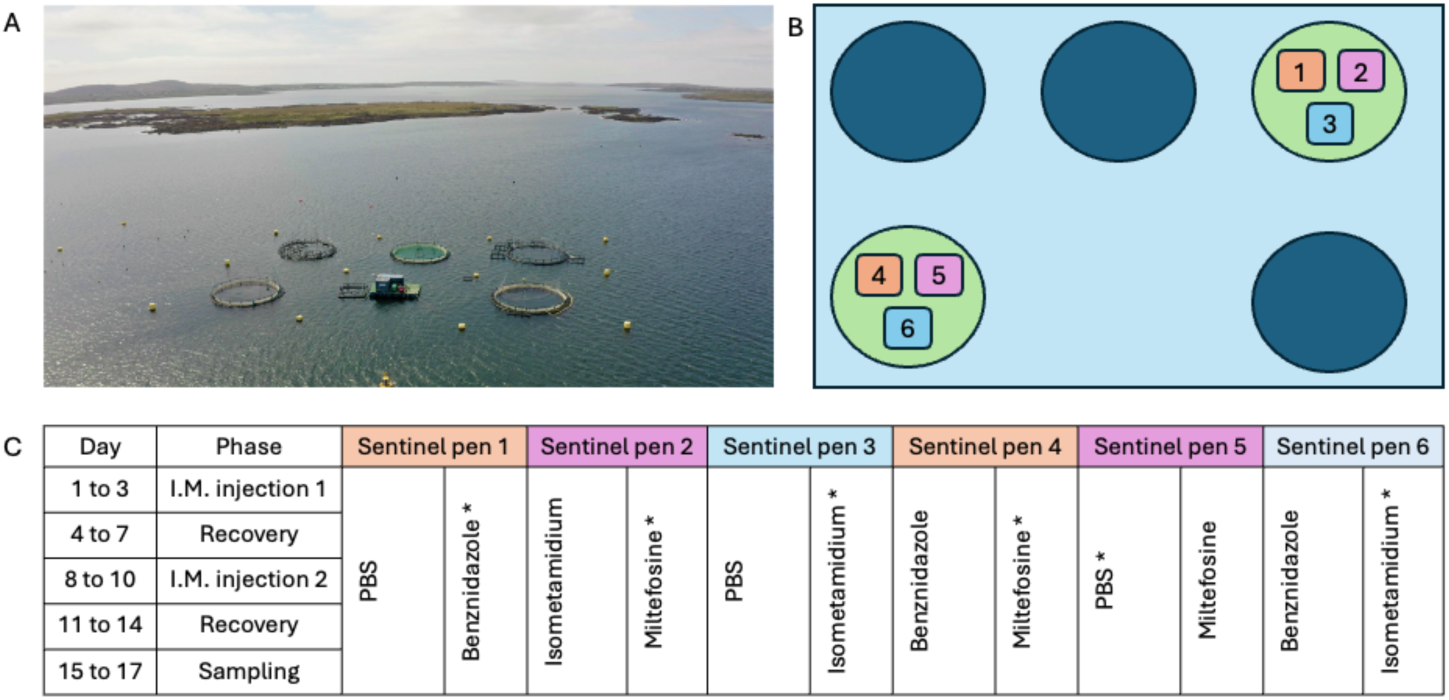
Trial set-up at the Marine Institute’s Lehanagh Pool, Bertraghboy Bay, Connemara, Co. Galway (July 2023). Six 4×4 m sentinel pens (numbered 1–6) were positioned inside a larger 50-m-circumference circular net pen. Each sentinel pen held 298 Atlantic salmon and contained two treatments (n = 149 per treatment), distinguished by fin-clipping one treatment group (see Methods). Arrows indicate the locations of sentinel pens 1–3 and 4–6.

Four treatments were applied in triplicate: untreated control, benznidazole, isometamidium, and miltefosine (doses as in Trial 1). Two intramuscular injections were administered one week apart (Days 1–3 and 8–10).

#### 2.3.3. Trial 2: Endpoints and monitoring

One week post-treatment, fish were euthanised for gill scoring, gill swabbing, and morphometric measurements. Condition factor (K) was calculated:

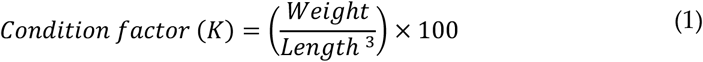

**Equation 1**. Fulton’s Condition Factor [45].

DNA was extracted from gill swabs (Qiagen MiniPrep) and *N. perurans* load quantified by qPCR targeting the 18S rRNA gene [46], with all samples run in technical triplicate alongside standard curves.

#### 2.3.4. Trial 2: Statistical analysis

Gill scores were analysed using proportional odds logistic regression with random effects for cage and sentinel pen. Associations between gill score, amoebae load, and condition factor were assessed using linear and linear mixed-effects models, accounting for sentinel pen and qPCR batch effects. Model selection used likelihood-ratio tests. All analyses were conducted in R (v. 4.3.2).

Equations 2–8 describe the full statistical model framework.

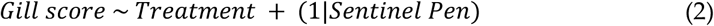

**Equations 2**. a proportional odds logistic regression model describing all fish for gill score (a ranked categorical variable) is depending on both treatment as a fixed effect and sentinel pen as a random variable.

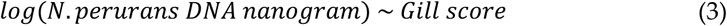

**Equations 3**. a simple linear regression relating the log-transformed N. perurans DNA load (ng) to gill score (ordinal predictor).

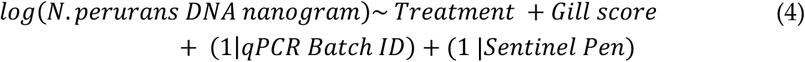

**Equations 4**. a linear mixed-effects model for the log-transformed N. perurans DNA load with treatment and gill score as fixed effects, and random intercepts for qPCR batch and sentinel pen to account for laboratory and pen-level clustering.

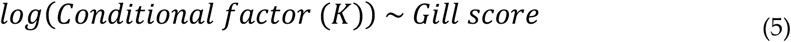

**Equations 5**. a simple linear regression relating the log-transformed condition factor (K) to gill score (ordinal predictor).

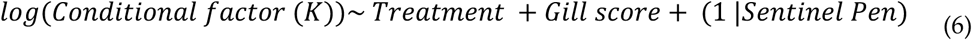

**Equations 6**. an additive linear mixed-effects model for the log-transformed condition factor (K), including treatment and gill score as fixed effects and a random intercept for sentinel pen.

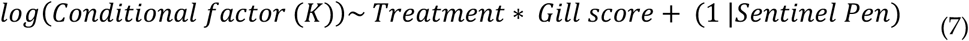

**Equations 7**. an interactive linear mixed-effects model for the log-transformed condition factor (K), including treatment and gill score as fixed effects and a random intercept for sentinel pen.

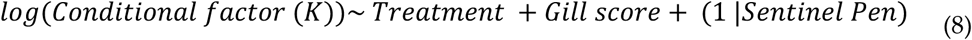

**Equations 8**. a linear mixed-effects model for the log-transformed condition factor (K), with treatment as fixed effects and a random intercept for sentinel pen.

## 3. Results

### 3.1. Eight trypanocidal drug classes of trypanocidal were tested on Neoparamoeba perurans using fluorescence microscopy with propidium iodide

The ability of propidium iodide (PI) to indicate cell viability was confirmed using fixed and untreated amoebae as positive and negative controls, respectively (Figure 4). In fixed controls, PI penetrated non-viable cells and produced strong red fluorescence, while untreated amoebae remained unstained. Based on this validation, a drug was considered effective if it induced pseudocyst formation and PI uptake at concentrations of 20 µM or 2 µM, resembling the positive control. Conversely, drugs were classified as ineffective if most treated amoebae retained pseudopodia and did not stain with PI, similar to the neative control.

**Figure 4.**
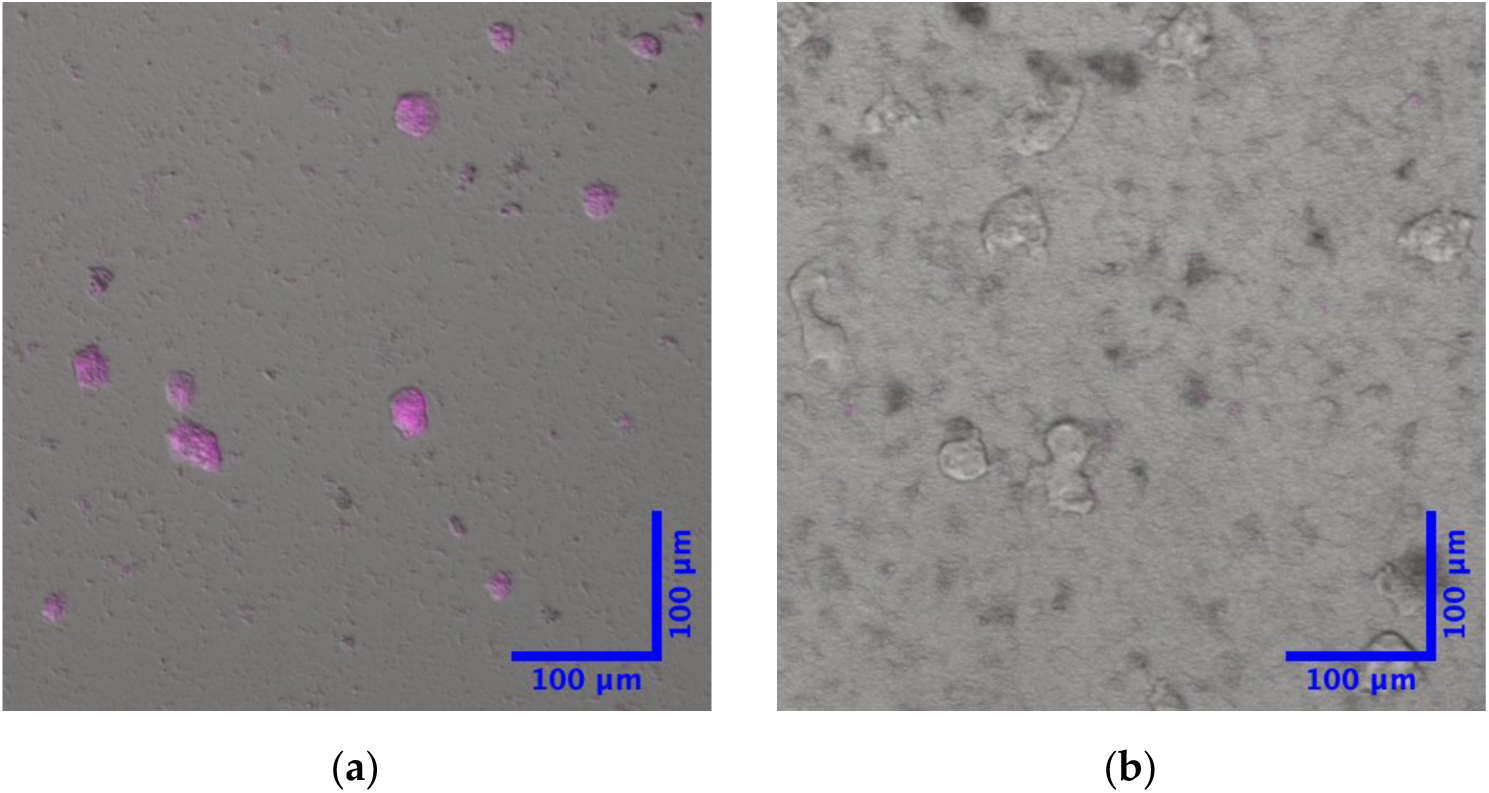
Fluorescence microscopy of (a) positive (formaldehyde-fixed amoeba) and (b) negative controls (non-treated amoeba). These images were generated by merging the red fluorescence channel with the bright field channel. To distinguish dead amoeba, cells were stained with 20 µM propidium iodide. Cells are considered to be dead if they are stained in red (filtered pink), whereas viable cells are not stained.

Eight trypanocidal drug classes were tested against *N. perurans* using this assay (**Figure S4**). Applying the criterion established above (pseudocyst formation with PI uptake at ≤20 µM), five compounds were classified qualitatively as effective: miltefosine, isometamidium, nifurtimox, benznidazole, and suramin. The remaining compounds induced little or no PI uptake and resembled the untreated control. Attempts to obtain quantitative counts using a haemocytometer were unsuccessful due to consistently low cell numbers per well, and these data were therefore excluded from comparative analysis.

### 3.2. CellTox Green assays

CellTox Green fluorescence was absent in live *N. perurans*, while 4% formaldehyde-fixed amoebae showed strong staining, confirming reliable detection of dead cells. In contrast, treatment with the lysis buffer failed to produce detectable fluorescence even after 72 h, and amoebae appeared morphologically intact, indicating that the lysis buffer was ineffective as a positive control in this system (Figure 5).

**Figure 5.**
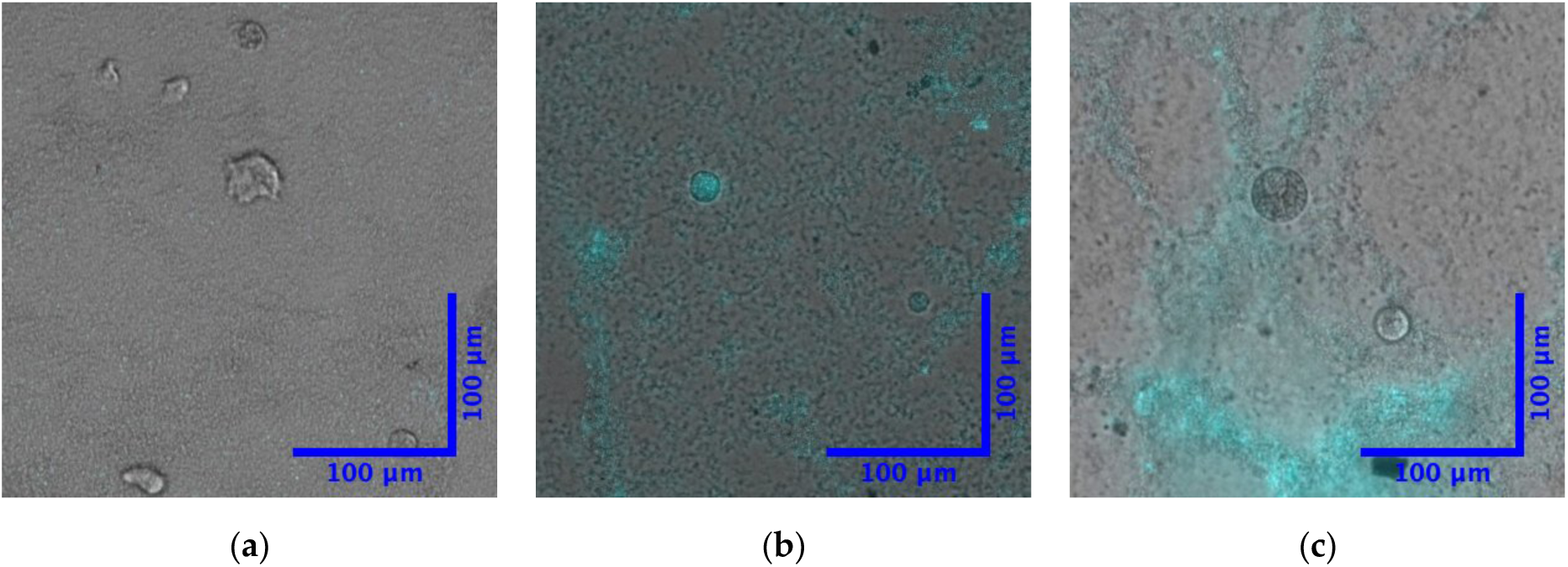
*Neoparamoeba perurans* subjected to three different treatments: (a) no treatment and (b) 4% formaldehyde fixation, and (c) Promega lysis buffer, followed by a 72 hrs exposure to 1x CellTox Green dye, where dead cells are indicated by green staining visualised with a cyan filter). Images were captured at a magnification of 10x using a Leica DiM8 microscope.

Quantitative fluorescence measurements were consistently higher in non-homogenised than in homogenised samples across treatments and cell densities, particularly at 3,250 and 1,625 cells per well. However, no significant differences were detected between non-homogenised and homogenised samples for untreated controls, formaldehyde-fixed samples, or lysis buffer–treated samples at either cell density, indicating a substantial bacterial contribution to the fluorescence signal that could mask amoeba-specific effects (Figure 6).

**Figure 6.**
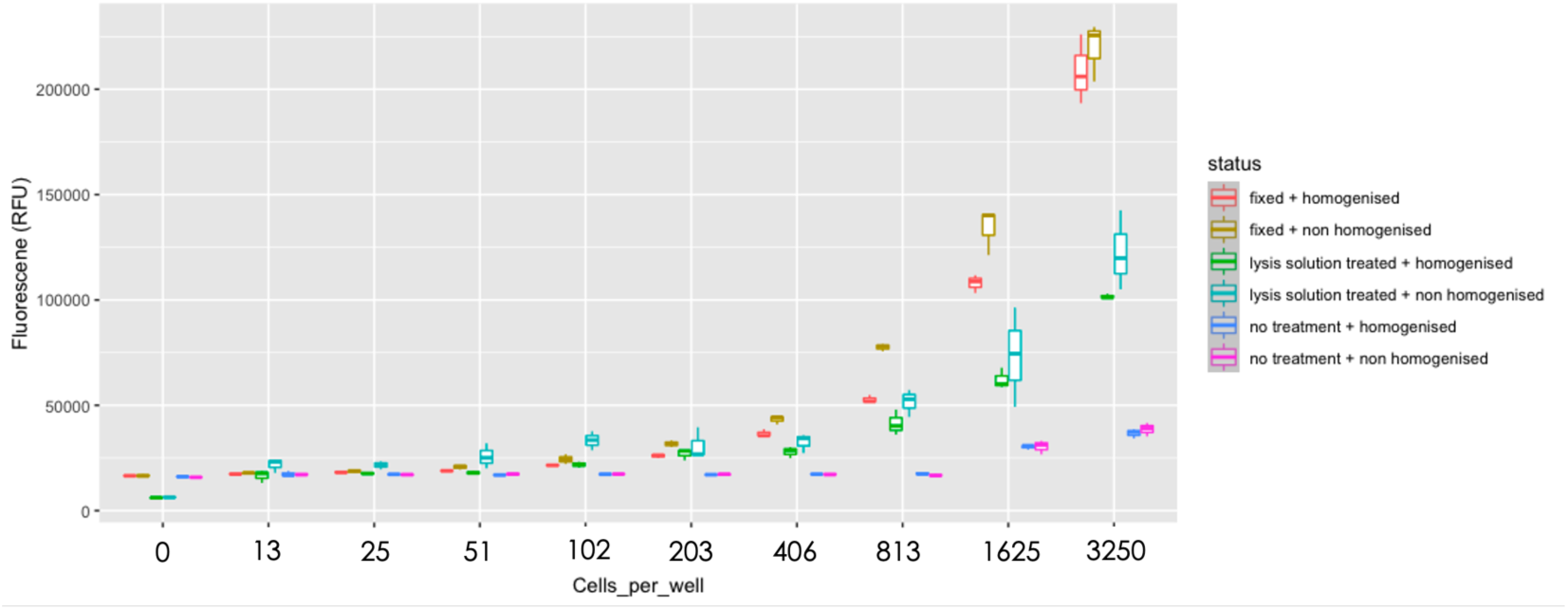
Cytotoxicity levels were assessed using the Promega CellTox Green Cytotoxicity assay for non-axenically cultured *Neoparamoeba perurans*. These amoebae were either non-homogenised (containing both amoebae and their intracellular and extracellular bacteria) or homogenised (containing only intracellular and extracellular bacteria). These non-homogenised and homogenised amoebae were then subjected to the treatments including no treatment, lysis solution (Promega lysis buffer) treatment, and fixation with 4% formaldehyde. On the x-axis, a 2-fold serial dilution of cell numbers is displayed, ranging from 3250 cells/well to 0 cells/well. The y-axis represents fluorescence levels (RFU) measured using the Promega CellTox Green Cytotoxicity assay kit and a plate reader.

Formaldehyde-fixed samples showed significantly higher cytotoxicity than untreated controls in both non-homogenised (3,250 cells/well: p = 0.0074; 1,625 cells/well: p = 0.0029) and homogenised samples (3,250 cells/well: p = 0.0047; 1,625 cells/well: p = 0.012), confirming that the assay detected cell death but reflected combined amoebal and bacterial mortality. In contrast, no significant differences were observed between lysis buffer– treated and untreated samples in either non-homogenised or homogenised cultures at either cell density. Cytotoxicity values were consistently higher in formaldehyde-fixed samples than in lysis buffer–treated samples, although these differences were not statistically significant (Figure 7).

**Figure 7.**
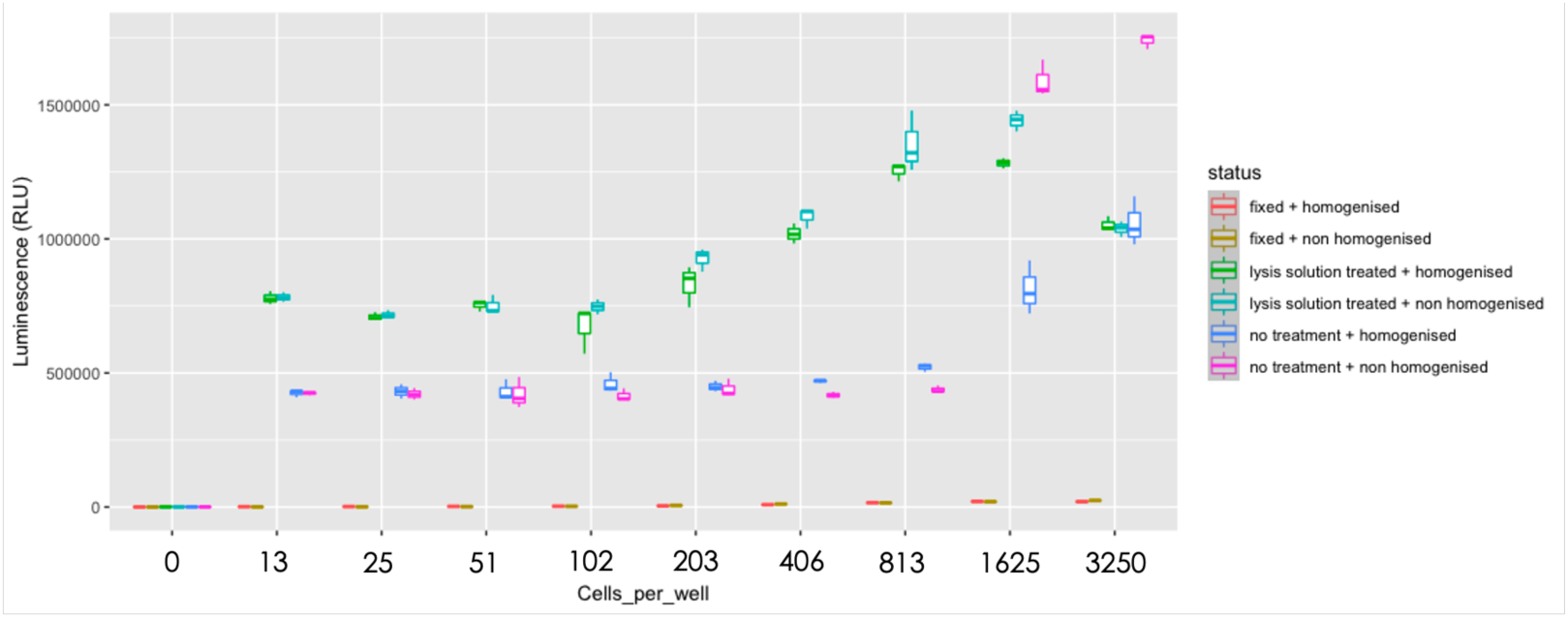
Metabolic ATP production activity levels were assessed using the Promega CellTitre Glo Viability assay for non-axenically cultured Neoparamoeba perurans. These amoebae were either non-homogenised (containing both amoebae and their intracellular and extracellular bacteria) or homogenised (containing only intracellular and extracellular bacteria). These non-homogenised and homogenised amoebae were then subjected to the treatments including no treatment, lysis solution (Promega lysis buffer) treatment, and fixation with 4% formaldehyde. On the x-axis, a 2-fold serial dilution of cell numbers is displayed, ranging from 3250 cells/well to 0 cells/well. The y-axis represents luminescence levels (RLU) measured using the Promega CellTitre Glo Viability assay and a plate reader.

Overall, these results demonstrate that formaldehyde fixation was an effective positive control for CellTox Green–based detection of cell death, whereas the lysis buffer was ineffective in *N. perurans* cultures and unsuitable for use in this assay.

### 3.3. CellTiter-Glo assays

ATP levels measured using CellTiter-Glo were generally higher in non-homogenised samples than in homogenised samples in the untreated group, particularly at 3,250 and 1,625 cells per well (Figure 8). At 3,250 cells per well, this difference was not significant (p-value = 0.14), whereas at 1,625 cells per well it was significant (p = 0.038). This suggested that 1,625 cells per well may approximate an optimal cell density for distinguishing amoebal ATP from bacterial background. However, this pattern was not consistent, as the higher cell density did not reach significance. Moreover, at cell densities below 1,625 cells per well, ATP levels in non-homogenised samples were unexpectedly lower than those in homogenised samples, indicating substantial variability in the assay. Together, these results demonstrate that bacterial ATP contributed strongly to the luminescence signal and could obscure amoeba-specific measurements, particularly at lower cell numbers.

**Figure 8.**
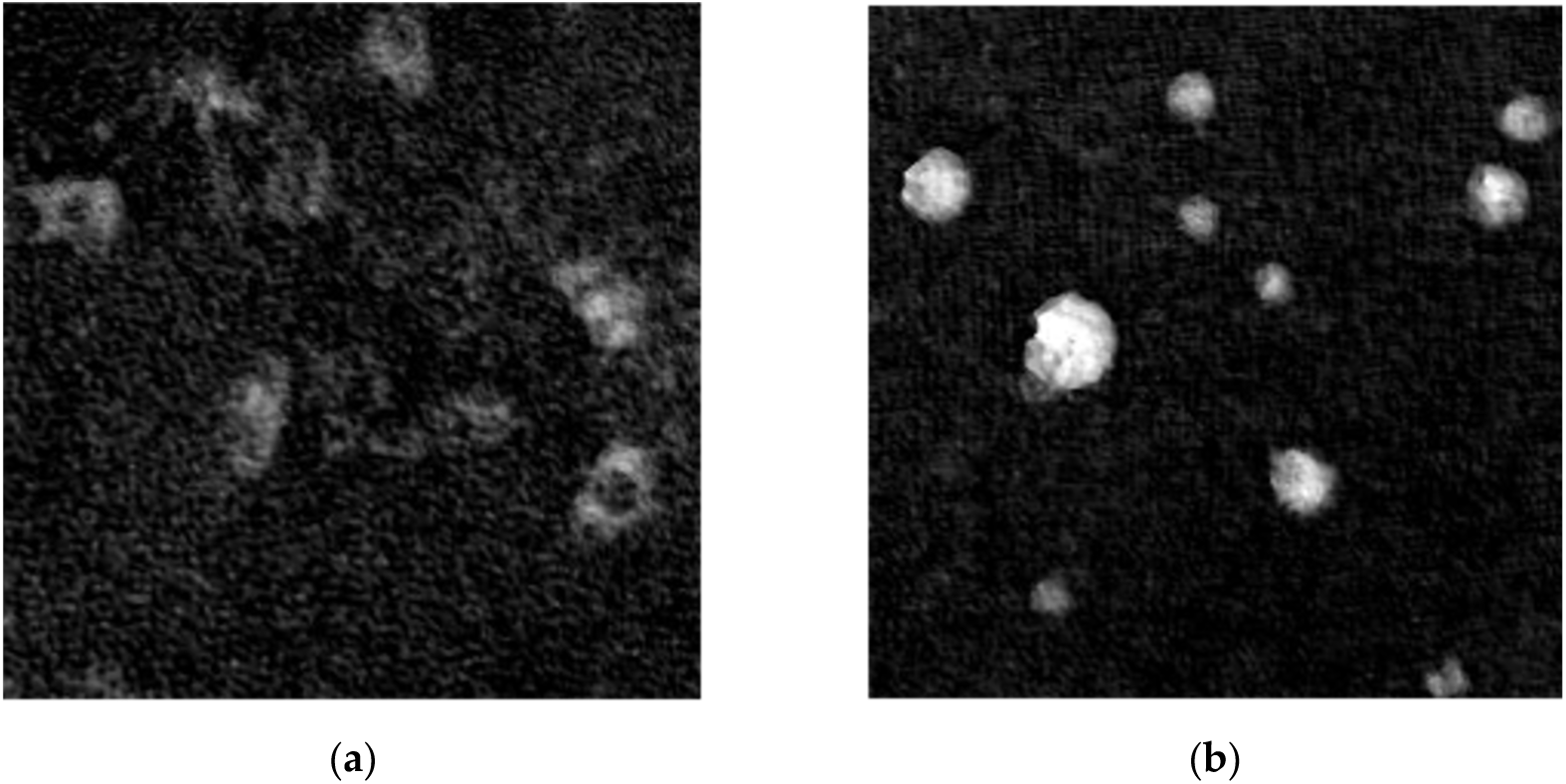
*Neoparamoeba perurans* has two observable forms in culture: a) trophozoite, and b) pseudocyst. Images were captured using a label-free holographic microscopy, Holomonitor.

ATP levels were significantly reduced in formaldehyde-fixed samples compared with untreated controls, confirming that fixation effectively suppressed metabolic activity. In non-homogenised samples, this reduction was significant at both 3,250 cells per well (p = 0.0059) and 1,625 cells per well (p = 0.00057). In homogenised samples, a significant reduction was observed at 3,250 cells per well (p = 0.047) but not at 1,625 cells per well (p = 0.49). These findings indicate that fixation reduced ATP levels in both amoebae and bacteria, confirming that the assay response was not amoeba-specific.

In contrast, lysis buffer treatment did not consistently reduce ATP levels relative to untreated controls. No significant differences were observed in non-homogenised samples at either 3,250 cells per well (p = 0.88) or 1,625 cells per well (p = 0.49). In homogenised samples, ATP levels were unchanged at 1,625 cells per well (p = 0.49) but were significantly reduced at 3,250 cells per well (p = 0.047). This inconsistency mirrors the CellTox Green results and further indicates that the lysis buffer was not a reliable control.

Comparisons between fixed and lysis buffer–treated samples showed generally lower ATP levels in fixed samples. In non-homogenised cultures, this difference was significant at 1,625 cells per well (p = 0.0059) but not at 3,250 cells per well (p = 0.17). In homogenised samples, differences were significant at 3,250 cells per well (p = 0.032) but not at 1,625 cells per well (p = 0.17). The stronger and more consistent suppression of ATP following fixation confirms its suitability as a positive control. However, the detection of significant effects in homogenised samples again highlights the substantial contribution of bacterial ATP to the luminescence signal, reinforcing that CellTiter-Glo was not amoeba-specific under these non-axenic culture conditions. Together with the CellTox Green data above, these results demonstrate that bulk fluorescence- and luminescence-based readouts cannot reliably discriminate amoebal from bacterial activity in xenic *N. perurans* cultures, and that an amoeba-specific readout was required to derive robust drug-response conclusions.

### 3.4. Holographic microscopy

Holographic microscopy enabled label-free, amoeba-specific assessment of drug activity in *N. perurans* and directly addressed the bacterial interference limitations identified for the CellTox Green and CellTiter-Glo assays above. Because amoebae are several orders of magnitude larger than the co-cultured bacteria, size-based selection during single-cell tracking excluded any bacterial contribution to the readout. In addition, motility-based tracking measures a behaviour that is specific to viable amoeboid cells and cannot be produced by bacteria under the imaging conditions used, so drug effects reported here reflect activity on the amoeba itself rather than on the associated microbial community. Motile trophozoites displayed extended pseudopodia and high migration directness, whereas pseudocysts formed under suboptimal conditions and dead amoebae were rounded and immobile (Figure 9). Migration directness, expressed as the ratio of migration distance to motility (range 0 to 1), reduced background noise from instrument movement and provided a robust metric of drug response. Drug responses were monitored over 72 h with analysis windows at 0, 24, 48, and 72 h, enabling the timing of treatment effects to be resolved. Recovery assays, in which drugs were removed after 24 h, further differentiated amoebicidal activity (no recovery of motility) from amoebastatic effects (motility restored). Links to time-lapse videos illustrating amoeba behaviour under control and treatment conditions are provided in **Table S5**, and representative trajectories are shown in **S3**. This assay was applied to six drug classes: alkylphosphocholine (miltefosine), phenathridine (isometamidium), benzoxaboroles (AN11738), diamidines (pentamidine, DB75, and diminazene), nitroheterocyclics (nifurtimox and benznidazole), and suramin.

**Figure 9.**
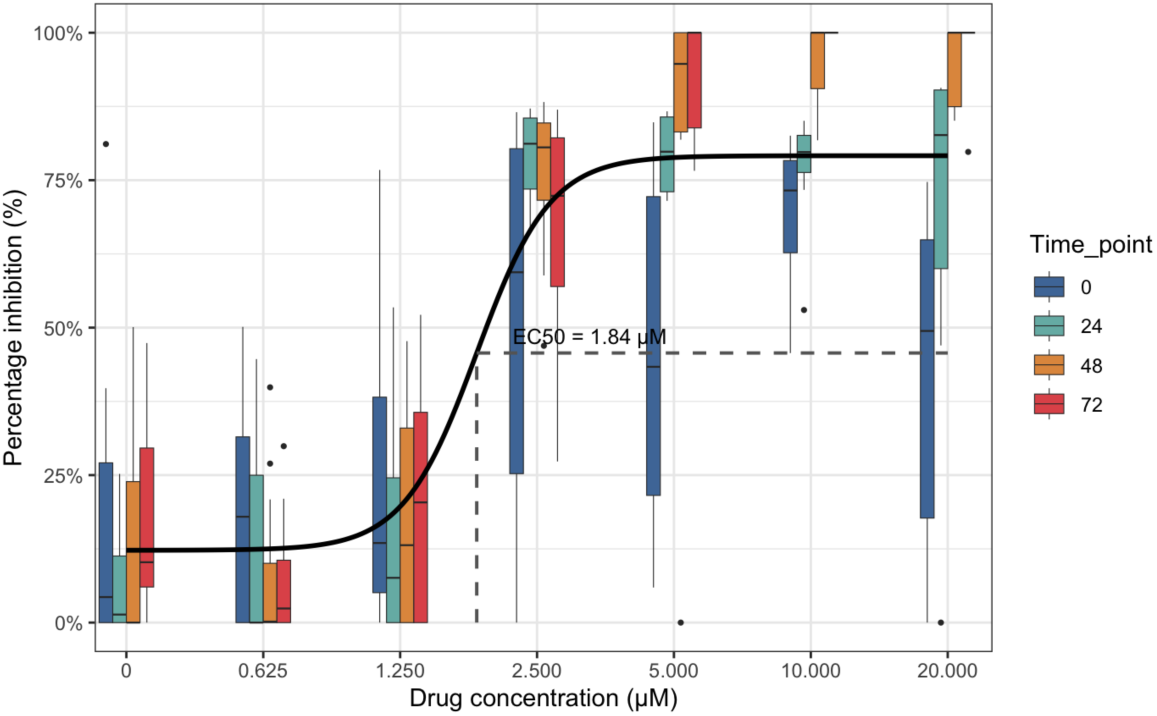
Half maximal effective concentration (EC_50_) of miltefosine for inhibiting *Neoparamoeba perurans* migration directness. Drug responses were monitored over 72 h with 4-hour analysis windows at 0, 24, 48, and 72 h. The EC_50_ represents the concentration required to reduce amoeba migration directness by 50% compared with untreated controls across all time points analysed. Miltefosine concentrations ranged from 20 µM to 0.625 µM.

#### 3.4.1. Alkylphosphocholine – miltefosine

As shown in **Table S5**, miltefosine treatment caused *N. perurans* trophozoites to rapidly round into pseudocysts, followed by cell rupture. There was a strong interaction between drug concentration and time point (χ² = 63.7, df = 18, p < 0.001), showing that migration directness declined more strongly with increasing drug concentration and with longer exposure, with both factors reinforcing each other. The EC₅₀, defined as the concentration causing a 50% reduction in migration directness relative to the untreated control, was estimated at 1.84 µM across all time points (**Figure 10**). In the recovery assay, amoeba exposed to miltefosine at 0 and 2.5 µM regained motility within 24 h of drug removal, whereas those treated at 5 and 10 µM remained immotile for the full 72 h observation period (**Figure 11**). This concentration-dependent effect is illustrated in the recovery videos provided in **Table S5**. Statistical modelling using a zero-inflated Gamma distribution failed to converge at concentrations ≥5 µM due to the complete absence of recovery. Therefore, this suggested miltefosine begins to exert a sustained amoebicidal effect at concentrations between 2.5 and 5 µM.

**Figure 10.**
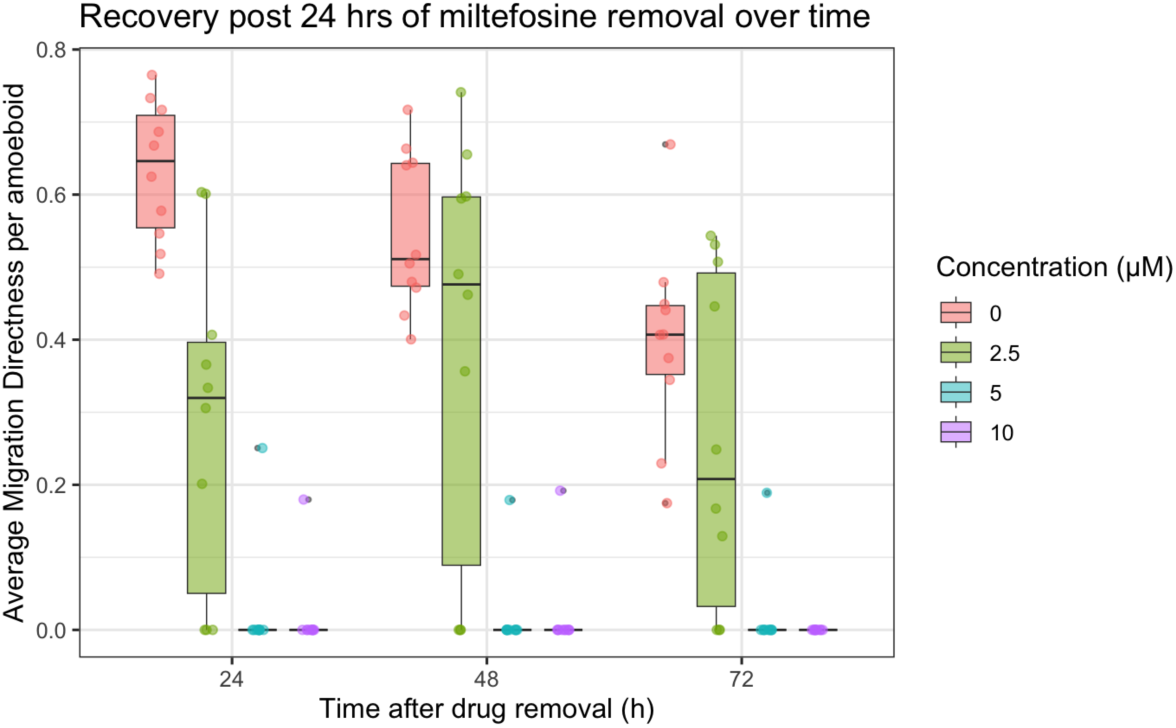
Recovery of amoeboid migration directness following miltefosine removal. Each dot represents a single amoeba tracked for 4 h (n = 10 amoebae per condition, measured in duplicates). Boxes show the distribution of average migration directness (median, interquartile range, and whiskers) for each drug concentration (coloured) at each time point after drug removal (24, 48, and 72 h). The y-axis shows raw average migration directness per cell.

**Figure 11.**
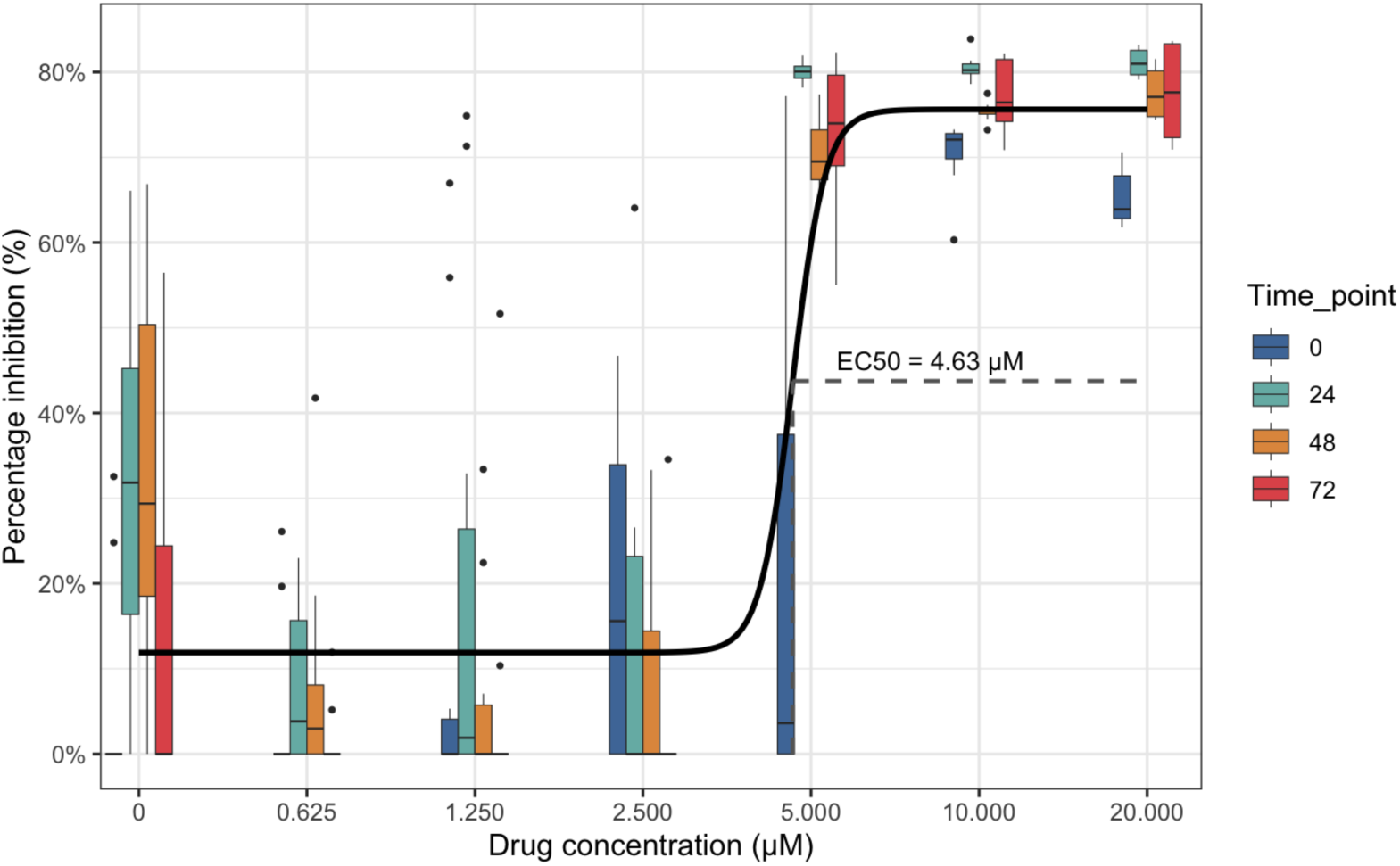
Half maximal effective concentration (EC_50_) of isometamidium for inhibiting *N. perurans* migration directness. Drug responses were monitored over 72 h with 4-hour analysis windows at 0, 24, 48, and 72 h. The EC_50_ represents the concentration required to reduce amoeba migration directness by 50% compared with untreated controls across all time points analysed. Isometamidium concentrations ranged from 20 µM to 0.625 µM.

#### 3.4.2. Phenathridine – isometamidium

Video capture (Table S5) showed isometamidium treatment caused *N. perurans* trophozoites to rapidly round into pseudocysts and become immotile, but without the cell rupture observed under miltefosine. Quantitative analysis of migration directness revealed a strong interaction between drug concentration and time point (LRT: χ² = 132.4, df = 18, p < 0.001), indicating that both higher concentrations and longer exposure progressively reduced motility. The EC₅₀ across all time points was estimated at 4.63 µM (**Figure 12**). Recovery assays showed that amoebae regained motility after 48 h following drug removal, consistent with an amoebastatic effect on *N. perurans.* In the recovery assay, isometamidium-treated amoeba initially lost motility but regained movement within 24 h of drug removal. Statistical analysis revealed a significant interaction between drug concentration and time point (χ² = 13.3, df = 6, p = 0.039), indicating that recovery depended on both factors. Amoebae exposed to lower concentrations recovered more quickly, whereas those treated at higher concentrations showed delayed restoration of motility (Table S5). These findings confirm that isometamidium caused a transient, time-dependent suppression of motility, consistent with an amoebostatic rather than a sustained amoebicidal effect.

**Figure 12.**
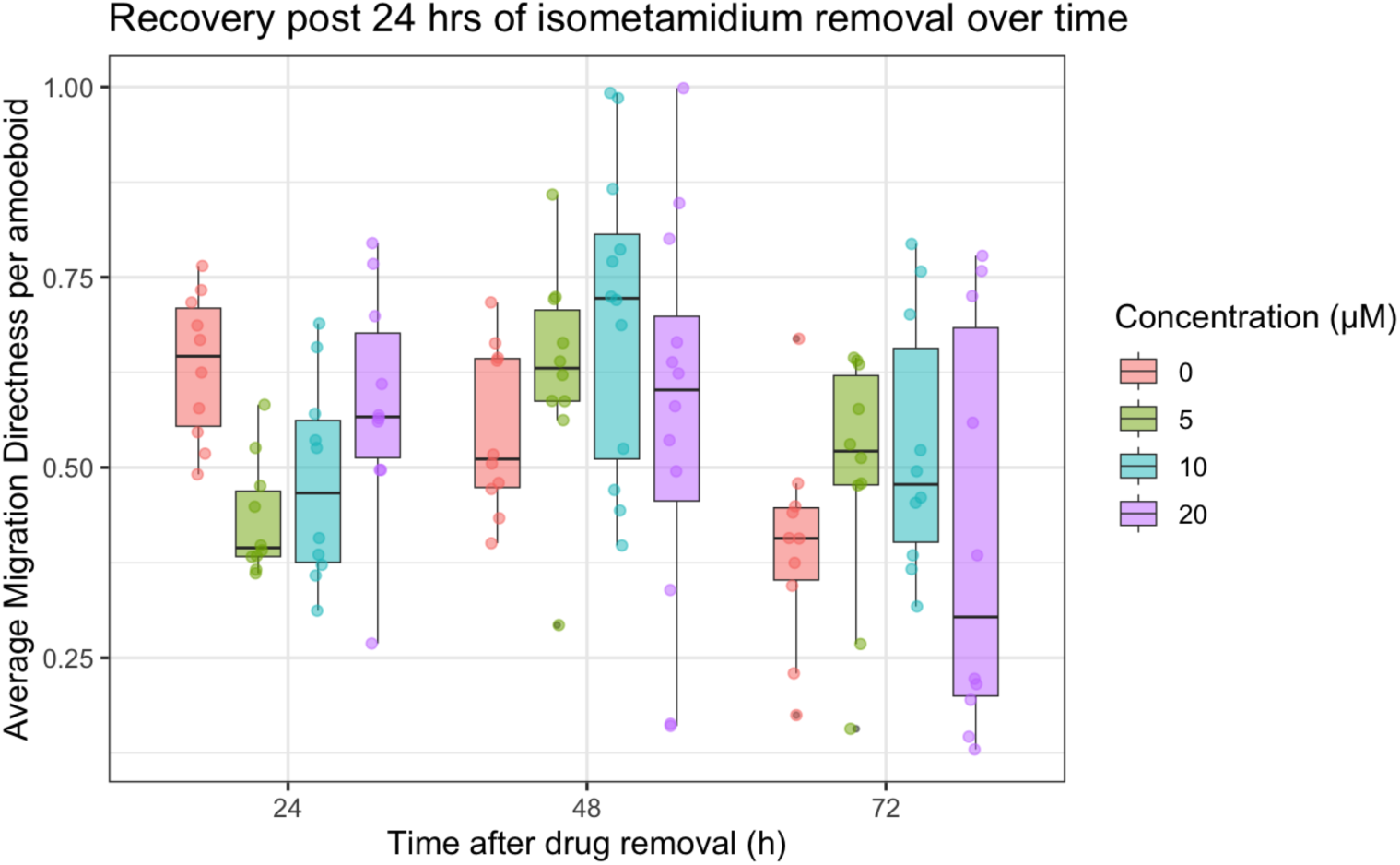
Recovery of amoeboid migration directness following isometamidium removal. Each dot represents a single amoeba tracked for 4 h (n = 10 amoebae per drug concentration, measured in duplicates). Boxes show the distribution of average migration directness (median, interquartile range, and whiskers) for each drug concentration (coloured) at each time point after drug removal (24, 48, and 72 h). The y-axis shows raw average migration directness per cell.

#### 3.4.3. Benzoxaboroles – AN11736

Unlike miltefosine and isometamidium, AN11736 did not induce pseudocyst formation but instead slowed trophozoite motility (Table S5). Quantitative analysis of migration directness showed a strong interaction between drug concentration and time point (LRT: χ² = 51.2, df = 18, p < 0.001). Both increasing drug concentration and longer exposure times were associated with significant reductions in migration directness (all p < 0.001). Even at the lowest concentration tested (0.625 µM), migration directness decreased markedly compared with controls (β = –0.76 ± 0.10, p < 0.001), with reductions approaching ∼80–90% at higher concentrations and later time points. The EC₅₀ was estimated at 0.249 µM (Figure 13), although this EC₅₀ value against amoeba motility may underestimate potency since a full sigmoidal dose–response curve could not be established at the concentrations tested. As there are no observable amoebae deaths after AN11736 treatment, a recovery assay on this drug is not performed. Collectively, these results indicate that AN11736 exerted a potent, concentration- and time-dependent inhibitory effect on N. perurans motility, but without inducing cell death.

**Figure 13.**
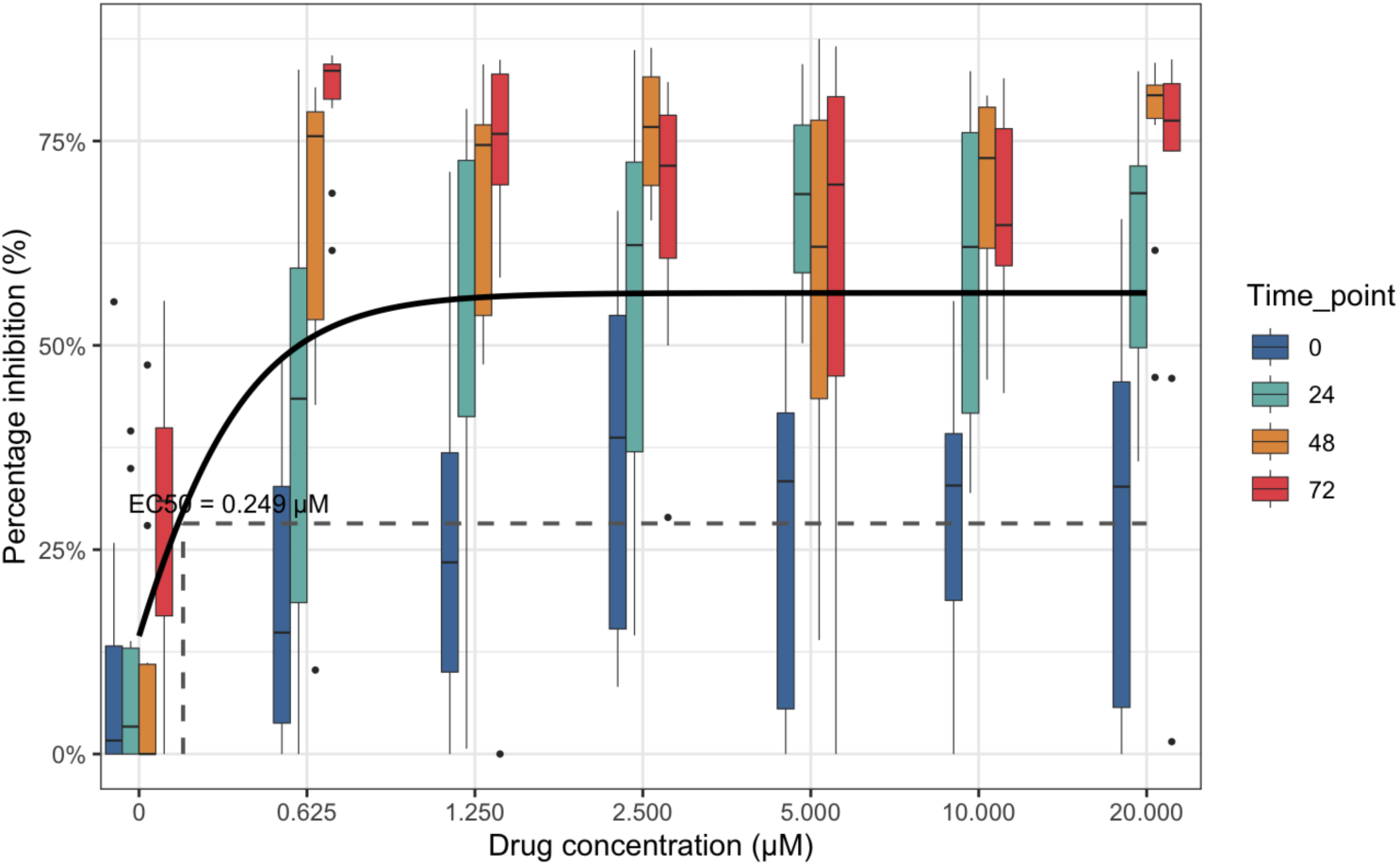
Half maximal effective concentration (EC_50_) of AN11736 for inhibiting *Neoparamoeba perurans* migration directness. Drug responses were monitored over 72 h with 4-hour analysis windows at 0, 24, 48, and 72 h. The EC_50_ represents the concentration required to reduce amoeba migration directness by 50% compared with untreated controls across all time points analysed. AN11736 concentrations ranged from 20 µM to 0.625 µM.

#### 4.3.4. Diamidines: pentamidines, DB75, and diminazene

Pentamidine did not induce visible changes in trophozoite morphology or motility (**Table S5**). Statistical analysis confirmed that migration directness varied significantly across time points (χ² = 16.2, df = 3, p = 0.001), but drug concentration had no detectable effect (χ² = 5.1, df = 6, p = 0.534), and no interaction between concentration and time was observed (χ² = 24.0, df = 18, p = 0.156). As drug concentration did not influence migration directness, an EC₅₀ could not be estimated, since the dose–response relationship was essentially flat rather than sigmoidal (**Figure 14**). As such, pentamidine treatment had no measurable impact on N. perurans motility.

**Figure 14.**
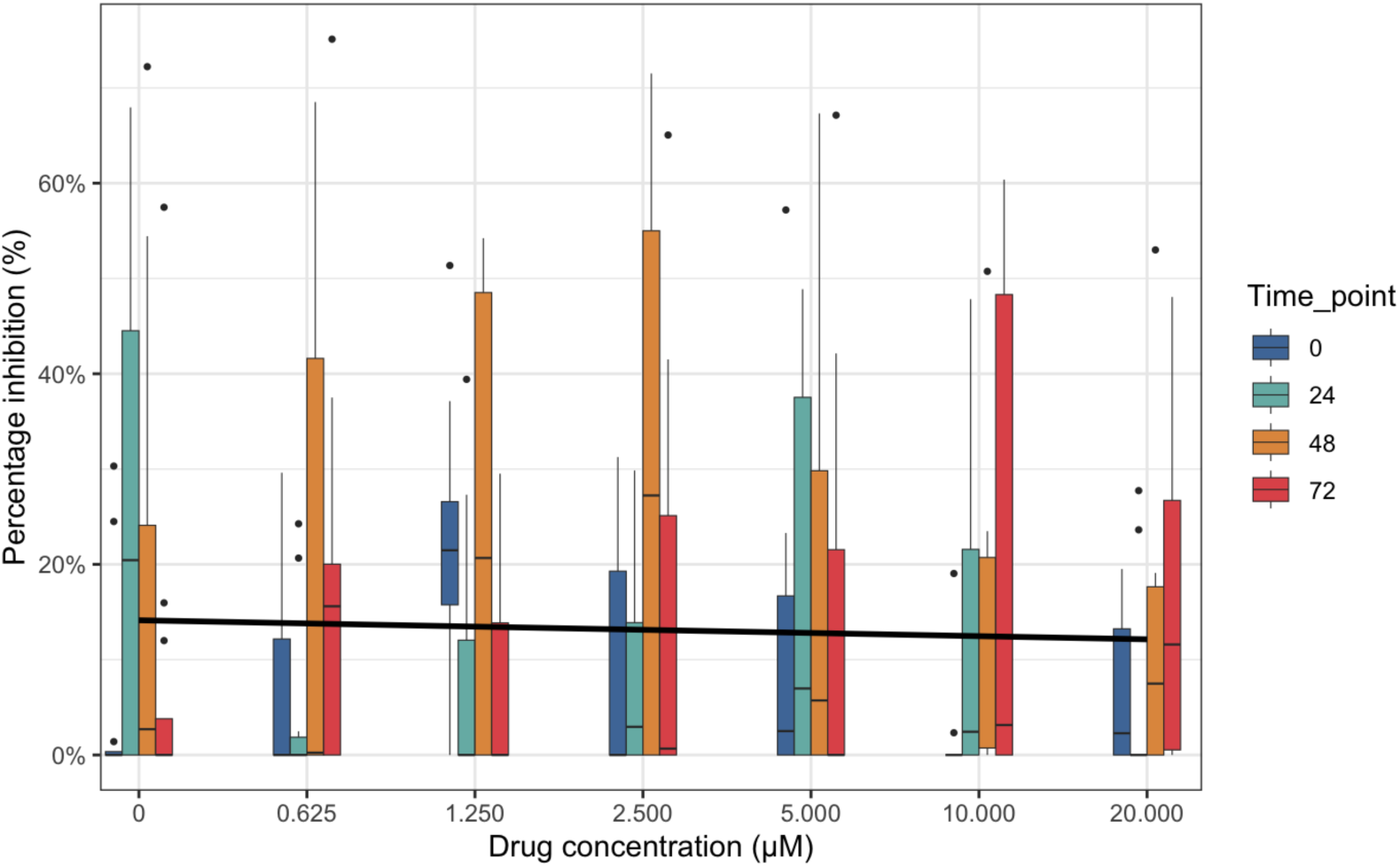
Half maximal effective concentration (EC50) of pentamidine for inhibiting *Neoparamoeba perurans* migration directness. Drug responses were monitored over 72 h with 4-hour analysis windows at 0, 24, 48, and 72 h. The EC50 represents the concentration required to reduce amoeba migration directness by 50% compared with untreated controls across all time points analysed. Pentamidine concentrations ranged from 20 µM to 0.625 µM.

DB75 did not induce visible changes in trophozoite morphology or motility (**Table S5**). Statistical analysis showed no evidence of an interaction between drug concentration and time point (χ² = 23.2, df = 18, p = 0.183). Furthermore, neither drug concentration (χ² = 8.3, df = 6, p = 0.215) nor time point (χ² = 5.5, df = 3, p = 0.136) significantly explained variation in migration directness compared with the null model. As no drug-induced effect was detected, an EC₅₀ could not be estimated. These results indicate that DB75 treatment had no measurable impact on N. perurans motility.

**Figure 15.**
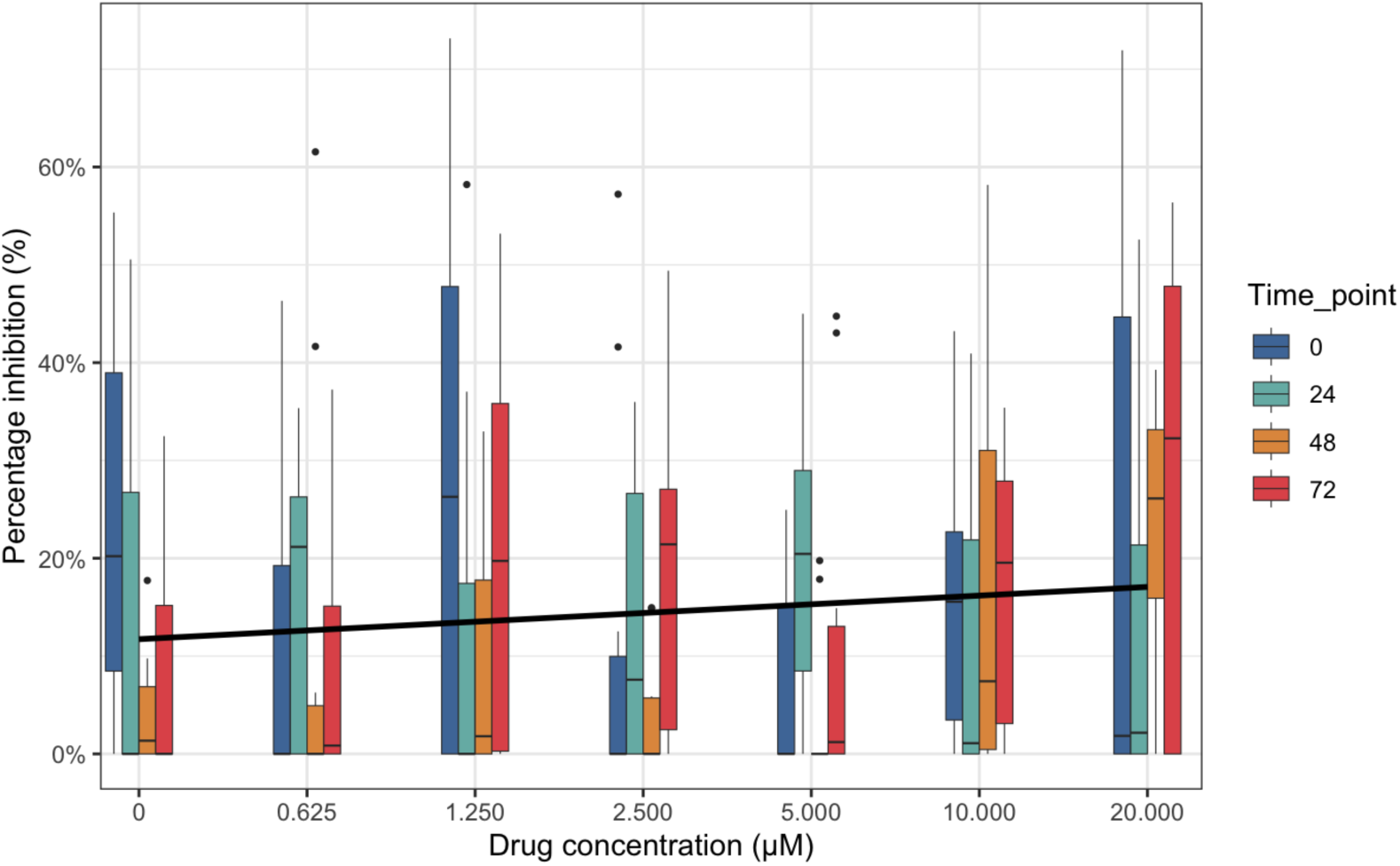
Half maximal effective concentration (EC_50_) of DB75 for inhibiting *Neoparamoeba perurans* migration directness. Drug responses were monitored over 72 h with 4-hour analysis windows at 0, 24, 48, and 72 h. The EC_50_ represents the concentration required to reduce amoeba migration directness by 50% compared with untreated controls across all time points analysed. DB75 concentrations ranged from 20 µM to 0.625 µM.

Diminazene did not induce visible changes in trophozoite morphology or motility (**Table S5**). Statistical analysis found no evidence of an interaction between drug concentration and time point (χ² = 20.8, df = 18, p = 0.291). Drug concentration alone showed a weak association with migration directness (χ² = 14.5, df = 6, p = 0.024), but this effect was not retained when time point was included in the model (χ² = 1.0, df = 3, p = 0.796). Similarly, time point alone had no effect relative to the null model (χ² = 0.79, df = 3, p = 0.852). As no consistent drug-induced effect was detected, an EC₅₀ could not be estimated, since the dose–response curve was essentially flat (Figure 16). These findings indicate that diminazene treatment had no measurable impact on N. perurans motility.

**Figure 16.**
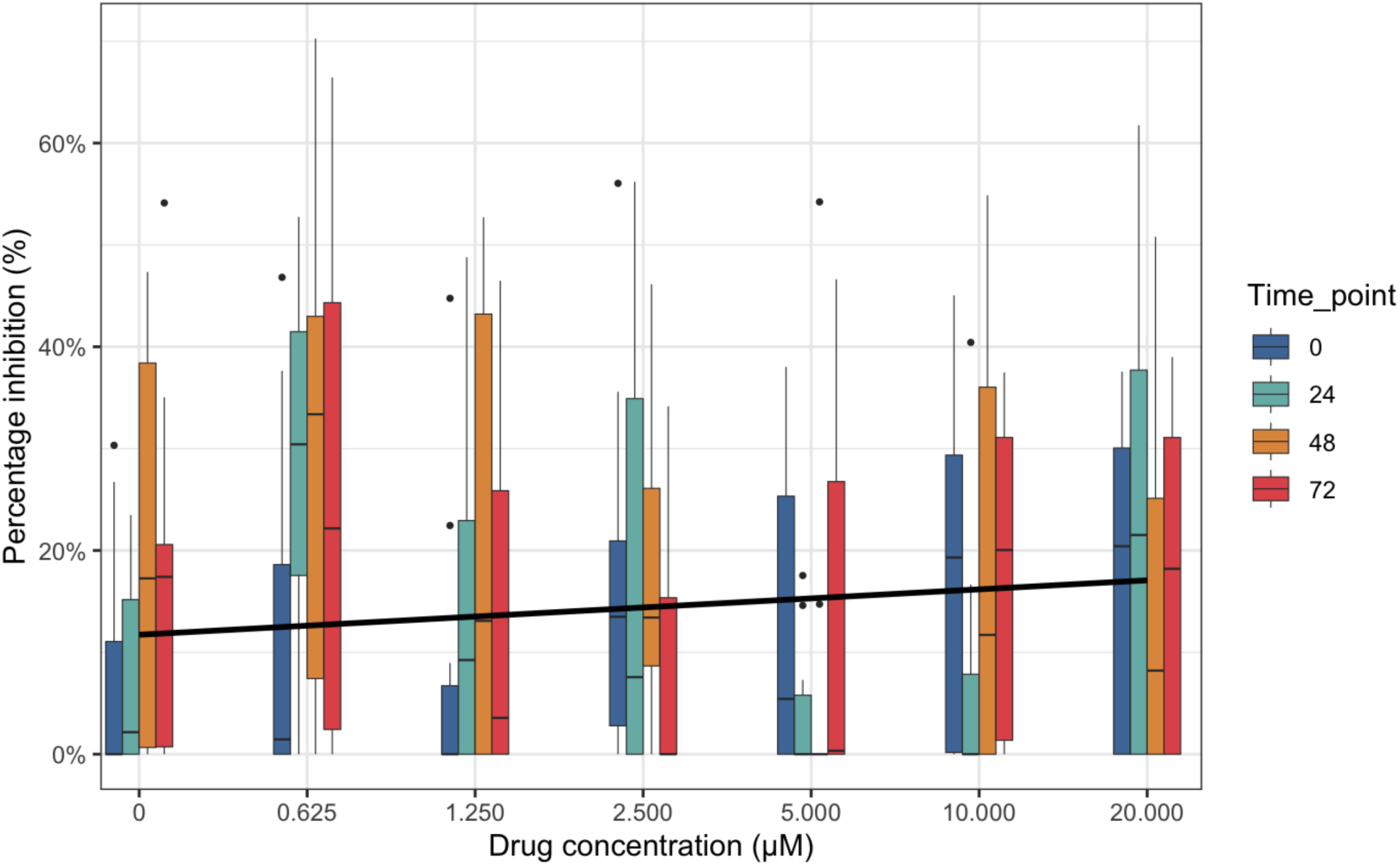
Half maximal effective concentration (EC_50_) of diminazene for inhibiting *Neoparamoeba perurans* migration directness. Drug responses were monitored over 72 h with 4-hour analysis windows at 0, 24, 48, and 72 h. The EC_50_ represents the concentration required to reduce amoeba migration directness by 50% compared with untreated controls across all time points analysed. Diminazene concentrations ranged from 20 µM to 0.625 µM.

#### 4.3.5. Nitroheterocyclics: Nifurtimox and Benznidazole

Nifurtimox did not induce visible changes in trophozoite morphology or motility (**Table S5**). Statistical analysis showed no evidence of an interaction between drug concentration and time point (χ² = 19.7, df = 18, p = 0.351). Neither drug concentration (χ² = 5.1, df = 6, p = 0.528) nor time point (χ² = 5.6, df = 3, p = 0.134) significantly explained variation in migration directness compared with the null model. As no drug-induced effect was detected, an EC₅₀ could not be estimated, since the dose–response curve was essentially flat rather than sigmoidal (Figure 17). These results indicate that nifurtimox treatment had no measurable impact on *N. perurans* motility.

**Figure 17.**
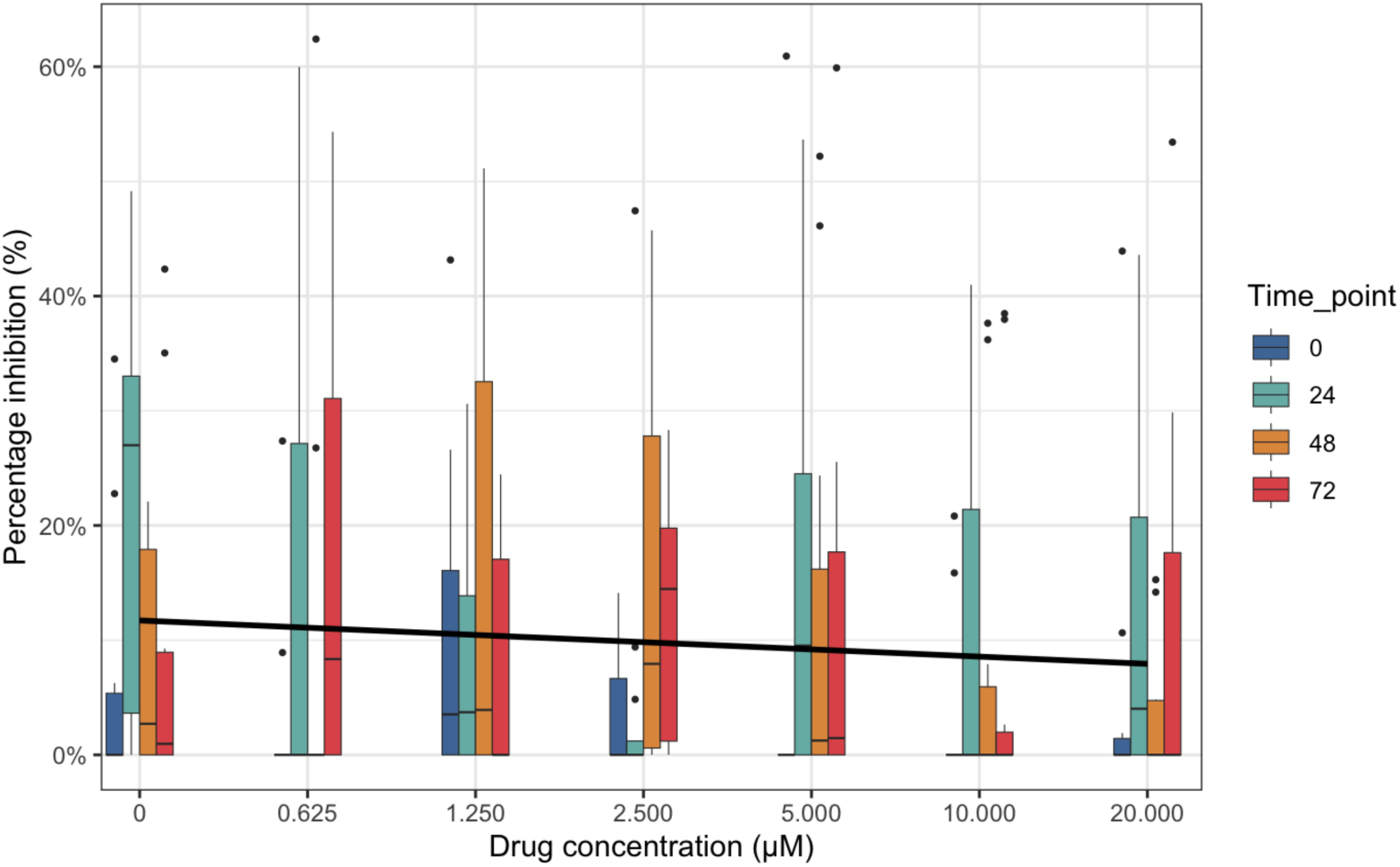
Half maximal effective concentration (EC_50_) of nifurtimox for inhibiting *Neoparamoeba perurans* migration directness. Drug responses were monitored over 72 h with 4-hour analysis windows at 0, 24, 48, and 72 h. The EC_50_ represents the concentration required to reduce amoeba migration directness by 50% compared with untreated controls across all time points analysed. Nifurtimox concentrations ranged from 20 µM to 0.625 µM.

Benznidazole did not induce visible changes in trophozoite morphology or motility (**Table S5**). Statistical analysis indicated no interaction between drug concentration and time point (χ² = 13.7, df = 18, p = 0.746). Time point was a significant predictor of migration directness (χ² = 34.1, df = 3, p < 0.001), whereas drug concentration had no effect (χ² = 6.2, df = 6, p = 0.399). As no drug-induced effect was detected, an EC₅₀ could not be estimated, since the dose–response relationship was essentially flat (Figure 18). These findings indicate that benznidazole treatment had no measurable impact on *N. perurans* motility.

**Figure 18.**
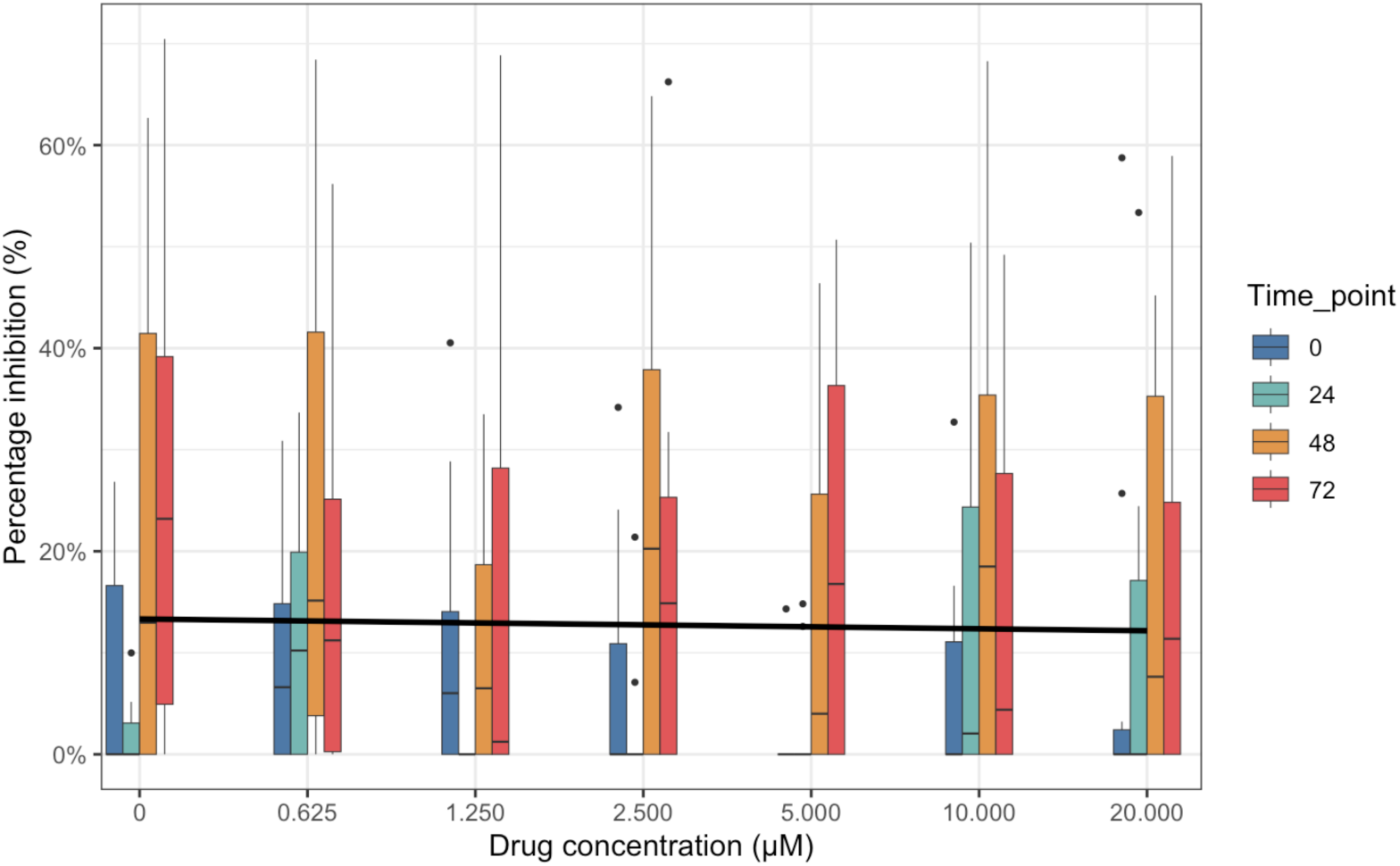
Half maximal effective concentration (EC_50_) of benznidazole for inhibiting *Neoparamoeba perurans* migration directness. Drug responses were monitored over 72 h with 4-hour analysis windows at 0, 24, 48, and 72 h. The EC_50_ represents the concentration required to reduce amoeba migration directness by 50% compared with untreated controls across all time points analysed. Benznidazole concentrations ranged from 20 µM to 0.625 µM.

Average migration directness was analysed using a Gamma GLM with log link. Time point had a significant effect (χ² = 34.1, df = 3, p < 0.001), with directness reduced at 48 h (β = –0.25 ± 0.06, p < 0.001) and 72 h (β = –0.23 ± 0.06, p < 0.001) compared with baseline. Drug concentration had no significant effect (all p > 0.07), and there was no evidence of a drug concentration and time interaction (p = 0.75).

#### 4.3.6. Suramin

Suramin did not induce visible changes in trophozoite morphology or motility (**Table S5**). Statistical analysis showed no evidence of an interaction between drug concentration and time point (χ² = 16.8, df = 18, p = 0.540). Time point was a significant predictor of migration directness (χ² = 9.4, df = 3, p = 0.024), and drug concentration contributed weakly when included alongside time (χ² = 9.2, df = 3, p = 0.027). However, drug concentration alone was not significant relative to the null model (χ² = 6.8, df = 6, p = 0.344). The dose–response relationship was essentially flat, and no EC₅₀ could be estimated (Figure 19). These results indicate that suramin treatment had no measurable effect on *N. perurans* motility.

**Figure 19.**
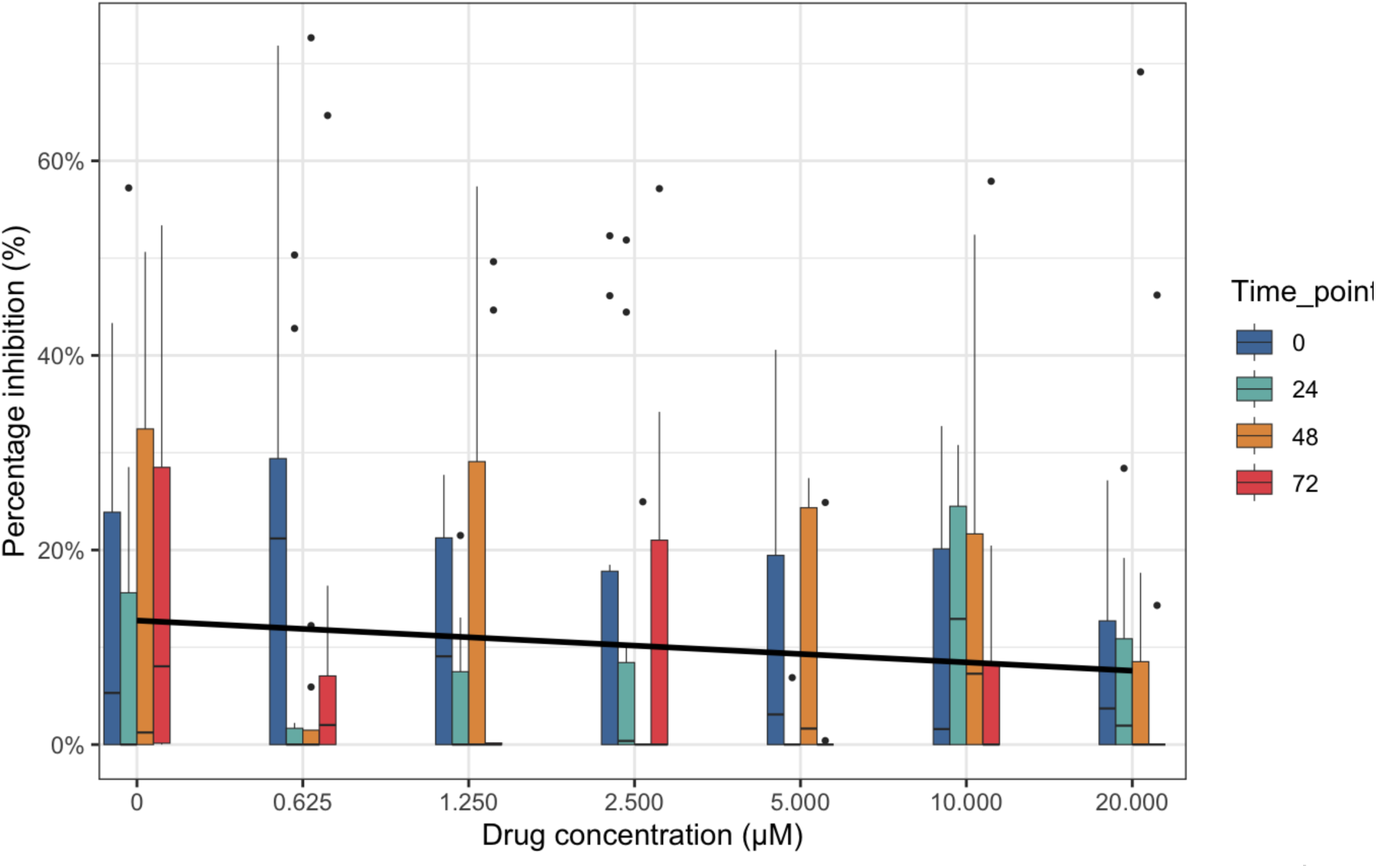
Half maximal effective concentration (EC_50_) of suramin for inhibiting *Neoparamoeba perurans* migration directness. Drug responses were monitored over 72 h with 4-hour analysis windows at 0, 24, 48, and 72 h. The EC_50_ represents the concentration required to reduce amoeba migration directness by 50% compared with untreated controls across all time points analysed. Suramin concentrations ranged from 20 µM to 0.625 µM.

### 4.4. In vivo drug tolerance assessment

Miltefosine, isometamidium, nifurtimox and benznidazole were selected to be carried forward to an *in vivo* tolerance assessment on Atlantic salmon (*Salmo salar*) to evaluate their safeness before deployment in further field or lab trials (Nifurtimox and benznidazole were chosen based on preliminary qualitative testing (i.e. via propidium iodide) before cell mortality assays had been developed). In this *in vivo* tolerance assessment, a total of 250 fish were intramuscularly (IM) injected with either control (PBS with 1% DMSO) or one of the four drugs on day 1, day 3, and day 5 (one injection on each dosing day). The binomial GLM indicated that mortality differed significantly among treatments (likelihood-ratio test, χ² = 14.79, df = 4, p = 0.030, p < 0.05) and across days (likelihood-ratio test, χ² = 71.80, df = 5, p = 4.32 × 10⁻¹⁴, p < 0.001). A stratified Mantel–Haenszel chi-squared test was also conducted to compare mortality between each treatment and the Control group, adjusting for day as a stratification factor. Benznidazole showed a statistically significant reduction in mortality compared to Control (χ² = 6.01, df = 1, p = 0.014; OR = 0.11, 95% CI ≈ 0.0137–0.90). No significant differences were detected for isometamidium (p = 0.57, OR = 0.72, 95% CI ≈ 0.24–2.22), Miltefosine (p = 0.97, OR = 0.98, 95% CI ≈ 0.35–2.72), or Nifurtimox (p = 0.22, OR = 1.80, 95% CI ≈ 0.70–4.60). These results corroborate the binomial GLM findings, indicating that benznidazole treatment was associated with significantly lower mortality relative to the Control group. Based on the binomial GLM, the majority of deaths occurred on day 2, which also coincided with the day of the most intense handling stress for the fish such as tank transfer and IM injection (Figure 20). Mortality varied significantly by day, with the highest mortality observed on day 2, coinciding with the period of greatest handling stress. Post-hoc Tukey comparisons indicated higher mortality on day 2 than on day 5 (p = 0.010, OR = 6.0, 95% CI ≈ 2.0–18.0) and day 6 (p = 0.033, OR = 4.0, 95% CI ≈ 1.3–12.0). No other day-to-day differences were significant (p > 0.05).

**Figure 20.**
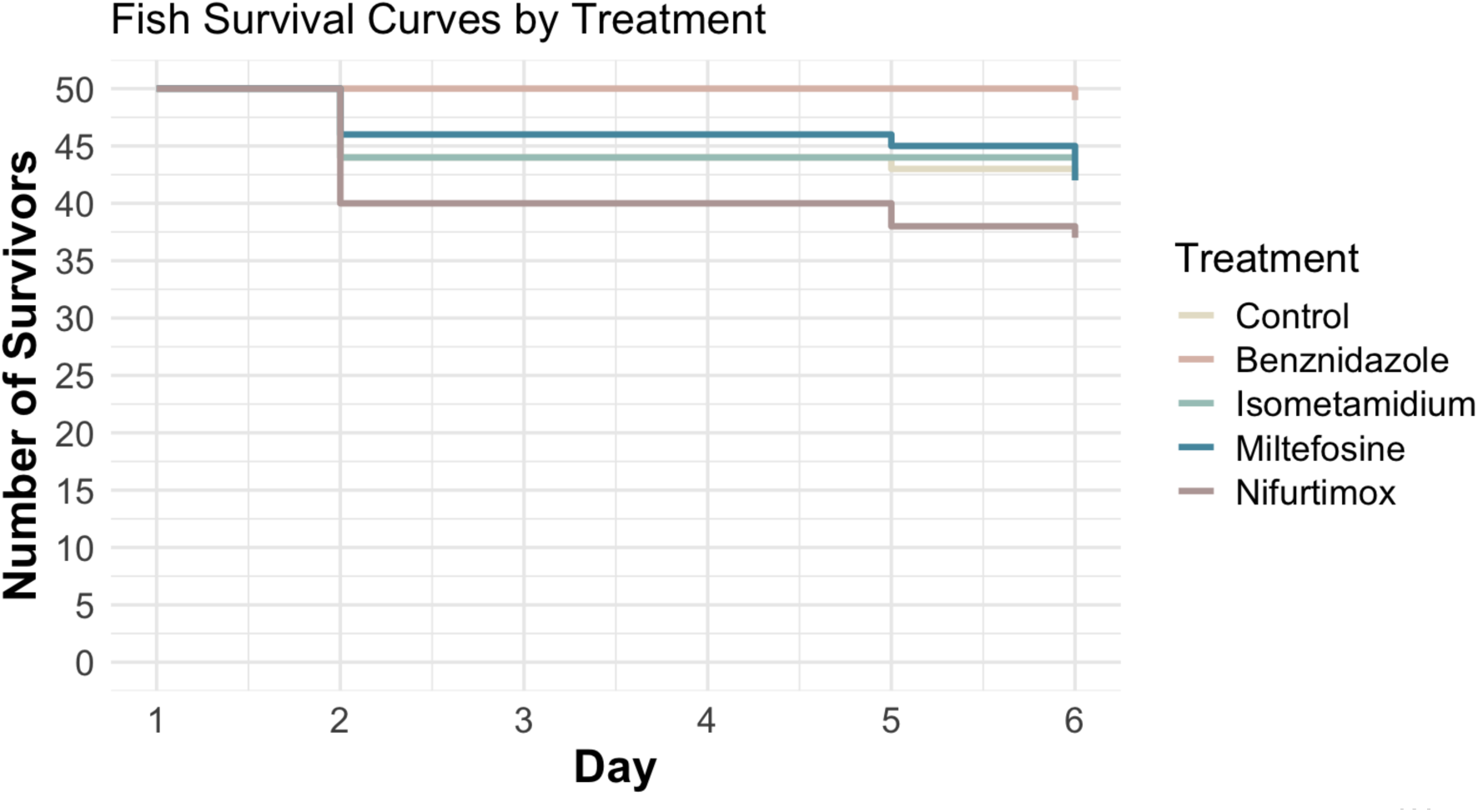
Survival of Atlantic salmon (*Salmo salar*) during a six day tolerance trial following intramuscular (IM) dosing of candidate drugs. Step curves show the number of surviving fish per treatment from Day 1 to Day 6 (n = 50 per treatment at Day 1). Fish received IM injections on Days 1, 3, and 5; the control group received vehicle (PBS with 1% DMSO). Survival was recorded daily with initial 50 minus cumulative mortalities.

To quantify the stress response following IM drug injection, serum cortisol (ng/mL) was measured from 3 fish per treatment group on alternate days. Cortisol concentrations did not differ among treatments (likelihood-ratio test: χ² = 2.70, df = 4, p = 0.609) and did not vary across sampling days (likelihood-ratio test: χ² = 0.703, df = 1, p = 0.402) (Figure 21). A global comparison of the model, including treatment and day against an intercept-only model was also non-significant (χ² = 3.12, df = 5, p = 0.681). Thus, the drug treatments did not significantly elevate cortisol levels.

**Figure 21.**
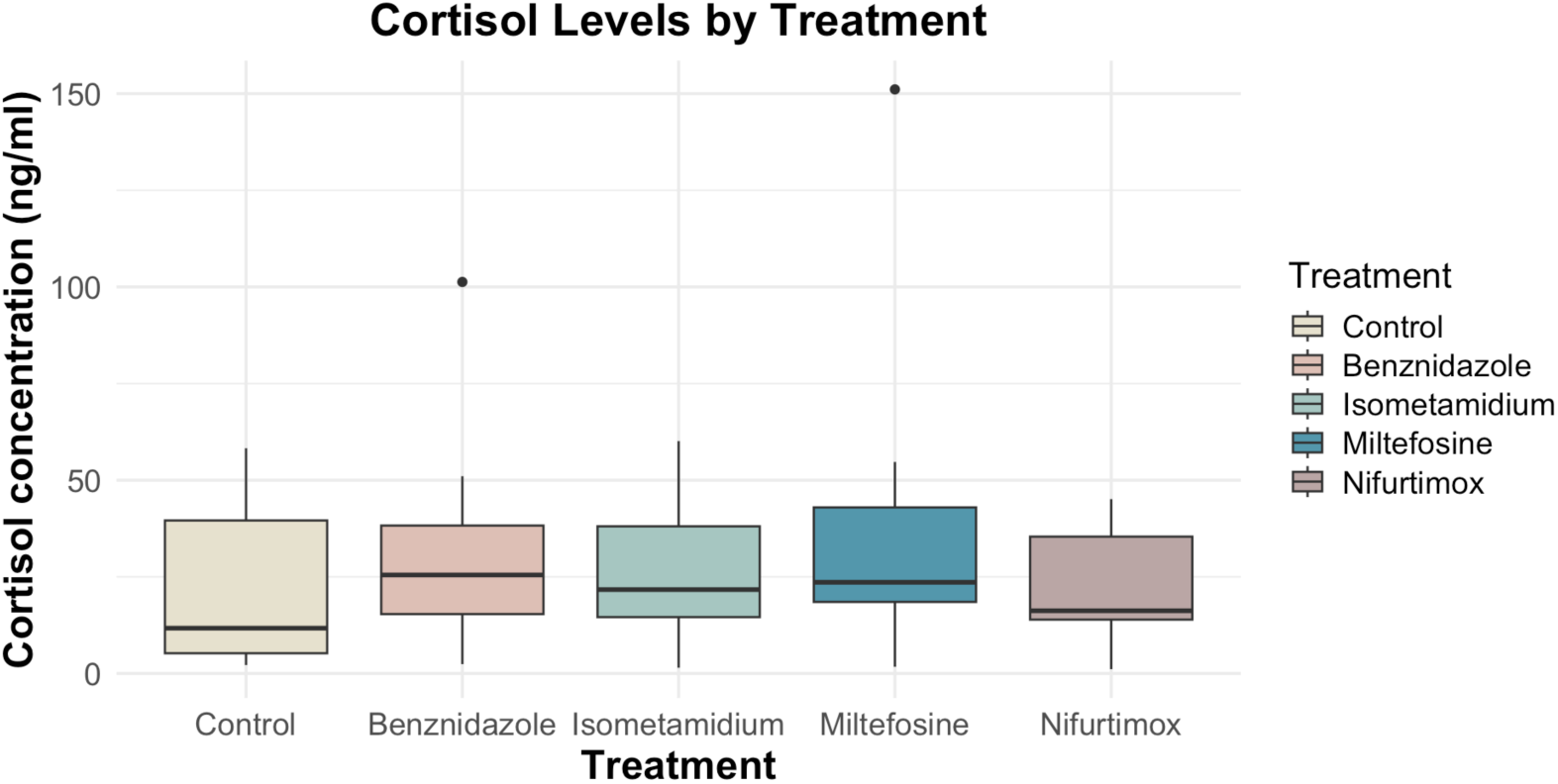
Serum cortisol in Atlantic salmon (*Salmo salar*) following intramuscular administration of candidate drugs. Blood was sampled from three fish per treatment every other day over a six-day tolerance trial (Days 1, 3, and 5), coinciding with dosing. Data from all sampling days were pooled; each boxplot represents the overall cortisol distribution for a treatment across the trial. The y-axis shows cortisol concentration (ng/ml). The x-axis shows treatment groups.

### 4.5. *In vivo* drug trial on efficacy against AGD

Benznidazole, isometamidium, and miltefosine were carried forward to the in vivo drug trial to determine their efficacy against amoebic gill disease (AGD). Nifurtomox was excluded based on high mortality in the tolerance trial. In this in vivo drug trial, a total of 1,231 fish, assigned to different treatments, were harvested and assessed for gill score using a 0 to 5 scale (0 = no lesions, 5 = severe lesions). To investigate the association between gill score and treatment, while accounting for random factors (sentinel pen), a proportional odds logistic regression model was fitted to the gill scores of all AGD fish. A significant association was found between improved gill score and the treatment received by the AGD fish (p = 0.02104, p < 0.05). As shown in **Figure 22**, significantly lower gill scores were observed when comparing benznidazole to no treatment control (p = 0.023, p < 0.05, OR = 0.64, 95% CI ≈ 0.44–0.94), isometamidium to no treatment control (p = 0.0092, p < 0.01, OR = 0.61, 95% CI ≈ 0.42–0.88), and miltefosine to no treatment control (p = 0.0092, p < 0.01, OR = 0.62, 95% CI ≈ 0.43–0.89). These data suggests a significant improvement in gill score in AGD infected fish for all treatments deployed, with the largest average improvement seen for miltefosine.

**Figure 22.**
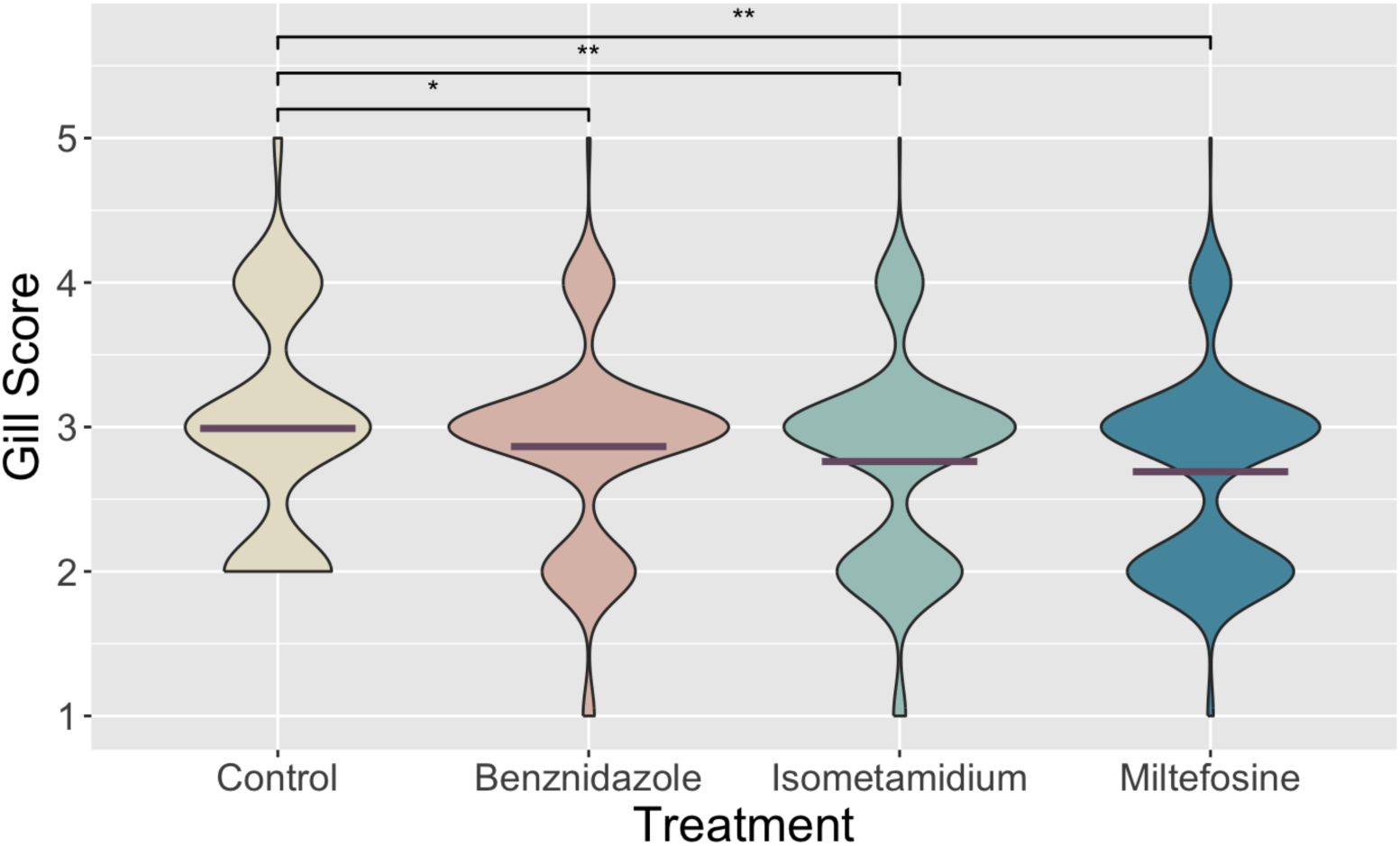
The distribution of gill scores in Atlantic salmon (*Salmo salar*) (n=1,231) following a two-week in vivo drug trial for amoebic gill disease. The X-axis represents the treatment group each fish received: benznidazole (n=310), isometamidium (n=310), miltefosine (n=307), or control (n=304). The Y-axis represents the gill score assessed using a standardised 0 to 5 scale (0 = no lesions, 5 = severe lesions). Statistical significance thresholds are represented using asterisks: one (*p ≤ 0.05), two (**p ≤ 0.01), and three (***p ≤ 0.001).

From the 1,231 AGD positive fish that received different treatments, a sub-sample of 396 gill swab samples (no treatment control, n = 112; benznidazole, n = 96; isometamidium, n = 96; miltefosine, n = 92) were analysed using qPCR to quantify the *N. perurans* load in nanogram (ng) of DNA. The linear regression model fitted to the log-transformed *N. perurans* DNA data confirmed a statistically significant positive association between gill score and amoebic load (F(4,391) = 6.36, p = 5.80 × 10⁻⁵, p<0.001, R² = 0.06) (**Figure 23**). Relative to gill score 1, mean DNA quantities increased progressively with higher gill scores, ranging from approximately 1.6 fold higher at gill score 2 to over 3-fold higher at gill score 5, indicating a clear monotonic relationship between disease severity and parasite load. Next, a linear mixed-effects model was fitted to the log-transformed DNA nanogram qPCR data, incorporating random factors for sentinel pen and qPCR batch ID to account for clustering and run-to-run variation. This analysis again revealed a significant association between treatment and *N. perurans* DNA load (p = 0.04534, p < 0.05) but, intriguingly, in the opposite direction to the AGD gill score, with more amoeba DNA measured in some treatments than the control. As such, the amoebic load, quantified by qPCR as shown in Figure 24, was significantly higher in the benznidazole group than in the no treatment control (p = 0.026, p < 0.05, OR = 1.87, 95% CI≈ 1.079 – 3.27) and also higher in the isometamidium group (p = 0.0076, p < 0.01, OR = 2.10, 95% CI ≈ 1.22 – 3.61). However, there was no significant difference between miltefosine and the control (p = 0.89, p >0.05, OR = 0.96, 95% CI ≈ 0.56– 1.67) (Figure 24). The decoupling of gill scores and qPCR represents a challenge to interpret therapeutic effect, and we reflect on this further in the Discussion.

**Figure 23.**
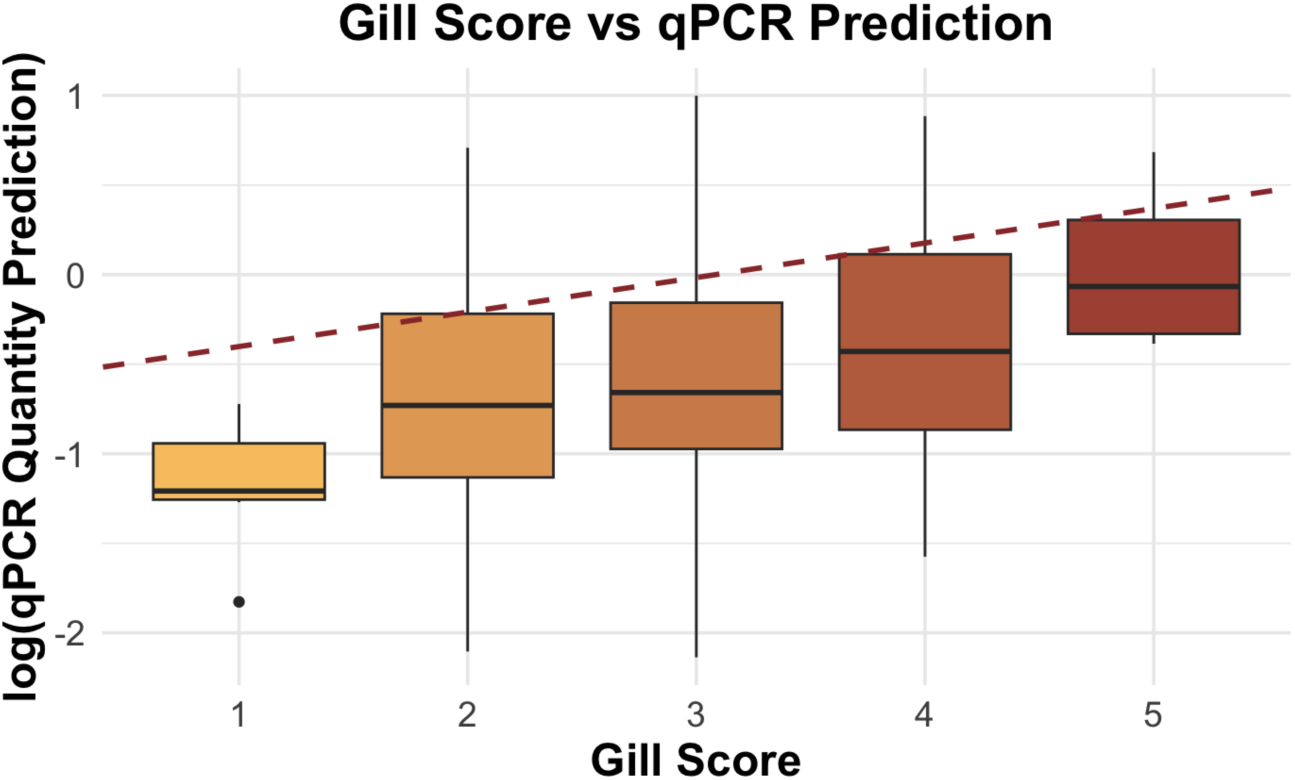
The association between the log-transformed *N. perurans* DNA load (nanograms) and gill score in Atlantic salmon (*Salmo salar*) following an in vivo drug trial for amoebic gill disease (AGD). The X-axis represents the gill score assessed using a standardised 0-to-5 scale (0 = no lesions, 5 = severe lesions). The Y-axis represents the log-transformed *N. perurans* DNA load quantified by qPCR from gill swabs. The brown dash line represents the linear regression model fitted to the data, indicating the predicted trend between these two variables.

**Figure 24.**
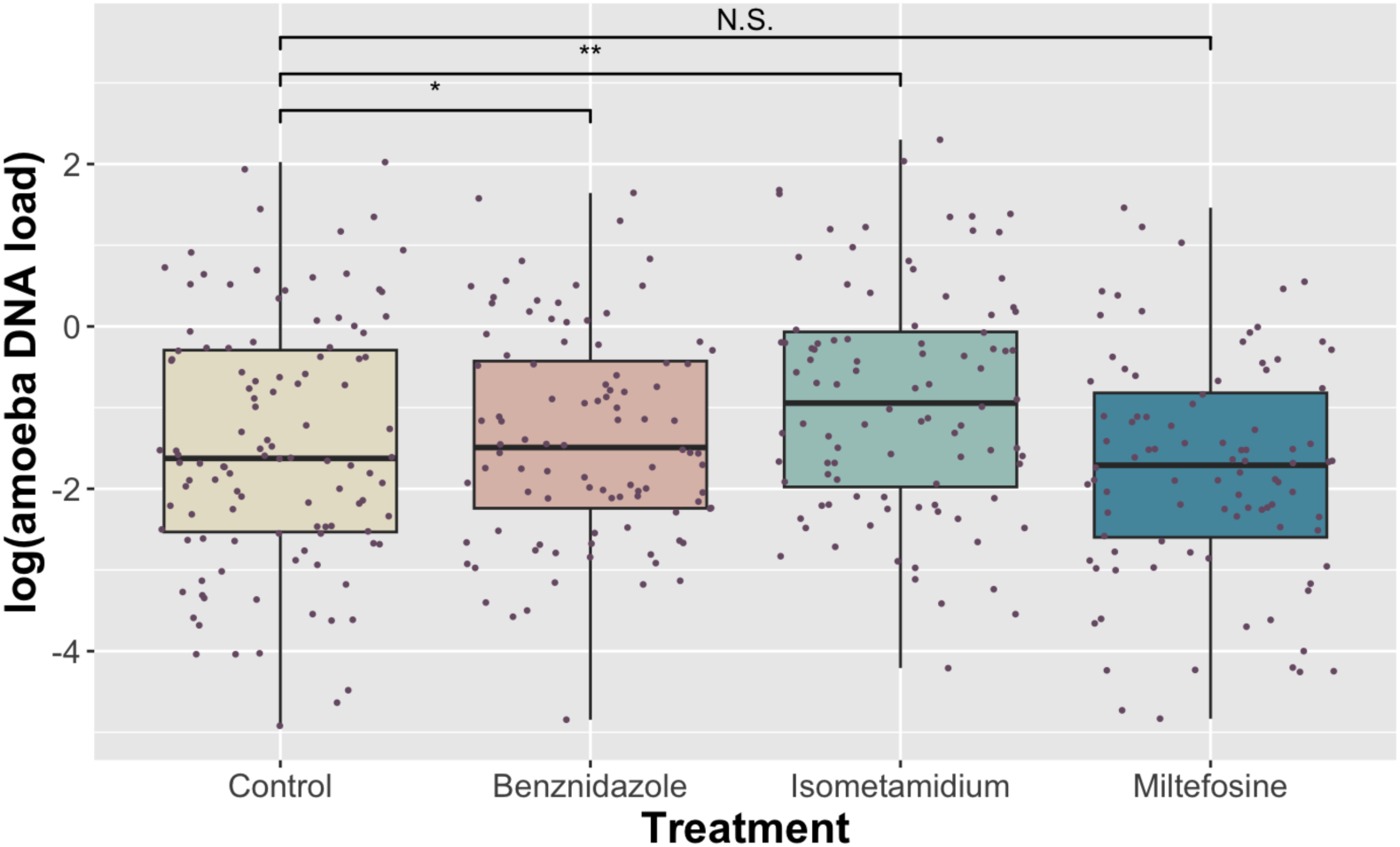
The log-transformed *N. perurans* DNA load (nanograms) in Atlantic salmon (*Salmo salar*) following an in vivo drug trial to evaluate treatment efficacy against amoebic gill disease (AGD). The X-axis represents the treatment group each fish received: Benznidazole, Isometamidium, Miltefosine, or Control. The Y-axis represents the log-transformed *N. perurans* DNA load quantified by qPCR from gill swab samples. Each dot represents the average of technical triplicate of amoeba DNA load after qPCR perform for each gill swab sample. Statistical significance thresholds are represented using asterisks: one (*p ≤ 0.05), two (**p ≤ 0.01), and three (***p ≤ 0.001); N.S. indicates no significant difference.

Of the 1374 fish with weight (g) and height (cm) recorded, only a subset of fish (n = 239) has fish IDs allowing linkage to their corresponding gill scores. Within this subset, a simple linear regression model on log-transformed condition factor (log K) (Equation 5) indicated a weak negative association with gill score (F(4,234) = 3.74, p = 0.0057, R² = 0.06) (Figure 25). Additionally, within this subset of fish (n = 239), to account for treatment and random effect (sentinel pen), an additive linear mixed-effects model was also fitted (Equation 6). An interaction linear mixed-effects model including the interaction term for treatment and gill score (Equation 7) improved overall model fit (LRT χ²(12) = 43.73, p < 0.001), suggesting possible treatment-specific variation in the gill score to K relationship. Previous analyses indicated that treatment significantly improved gill condition, with lower gill scores observed in treated groups. Furthermore, the interaction model provided a significantly better fit than models containing gill score alone (LRT χ²(11) = 43.43, p = 9.14 × 10⁻⁶) or treatment alone (LRT χ²(12) = 43.73, p = 1.70 × 10⁻⁵). However, the interaction model was rank-deficient because several treatment-gill score combinations were empty, and the individual treatment and gill score coefficients were not significant (all p > 0.05). In this subset, there were no control fish with gill score 1 and no fish with gill score 5 in any treatment group. The additive model (Equation 6) was therefore retained as the simpler model, with gill score acting as a supportive rather than a decisive predictor of condition factor.

**Figure 25.**
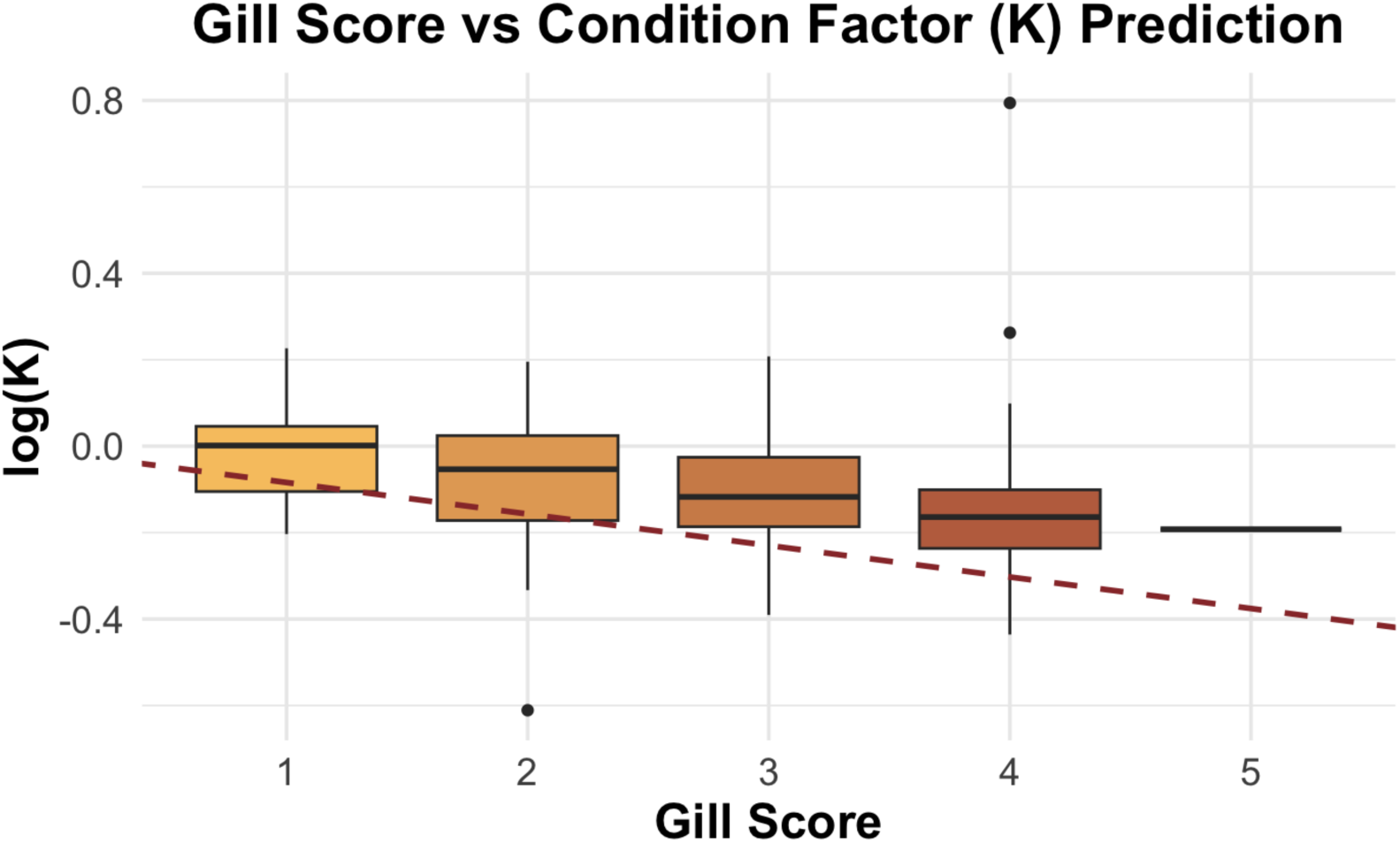
The association between the log-transformed condition factor (K) and gill score in Atlantic salmon (*Salmo salar*) following an *in vivo* drug trial for amoebic gill disease (AGD). The X-axis represents the gill score assessed using a standardised 0-to-5 scale (0 = no lesions, 5 = severe lesions). The Y-axis represents the log-transformed N. perurans DNA load quantified by qPCR from gill swabs. The brown dashed line represents the linear regression model fitted to the data, indicating the predicted trend between these two variables.

Condition factor (K) was analysed for all 1374 fish with weight (g) and height (cm) recorded. As before, gill scores were unavailable for most fish in this cohort and were therefore not included. A linear mixed-effects model (Equation 8) was fitted to the log-transformed condition factor data including cage as a random effect to account for between sentinel pen variation. No significant differences in condition factor were detected between treatment groups (Figure 26).

**Figure 26.**
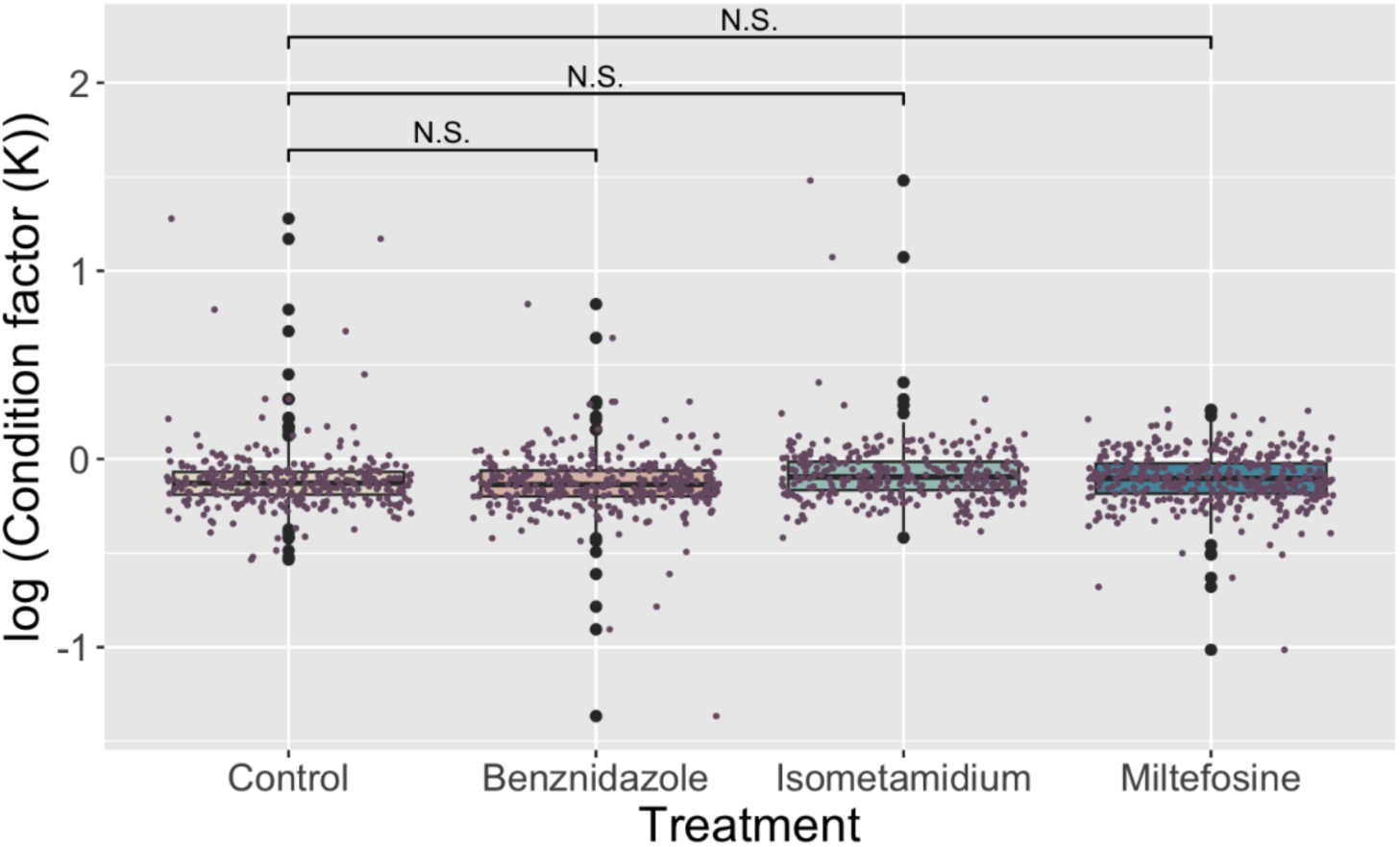
The log-transformed condition factor in Atlantic salmon (Salmo salar) following an in vivo drug trial to evaluate treatment efficacy against amoebic gill disease (AGD). The X-axis represents the treatment group each fish received: Benznidazole, Isometamidium, Miltefosine, or Control. The Y-axis represents the log-transformed condition factor calculated by Equation 1. Each dot denotes the condition factor of each fish. Statistical significance thresholds are represented using asterisks: one (*p ≤ 0.05), two (**p ≤ 0.01), and three (***p ≤ 0.001); N.S. indicates no significant difference.

## 4. Discussion

This study represents the first practical evaluation of repurposed trypanocidal drugs for amoebic gill disease (AGD). We (i) benchmarked three *in vitro* screening approaches under the constraints imposed by *Neoparamoeba perurans* xenic culture, (ii) identified a small subset of trypanocidal candidates with measurable activity in vitro, and (iii) translated leading candidates into in vivo tolerance and field efficacy trials under natural AGD exposure using clinically relevant endpoints (gill score) alongside molecular quantification (qPCR). Collectively, these results support the feasibility of cross-disciplinary drug repurposing for AGD, while also clarifying the methodological and biological barriers that must be addressed before such compounds can be credibly prioritised for development [1], [7], [8], [9].

A core challenge throughout was that xenic *N. perurans* culture undermines assay specificity and reproducibility. Nucleic-acid dyes such as Propidium iodide (PI) can generate misleading signals when extracellular DNA is abundant in EPS-rich biofilms, producing PI-positive background that can mask viable amoebae and inflate apparent death **[47], [48], [49]**. This is important because biofilms co-occur with ***N. perurans*** under farm conditions and can carry ***N. perurans*** DNA during outbreaks **[50]**. Likewise, bacterial co-culture, especially stable associations dominated by ***Vibrio spp***., complicates the interpretation of viability assays that are not amoeba-specific **[14]**. In this context, we found major limitations in plate-reader multiplex workflows based on membrane integrity dyes (CellTox Green) and ATP readouts (CellTiter-Glo®): both measures were strongly influenced by bacterial/biofilm signal and could not reliably discriminate bacterial-only controls from xenic amoeba cultures, even after attempted baseline subtraction **[41]**. This is not simply a technical inconvenience but a validity issue: assays validated using broad-spectrum disinfectants (e.g., freshwater, hydrogen peroxide, chloramine-T) will inevitably “work” in mixed systems because they act on bacteria as well as amoebae, but they do not establish whether a candidate drug is acting on *N. perurans* specifically **[41]**. By contrast, digital holographic microscopy provided the most amoeba-informative readout because amoebae are several orders of magnitude larger than the co-cultured bacteria, allowing size-based selection during single-cell tracking to exclude any bacterial contribution to the readout. Trophozoites and pseudocysts could be recognised on the basis of morphology, and motility-based tracking measures a behaviour that is specific to viable amoeboid cells and cannot be produced by bacteria under the imaging conditions used **[42], [43], [44]**. Drug effects reported here therefore reflect activity on the amoeba itself rather than on the associated microbial community, without fluorescence or ATP interference. The trade-off such method was analytical burden: amoebal polymorphism and stage transitions caused tracking fragmentation and identity switching that required manual curation. A clear next step is to reduce this bottleneck using machine-learning object detection and stage classification approaches (e.g., YOLO-family frameworks), as demonstrated for trophozoite/cyst recognition in other amoebae, which should enable scalable, standardised phenotyping across plates and timepoints **[51], [52]**.

We therefore restrict our mechanistic and translational interpretation of in vitro drug activity to the holographic findings and regard the quantitative fluorescence and luminescence data primarily as evidence of the methodological constraints of xenic culture rather than as independent confirmation of drug efficacy. Using this assay, only a subset of trypanocidal candidates produced meaningful phenotypes *in vitro*. Miltefosine induced rapid rounding, loss of motility, cytotoxicity and rupture with no recovery after washout, supporting amoebicidal activity; in contrast, isometamidium produced a reversible non-motile pseudocyst-like phenotype with recovery after drug removal, consistent with amoebostasis. Benzoxaboroles (AN11736/AN5568) reduced motility without inducing pseudocyst formation or complete arrest and did not meet an amoebicidal threshold, suggesting sublethal cytostasis under the conditions tested. Several other classes showed no measurable effects by motility (paromomycin; diamidines; amphotericin B; suramin), while nitroheterocyclics (benznidazole, nifurtimox) generated PI-positive signals without corresponding motility or morphology changes, reinforcing the risk of false inference from nucleic-acid dye endpoints in xenic systems. It should be emphasised that the PLO-targeting rationale underpinning this study remains a hypothesis rather than a demonstrated mechanism, and a central interpretive question is therefore whether the active phenotypes reflect action on the amoeba host, the kinetoplastid endosymbiont (PLO), or both. Miltefosine has documented activity against free-living amoebae that do not harbour a PLO, implying that amoebicidal activity need not be PLO-mediated and may reflect broader membrane and organelle disruption mechanisms **[53], [54], [55], [56]**. For isometamidium, mitochondrial and kinetoplast DNA-associated modes of action in trypanosomes make PLO targeting plausible, but we currently lack direct evidence of target engagement in the endosymbiont, and the reversibility of the phenotype is compatible with transient bioenergetic disruption in the amoeba itself **[57]**. Resolving this will require targeted cell biology and ‘omics approaches: cytology (e.g., PLO-specific staining/quantification, ultrastructure), coupled with transcriptomics and/or metabolomics during exposure and recovery to determine whether PLO-associated pathways are preferentially perturbed relative to amoebal housekeeping processes **[58], [59]**. Progress is also constrained by genomics: the lack of a complete *N. perurans* reference hampers both mechanistic inference and host– symbiont dissection. While draft resources and endosymbiont-focused assemblies have been generated, the inability to culture axenically and high bacterial background have historically complicated assembly and annotation efforts **[14], [24], [60]**. A high-quality *N. perurans* genome will be pivotal for confidently identifying conserved drug targets (e.g., CPSF3 orthologues and mRNA processing machinery) and for interpreting whether candidate phenotypes are plausibly PLO-directed or host-directed **[61], [62], [63], [64]**.

*In vivo*, all four candidates were first assessed for tolerance under an IM injection regimen, with mortality clustering early and cortisol remaining similar across groups, suggesting that handling and repeated injection, rather than pharmacological toxicity, were the dominant drivers of acute **stress [65], [66], [67], [68]**. On welfare grounds, nifurtimox was not progressed despite the lack of significant group differences, as its group showed the highest mortality under the tolerance regimen **[69]**. In the field efficacy trial, miltefosine, isometamidium, and benznidazole were each associated with improved clinical gill scores after two weekly IM injections, indicating a measurable clinical benefit under natural infection pressure. However, qPCR outcomes did not mirror gill score improvements. Amoebic DNA loads were unexpectedly higher in the benznidazole and isometamidium groups than in controls, and miltefosine did not show clear reduction relative to controls. Several non-exclusive explanations for this result are consistent with the biology and the measurement approach **[36], [70], [71].** First, and most importantly, conventional qPCR quantifies DNA rather than viable amoebae, and therefore cannot distinguish living, pathogenic trophozoites from arrested, dying, or recently killed cells whose genomic material persists on the gill surface **[36], [70], [71]**. Intact but arrested pseudocysts can retain genomic material and may be more readily detached and collected during swabbing, potentially inflating apparent load despite reduced pathogenic activity, particularly under amoebostatic pressure. This limitation complicates interpretation of therapeutic benefit in our study, as defined by the improvement in gross of gill pathology. Alternate approaches to parasite enumeration to target only viable cells might be considered in future (e.g propidium monoazide (PMA)-qPCR **[72])**. Second, sampling design and subsample size may limit inference. Gill score and qPCR were correlated, but qPCR was performed on a subset of all samples. As such, treatment contrasts may be sensitive to stochastic sampling and within-pen heterogeneity. Third, timing relative to pharmacokinetics is likely crucial: for miltefosine in particular, effective therapy in humans depends on sustained exposure and slow tissue accumulation over prolonged dosing schedules, while fish-specific absorption, distribution and elimination are unknown. Sample may therefore have missed the window when amoebicidal activity would most strongly translate into reduced detectable load **[73], [74], [75]**. Finally, host-directed effects may contribute to the clinical signal. Benznidazole has immunomodulatory effects in mammalian systems, and a reduction in gill pathology without corresponding parasite DNA decline is compatible with a shift in inflammatory pathology rather than direct amoebicidal activity **[76], [77]**. **[71], [77].** Together, these observations emphasise that clinical outcomes and conventional molecular diagnostics may decouple in AGD trials, and further studies are necessary.

From a translational perspective, the use of the drugs we have deployed may face several barriers. Deployment will depend on fish-specific pharmacokinetics, residue considerations, welfare impacts, and regulatory acceptability. At present, none of the compounds evaluated have established pharmacokinetic profiles in salmonids, limiting interpretation of efficacy and dose selection; future work should prioritise PK/PD profiling with multi-timepoint plasma and tissue sampling to define exposure windows and therapeutic indices **[78], [79], [80], [81], [82]**. IM dosing was deployed in this trial to minimise environmental contamination and to control for handling stress, as isometamidium must be administered via the IM route (www.fao.org). However, IM is labour-intensive and itself contributed to clustered early mortality, future work should also evaluate alternative delivery routes (oral or immersion) and/or administration strategies that reduce handling and improve scalability **[83], [84], [85]**. Little is known about the environmental toxicity and persistence of the drugs deployed in this study, although one might expect that they are no worse than the broad spectrum antiparasitic currently deployed in salmonid aquaculture (e.g. deltamethrin, ivermectin and azamethiphos **[86]**), and further work is required. Finally, while the cost of some repurposed drugs is a barrier in their original indications, aquaculture represents a potential new market that could, in principle, incentivise manufacturing efficiencies and support broader access though this remains speculative and would require careful stewardship to avoid unintended consequences for human and veterinary medicine.

## 5. Conclusions

In conclusion, this study demonstrates that repurposing trypanocidal drugs for AGD is scientifically plausible and operationally testable, but only if assay systems and endpoints are chosen to reflect the biological realities of xenic culture and stage plasticity. Digital holographic microscopy offered the most amoeba-informative in vitro readout and enabled prioritisation of candidates for field evaluation. *In vivo*, miltefosine, isometamidium and benznidazole produced measurable clinical improvement in gill scores under natural AGD exposure, yet molecular load estimates by conventional qPCR did not track these outcomes. The next phase of development should therefore focus on (i) automating holographic analysis using machine learning to enable scalable screening, (ii) establishing whether candidate drugs act via the PLO or the amoeba host using targeted cytology and ‘omics, (iii) generating salmonid PK/PD data to rationalise dosing and sampling windows, and (iv) adopting viability-sensitive molecular diagnostics to reconcile clinical and qPCR endpoints. Together, these steps will determine whether trypanocidal repurposing can progress from proof-of-concept to practical therapeutics for AGD control in salmon aquaculture.

## Author Contributions

Y.W.L., A.L.E.B., B.R., P.M., and M.L. drafted the manuscript. M.B., M.L., N.R., P.M., R.B., and F.H. conceived and designed the study and contributed to manuscript revision. K.D. contributed to manuscript drafting and revision. S.M., L.C., F.E., J.M., D.O., and B.C. performed the experimental work.

## Funding

This work was supported by UKRI grants BB/T016280/1 and BB/Y012437/1, and by the Sustainable Aquaculture Innovation Centre PhD Scheme. The funders had no role in study design, data collection, analysis, interpretation, or the decision to submit for publication.

## Abbreviations

The following abbreviations are used in this manuscript:

AGD: Amoebic Gill Disease

## Supplementary material

### S1 Different concentration of Propidium iodide used and viewed under fluorescent microscope

To identify the optimum concentration of propidium iodide if it can identify dead amoeba, propidium iodide of different concentration were added to formaldehyde fixed and no-treatment amoeba. As shown in Figure A, no staining occurs in the healthy amoeba but in the fixed amoeba which suggests that we can distinguish dead cells by propidium iodide. Furthermore, the optimal concentration of propidium iodide is 20µM as the fluorescence seems to be faint at lower concentrations but remains similar fluorescence level even in the concentration higher than 20µM. As a result, 20µM of propidium iodide was selected to be used in drug screening against N. perurans.

**Figure S1.**
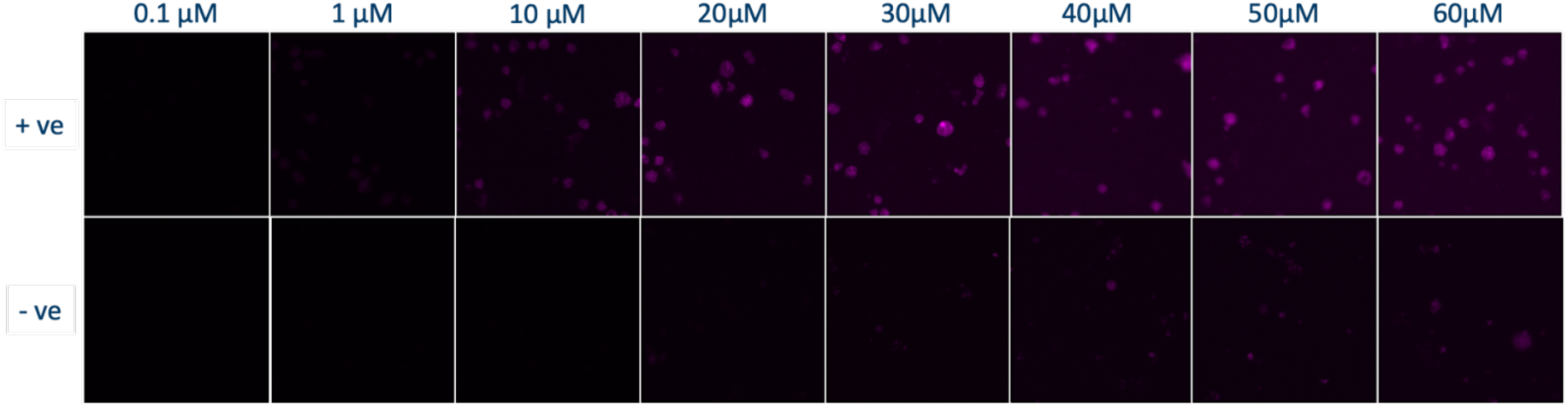
Formaldehyde killed Neoparamoeba perurans (+ve) and living Neoaramoeba perurans (-ve) were stained with different concentrations of propidium iodide from 0.1µM to 60µM.

### S2 Intracellular bacterial extraction using different methods

To develop effective lysing methods for eliminating amoebae while preserving the bacterial population from the non-axenically cultured *Neoparamoeba perurans*, three different methods were utilised: (1) manually breaking amoebae through rapid passaging with a gauged needle, water bath sonication, and the use of the isobiotec cell homogenizer. Despite attempting 50 passages with the fine gauge needle, it was insufficient in eliminating all amoebae (data not shown). (2) Similarly, 30 minutes of sonication also failed to eradicate all amoeba (data not shown). (3) The isobiotec cell homogeniser was particularly of interest due to its potential to break amoebae at 4 and 6 microns with 20 strokes (Figure A). By using isobiotec cell homogeniser, fluorescence and luminescence activity of the extracted extracellular and intracellular bacteria (hereafter homogenised amoebae) can be used as a background control, by comparing to the activity of non-axenically cultured *N. perurans* (hereafter non-homogenised amoebae), facilitating the estimation of the fluorescence and luminescence level originating from the amoebae themselves when conducting CellTox Green Cytotoxicity assay and CellTitre Glo Cell Viability assay.

**Figure S2.**
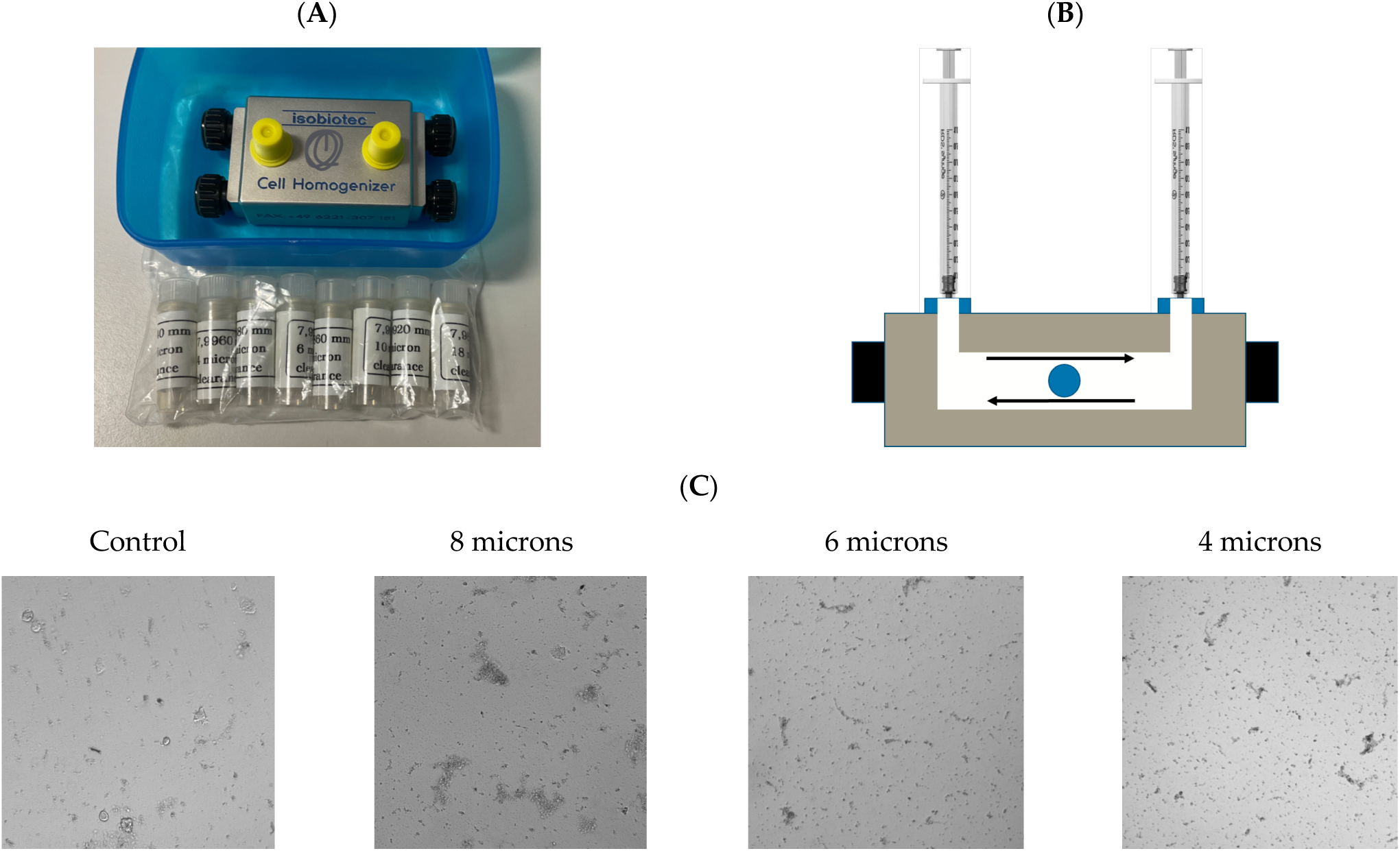
Preparation of intracellular and extracellular bacteria from *Neoparamoeba perurans* cultures using a Balch cell homogeniser. (A) Photograph of the isobiotec Balch homogeniser. (B) Schematic of the homogeniser chamber. A stainless-steel ball and an amoeba suspension in malt yeast broth were loaded into the chamber, then the sample was repeatedly forced through the narrow annular gap between the ball and the chamber wall to rupture cells and generate a homogenate. Balls of 8 µm, 6 µm and 4 µm diameter were used, and 20 strokes were applied. (C) Representative bright-field micrographs acquired at ×10 on a Leica DMi8 microscope, showing disrupted *N. perurans* cells of approximately 10–20 µm and release of associated bacteria.

### S3. Single Cell Tracking Analysis

#### Miltefosine

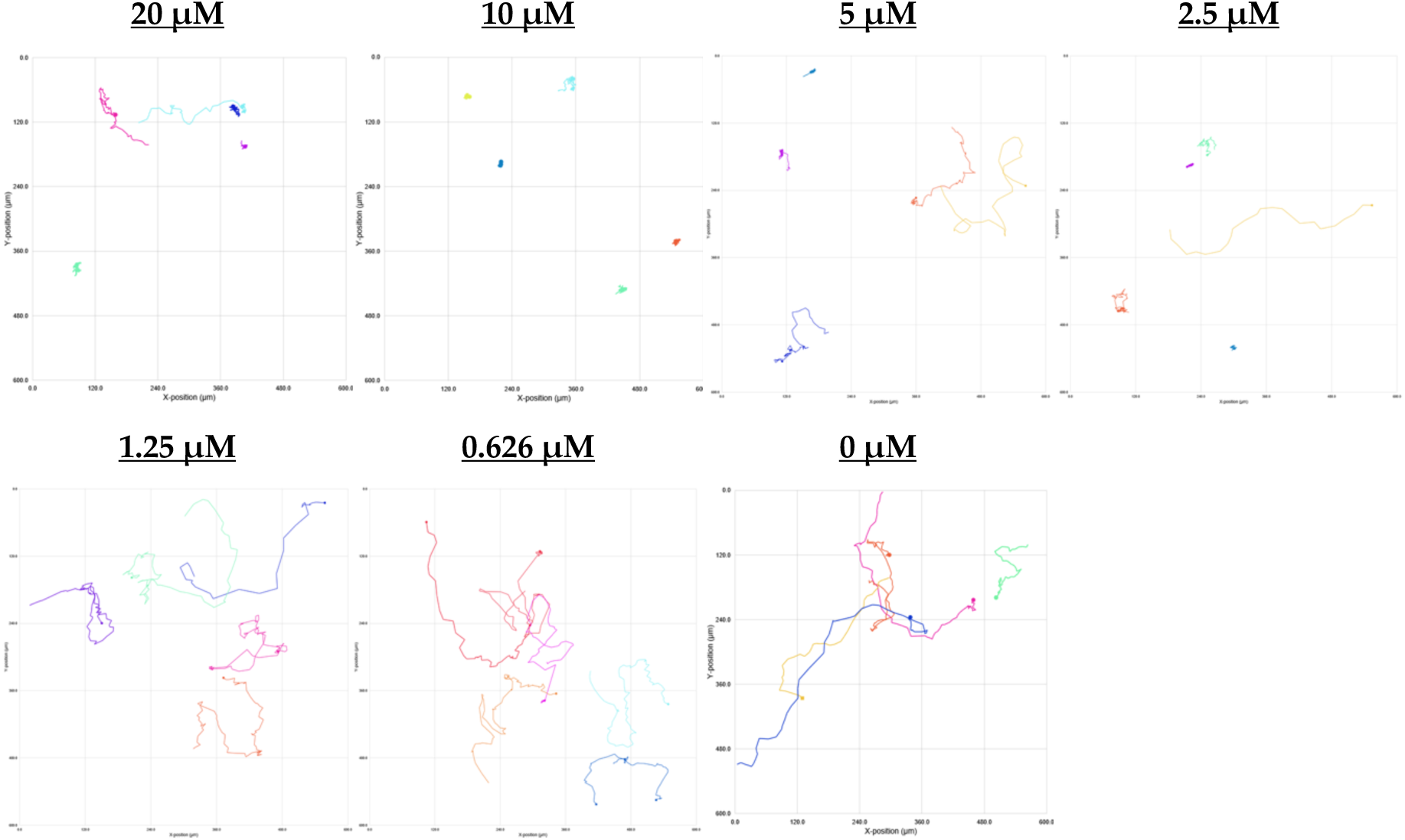

Single cell tracking analysis of *N. perurans* motility following exposure to miltefosine during a 4-hrs analysis windows at 0 h. The tested concentrations ranging from 20 µM to 0.625 µM. Each plot represents the movement of a single amoeba based on its X and Y coordinates captured by HoloMonitor at one randomly selected position within the well during the initial 6 hours of treatment. Lines depict the path each amoeba migrated during this period. Different colours indicate different amoeba being tracked.

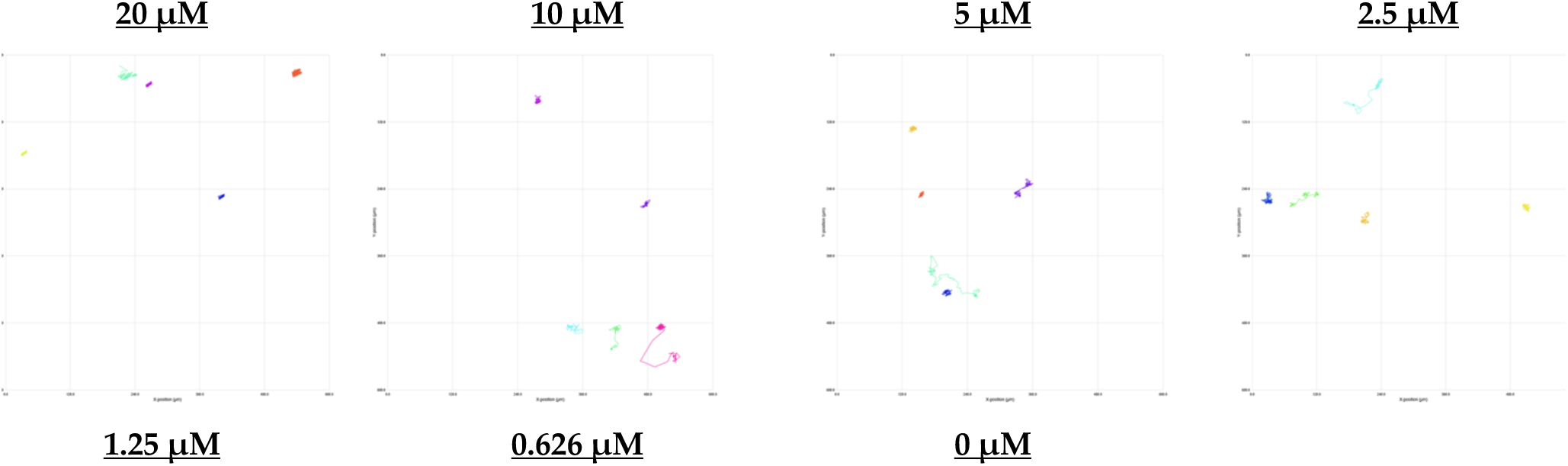
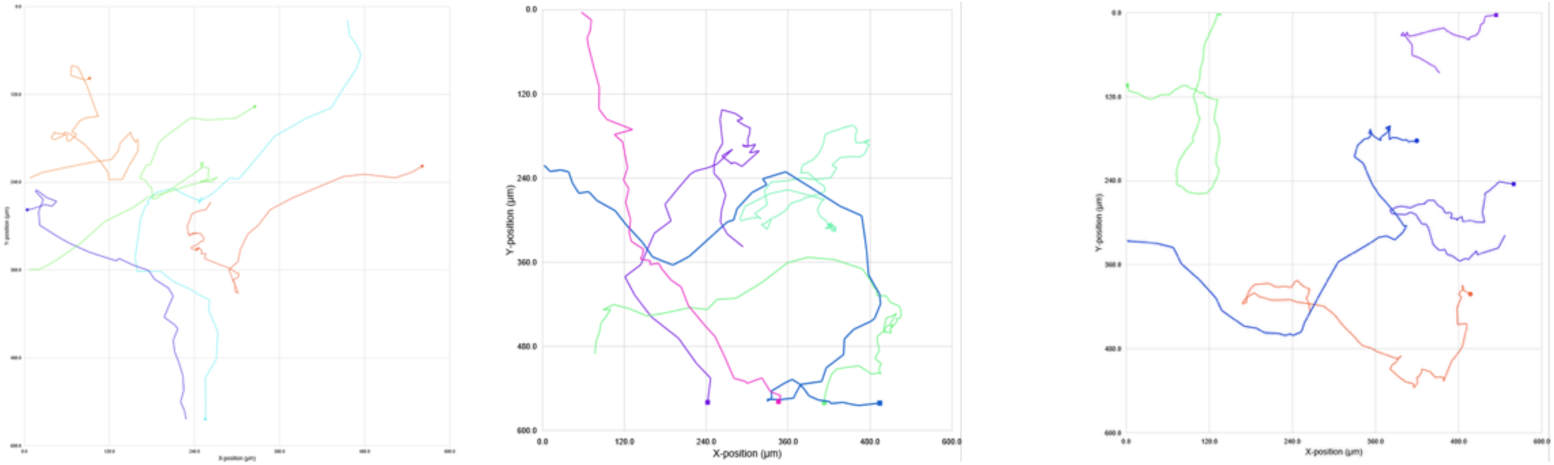

Single cell tracking analysis of *N. perurans* motility following exposure to miltefosine during a 4-hrs analysis windows at 24 h. The tested concentrations ranging from 20 µM to 0.625 µM. Each plot represents the movement of a single amoeba based on its X and Y coordinates captured by HoloMonitor at one randomly selected position within the well during the initial 6 hours of treatment. Lines depict the path each amoeba migrated during this period. Different colours indicate different amoeba being tracked.

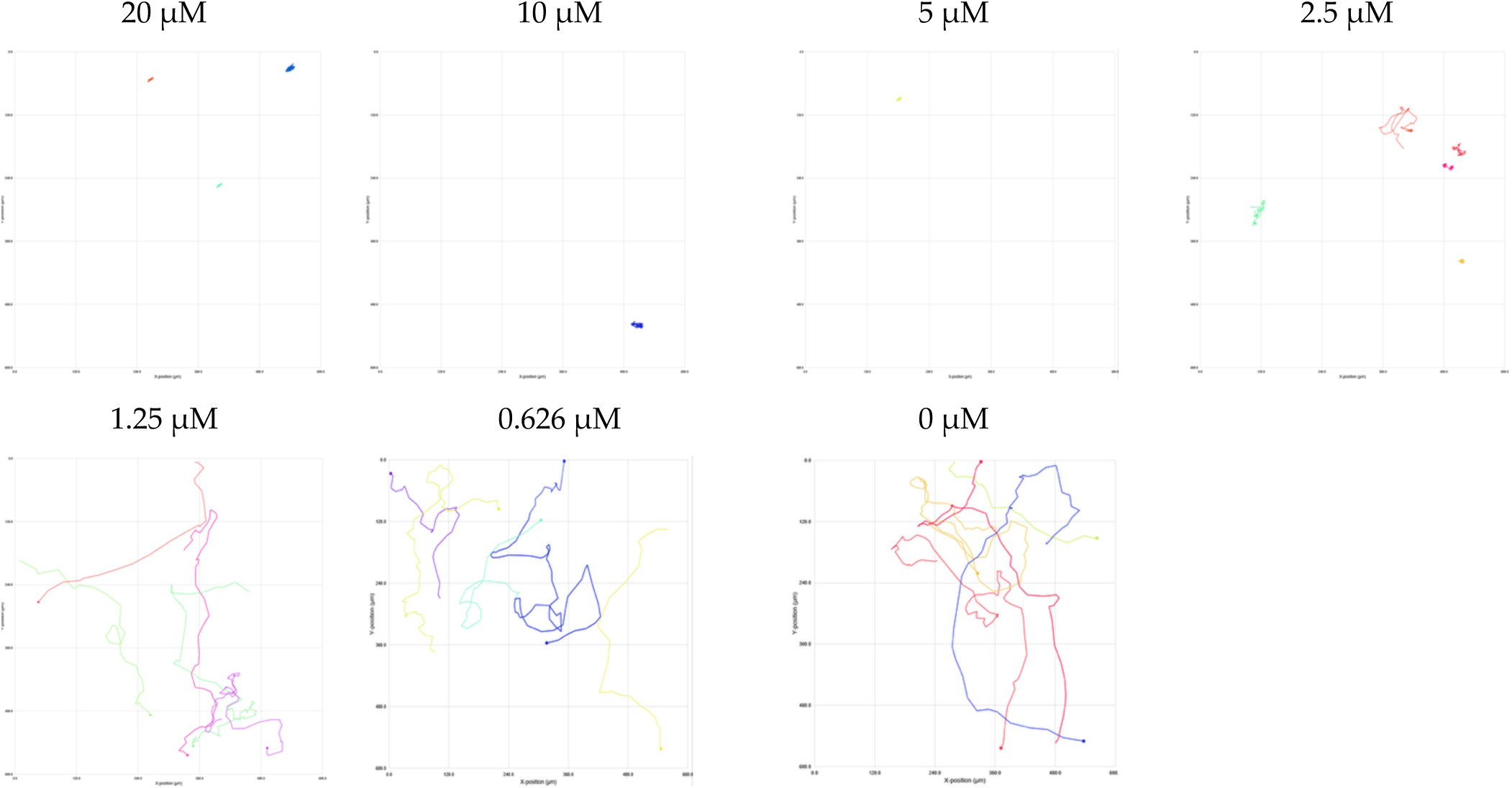

Single cell tracking analysis of N. perurans motility following exposure to miltefosine during a 4-hrs analysis windows at 48 h. The tested concentrations ranging from 20 µM to 0.625 µM. Each plot represents the movement of a single amoeba based on its X and Y coordinates captured by HoloMonitor at one randomly selected position within the well during the initial 6 hours of treatment. Lines depict the path each amoeba migrated during this period. Different colours indicate different amoeba being tracked.

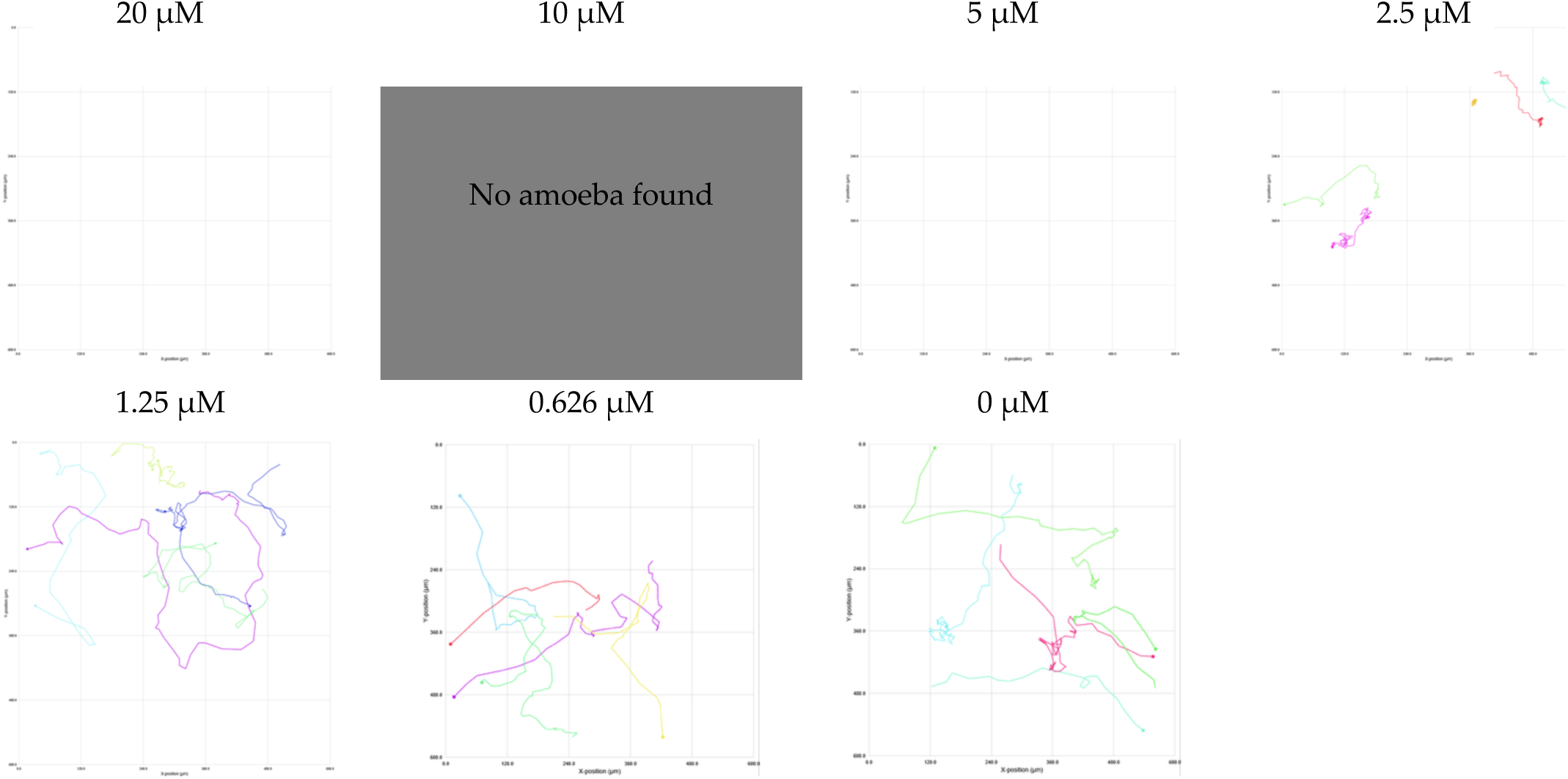

Single cell tracking analysis of N. perurans motility following exposure to miltefosine during a 4-hrs analysis windows at 72 h. The tested concentrations ranging from 20 µM to 0.625 µM. Each plot represents the movement of a single amoeba based on its X and Y coordinates captured by HoloMonitor at one randomly selected position within the well during the initial 6 hours of treatment. Lines depict the path each amoeba migrated during this period. Different colours indicate different amoeba being tracked.

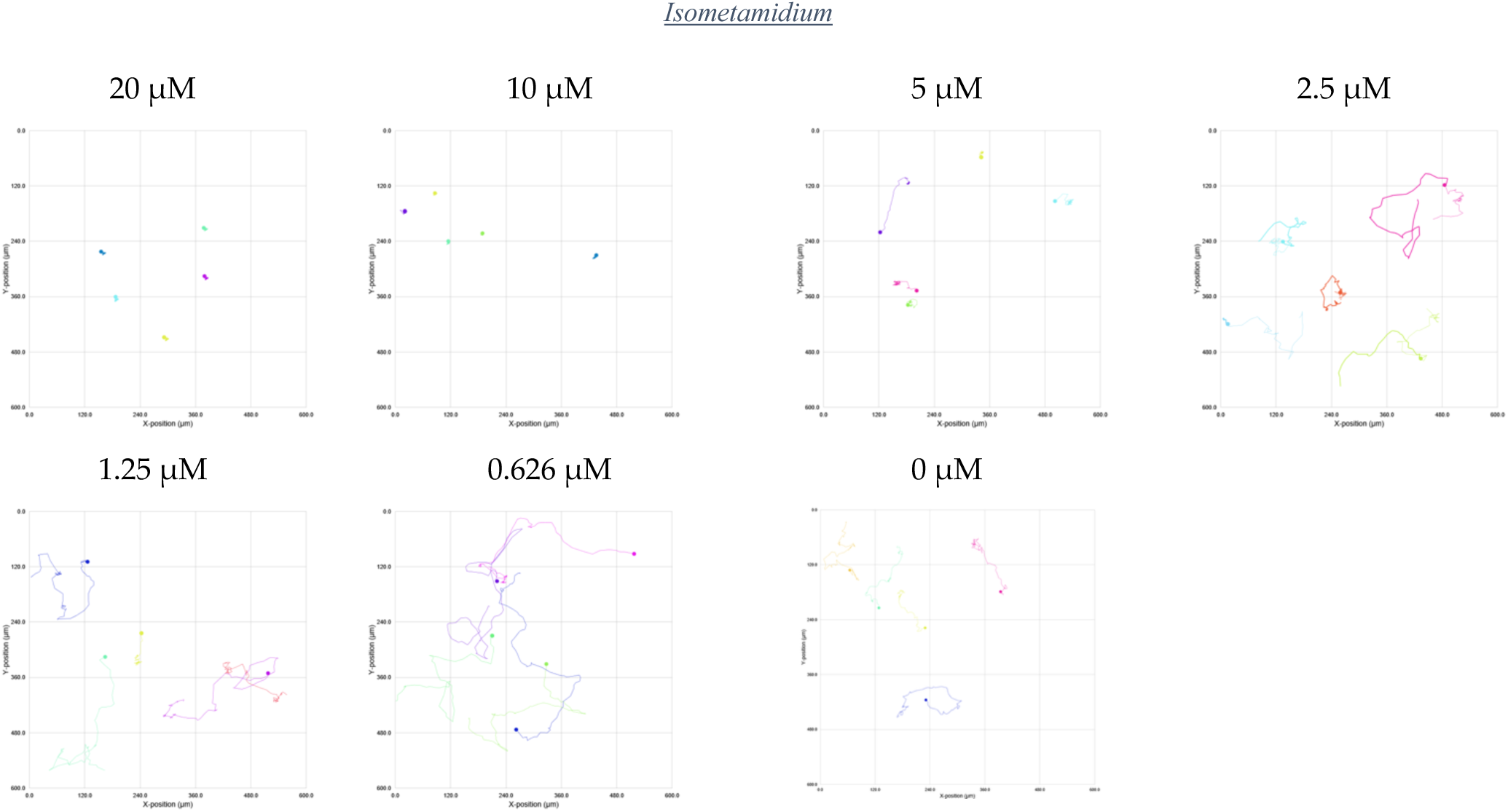

Single cell tracking analysis of N. perurans motility following exposure to isometamidium during a 4-hrs analysis windows at 0 h. The tested concentrations ranging from 20 µM to 0.625 µM. Each plot represents the movement of a single amoeba based on its X and Y coordinates captured by HoloMonitor at one randomly selected position within the well during the initial 6 hours of treatment. Lines depict the path each ameba migrated during this period. Different colours indicate different amoeba being tracked.

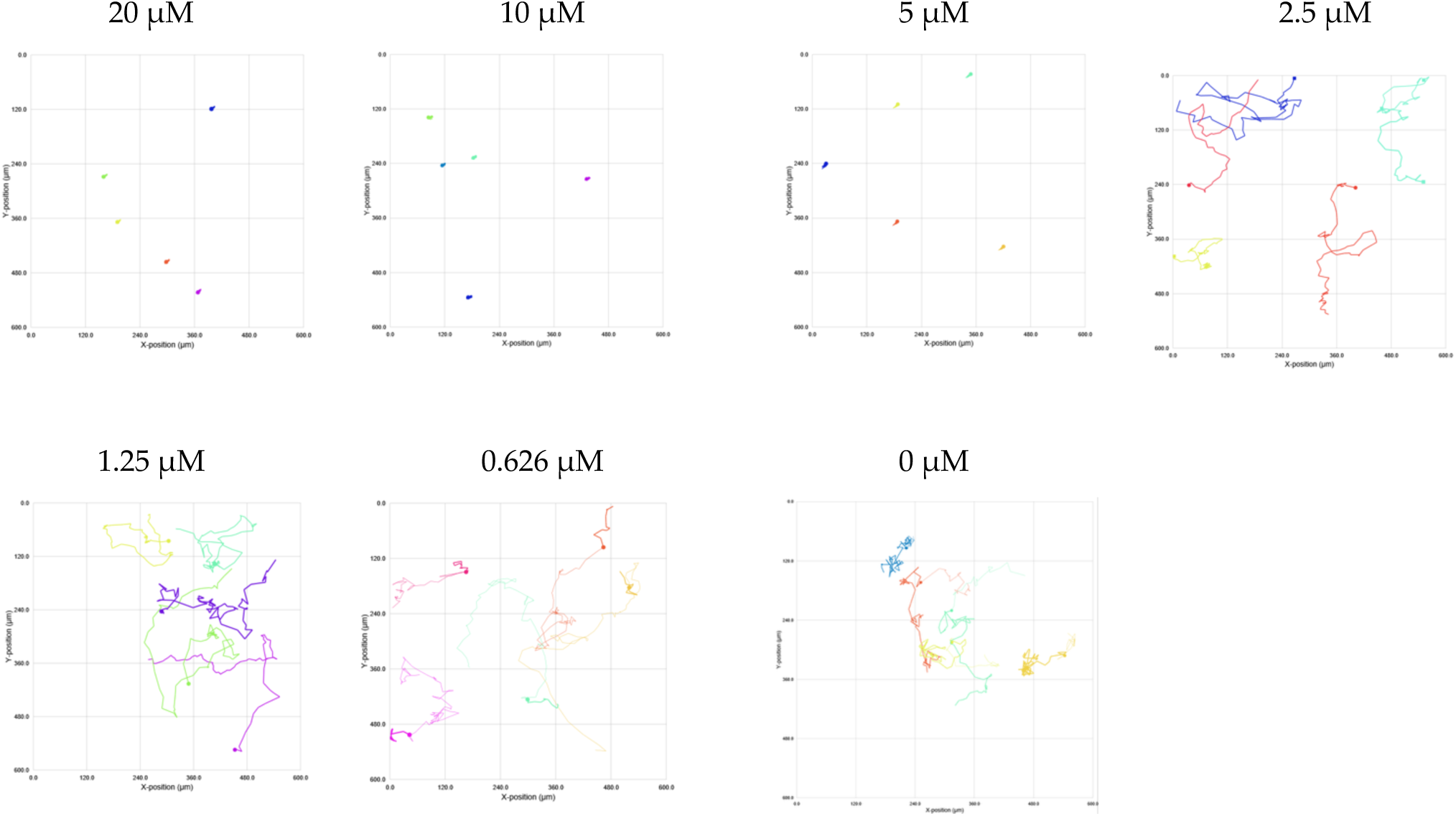

Single cell tracking analysis of *N. perurans* motility following exposure to isometamidium during a 4-hrs analysis windows at 24 h. The tested concentrations ranging from 20 µM to 0.625 µM. Each plot represents the movement of a single amoeba based on its X and Y coordinates captured by HoloMonitor at one randomly selected position within the well during the initial 6 hours of treatment. Lines depict the path each amoeba migrated during this period. Different colours indicate different amoeba being tracked.

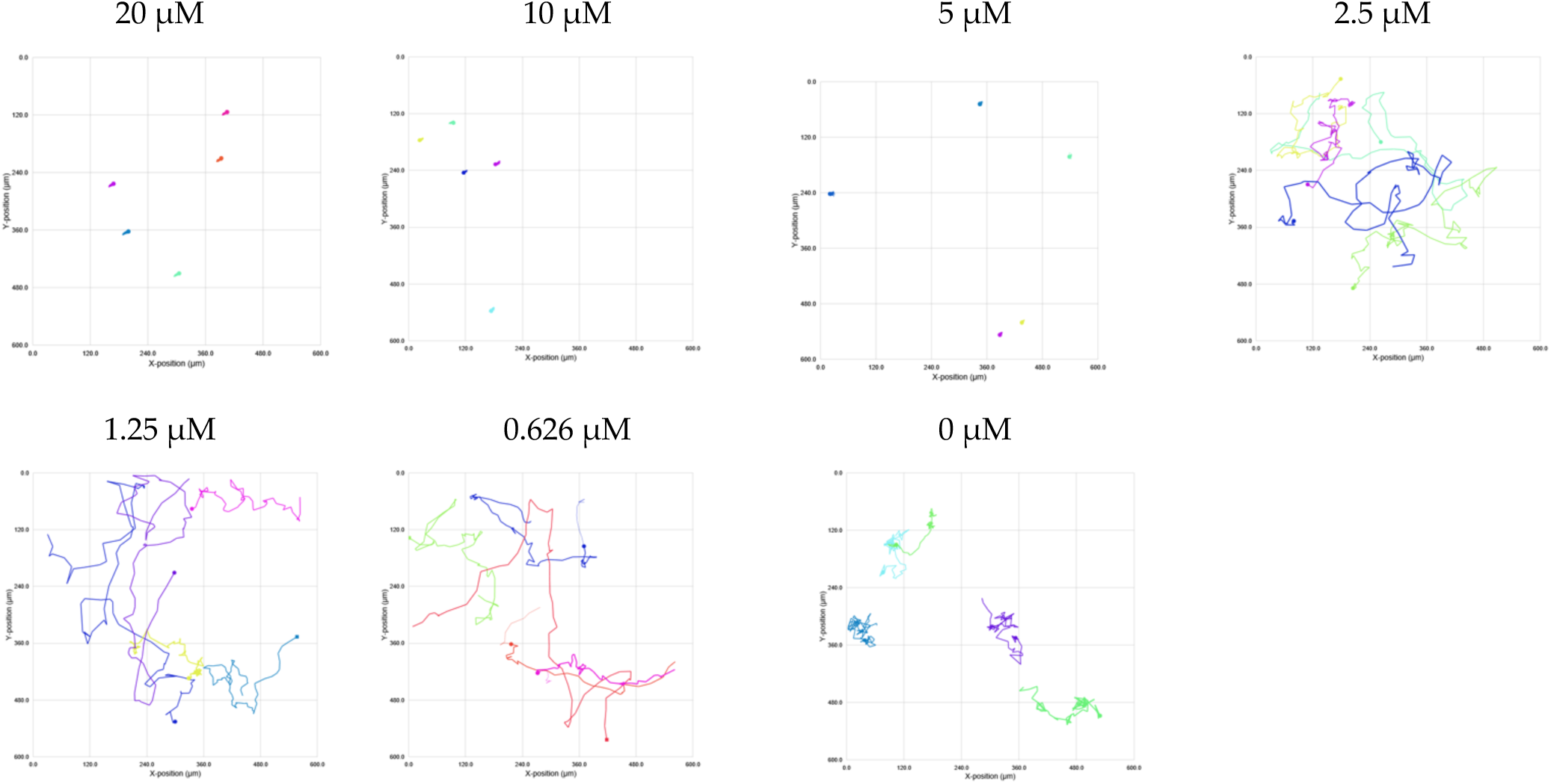

Single cell tracking analysis of *N. perurans* motility following exposure to isometamidium during a 4-hrs analysis windows at 48 h. The tested concentrations ranging from 20 µM to 0.625 µM. Each plot represents the movement of a single amoeba based on its X and Y coordinates captured by HoloMonitor at one randomly selected position within the well during the initial 6 hours of treatment. Lines depict the path each amoeba migrated during this period. Different colours indicate different amoeba being tracked.

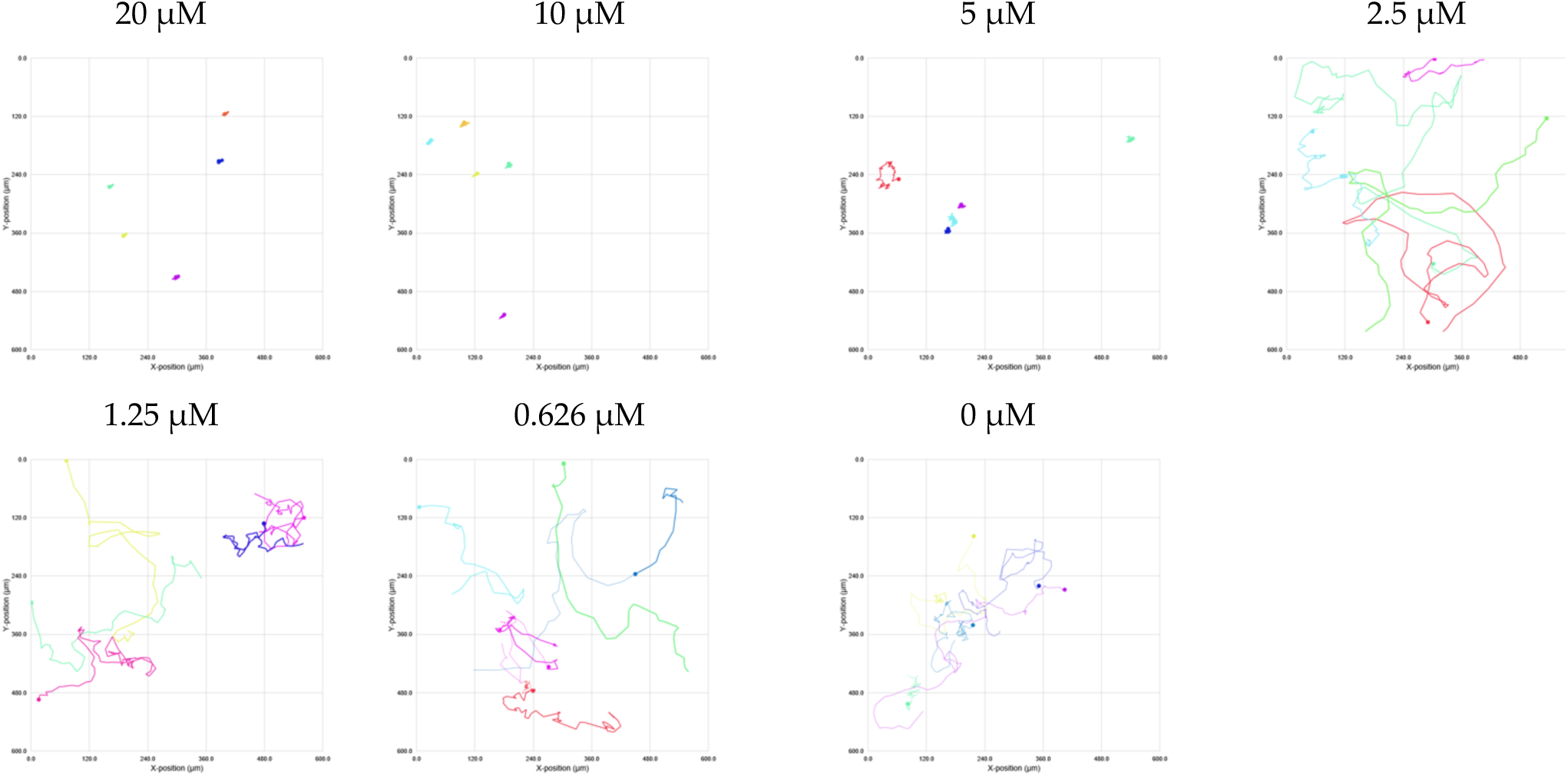

Single cell tracking analysis of *N. perurans* motility following exposure to isometamidium during a 4-hrs analysis windows at 72 h. The tested concentrations ranging from 20 µM to 0.625 µM. Each plot represents the movement of a single amoeba based on its X and Y coordinates captured by HoloMonitor at one randomly selected position within the well during the initial 6 hours of treatment. Lines depict the path each amoeba migrated during this period. Different colours indicate different amoeba being tracked.

#### AN11736

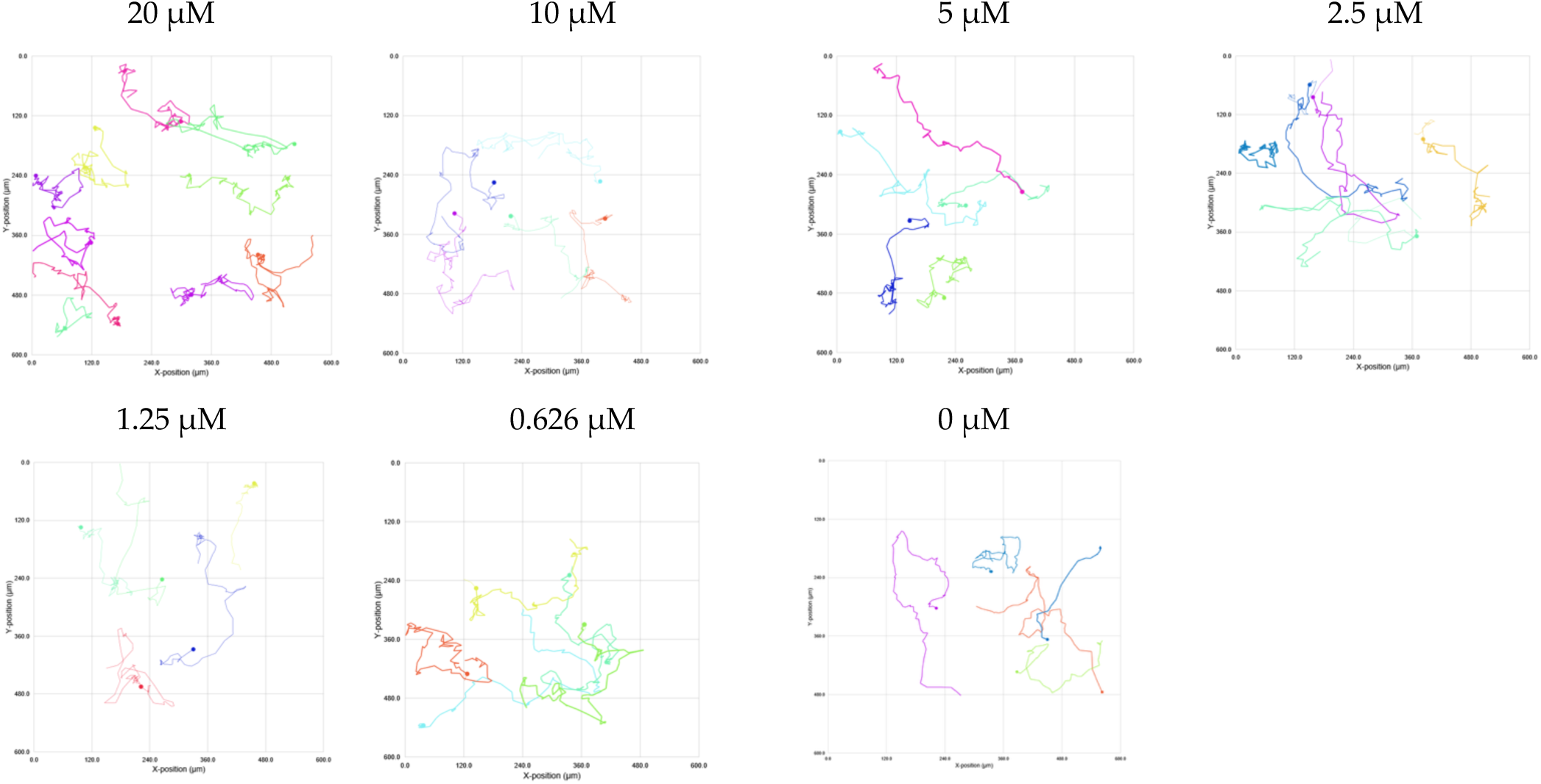

Single cell tracking analysis of *N. perurans* motility following exposure to AN11736 during a 4-hrs analysis windows at 0 h. The tested concentrations ranging from 20 µM to 0.625 µM.Each plot represents the movement of a single amoeba based on its X and Y coordinates captured by HoloMonitor at one randomly selected position within the well during the initial 6 hours of treatment. Lines depict the path each amoeba migrated during this period. Different colours indicate different amoeba being tracked.

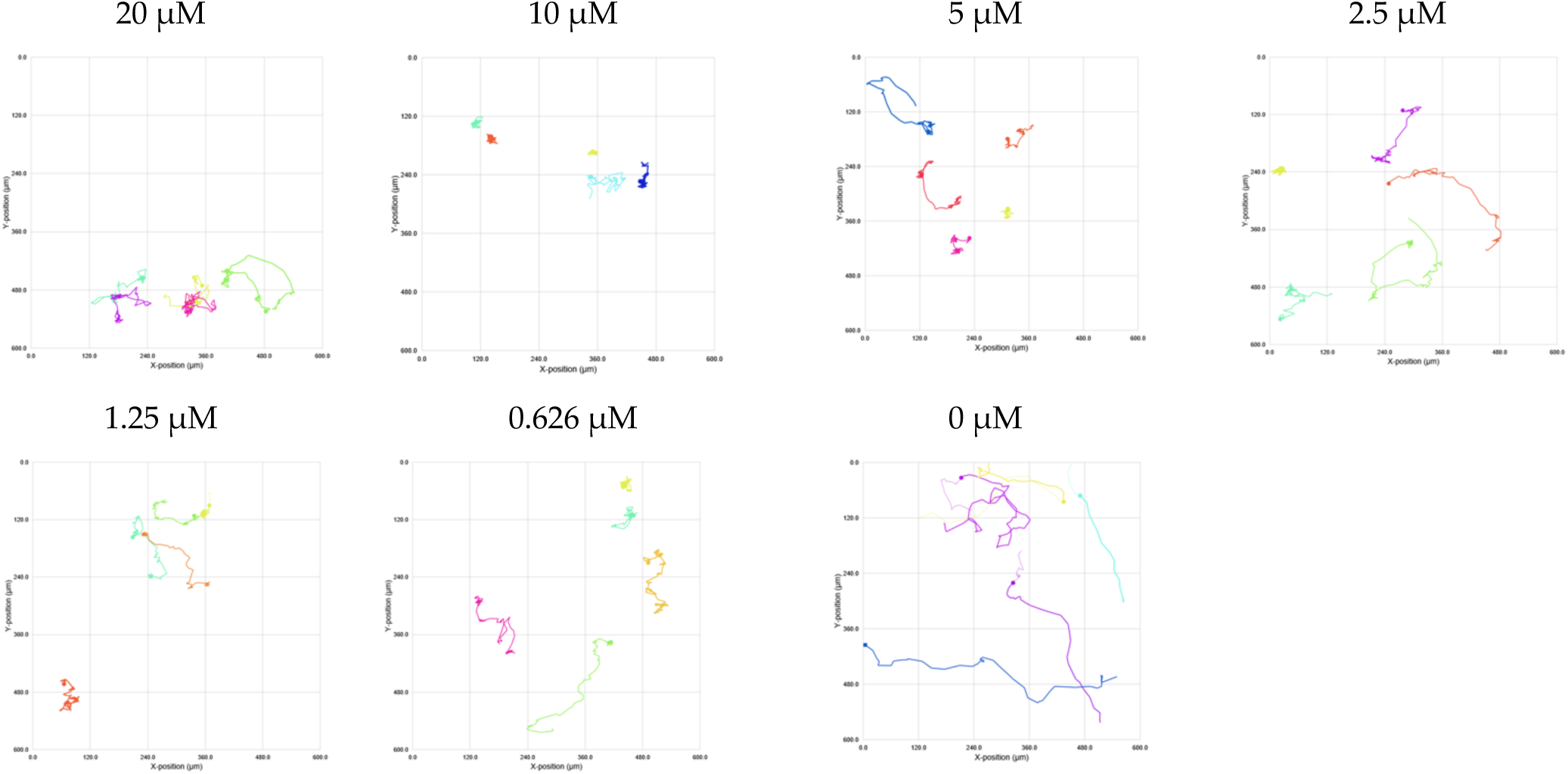

Single cell tracking analysis of *N. perurans* motility following exposure to AN11736 during a 4-hrs analysis windows at 24 h. The tested concentrations ranging from 20 µM to 0.625 µM.Each plot represents the movement of a single amoeba based on its X and Y coordinates captured by HoloMonitor at one randomly selected position within the well during the initial 6 hours of treatment. Lines depict the path each amoeba migrated during this period. Different colours indicate different amoeba being tracked.

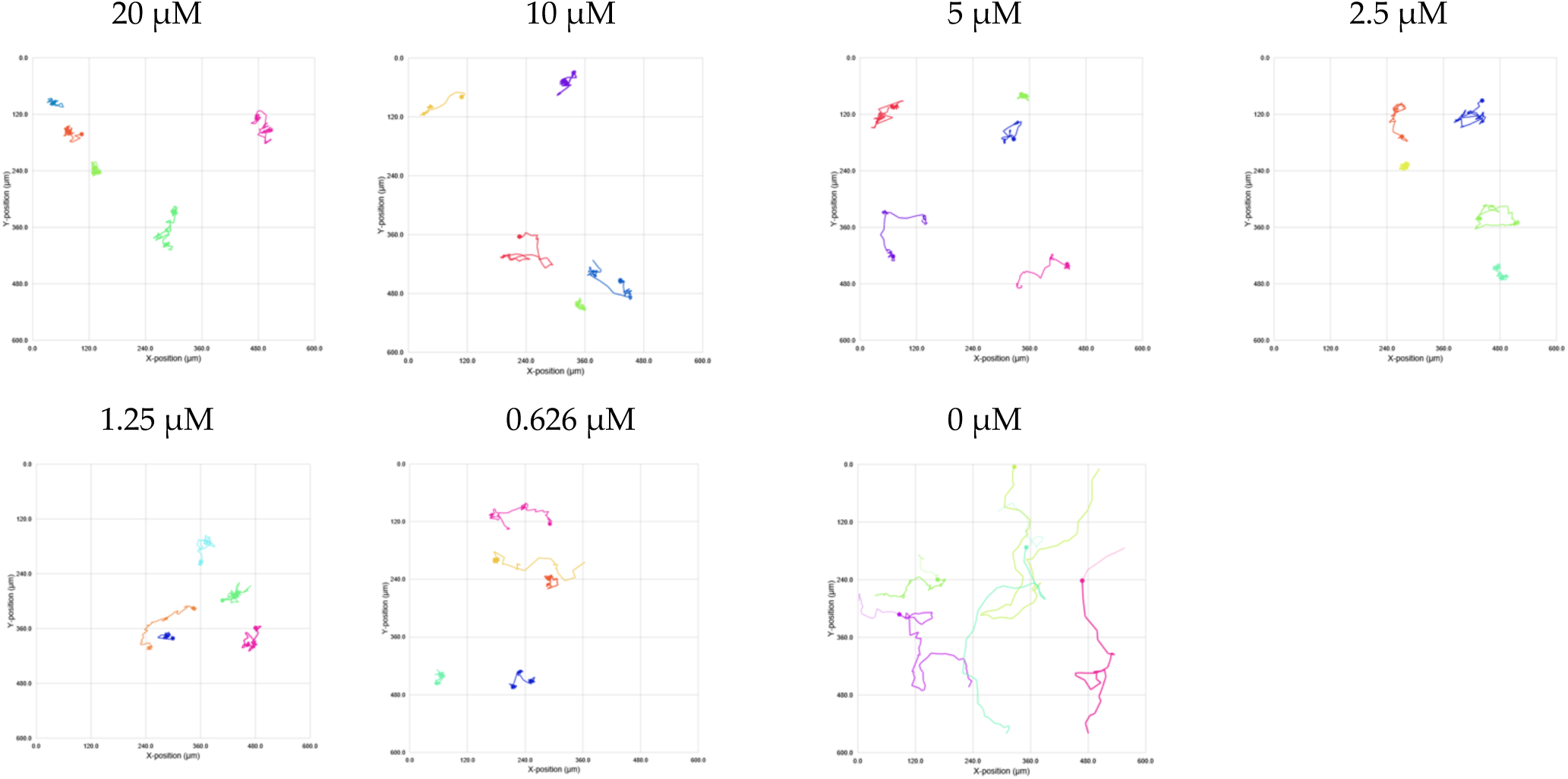

Single cell tracking analysis of *N. perurans* motility following exposure to AN11736 during a 4-hrs analysis windows at 48 h. The tested concentrations ranging from 20 µM to 0.625 µM. Each plot represents the movement of a single amoeba based on its X and Y coordinates captured by HoloMonitor at one randomly selected position within the well during the initial 6 hours of treatment. Lines depict the path each amoeba migrated during this period. Different colours indicate different amoeba being tracked.

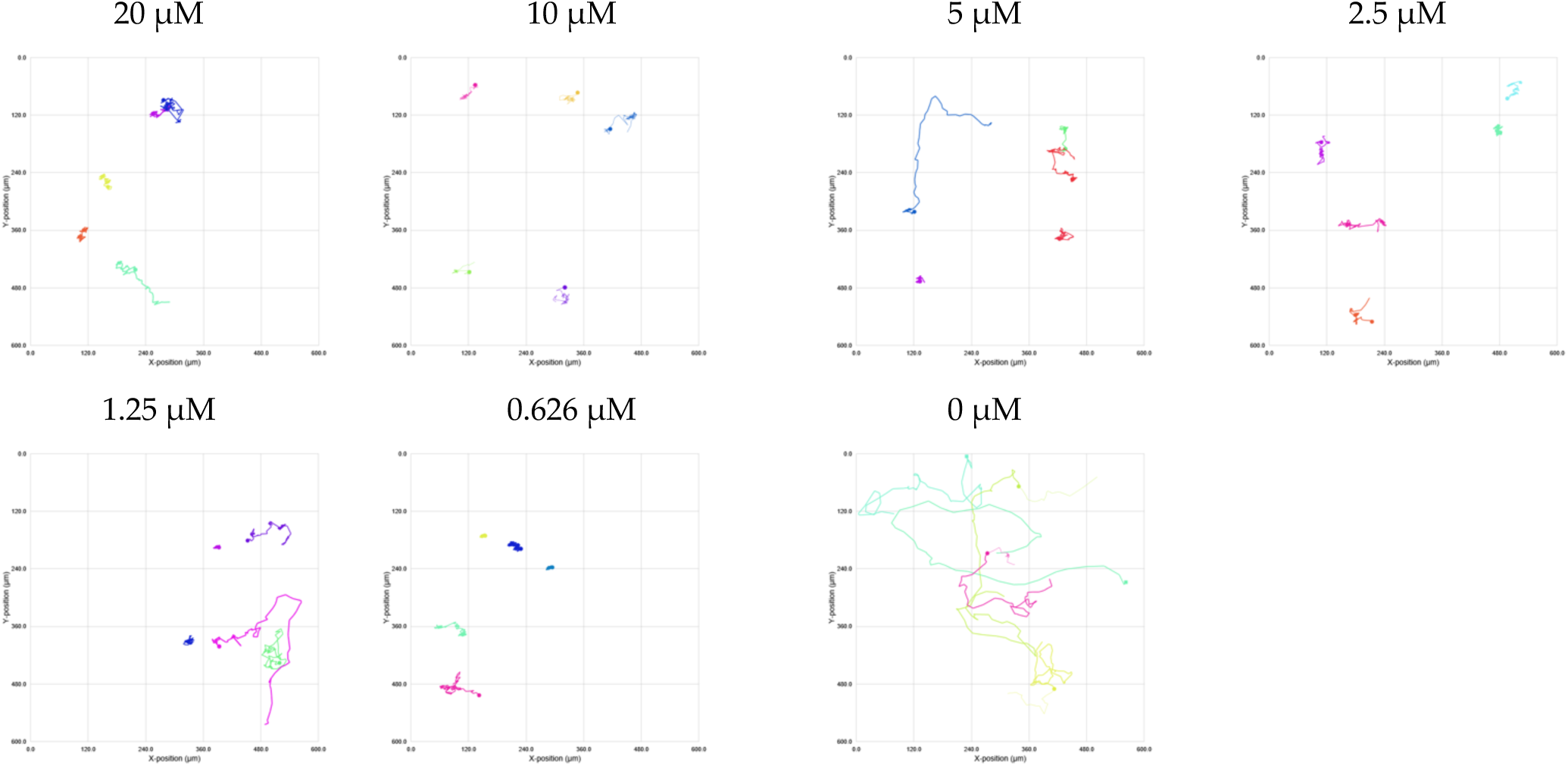

Single cell tracking analysis of *N. perurans* motility following exposure to AN11736 during a 4-hrs analysis windows at 72 h. The tested concentrations ranging from 20 µM to 0.625 µM. Each plot represents the movement of a single amoeba based on its X and Y coordinates captured by HoloMonitor at one randomly selected position within the well during the initial 6 hours of treatment. Lines depict the path each amoeba migrated during this period. Different colours indicate different amoeba being tracked.

#### Pentamidine

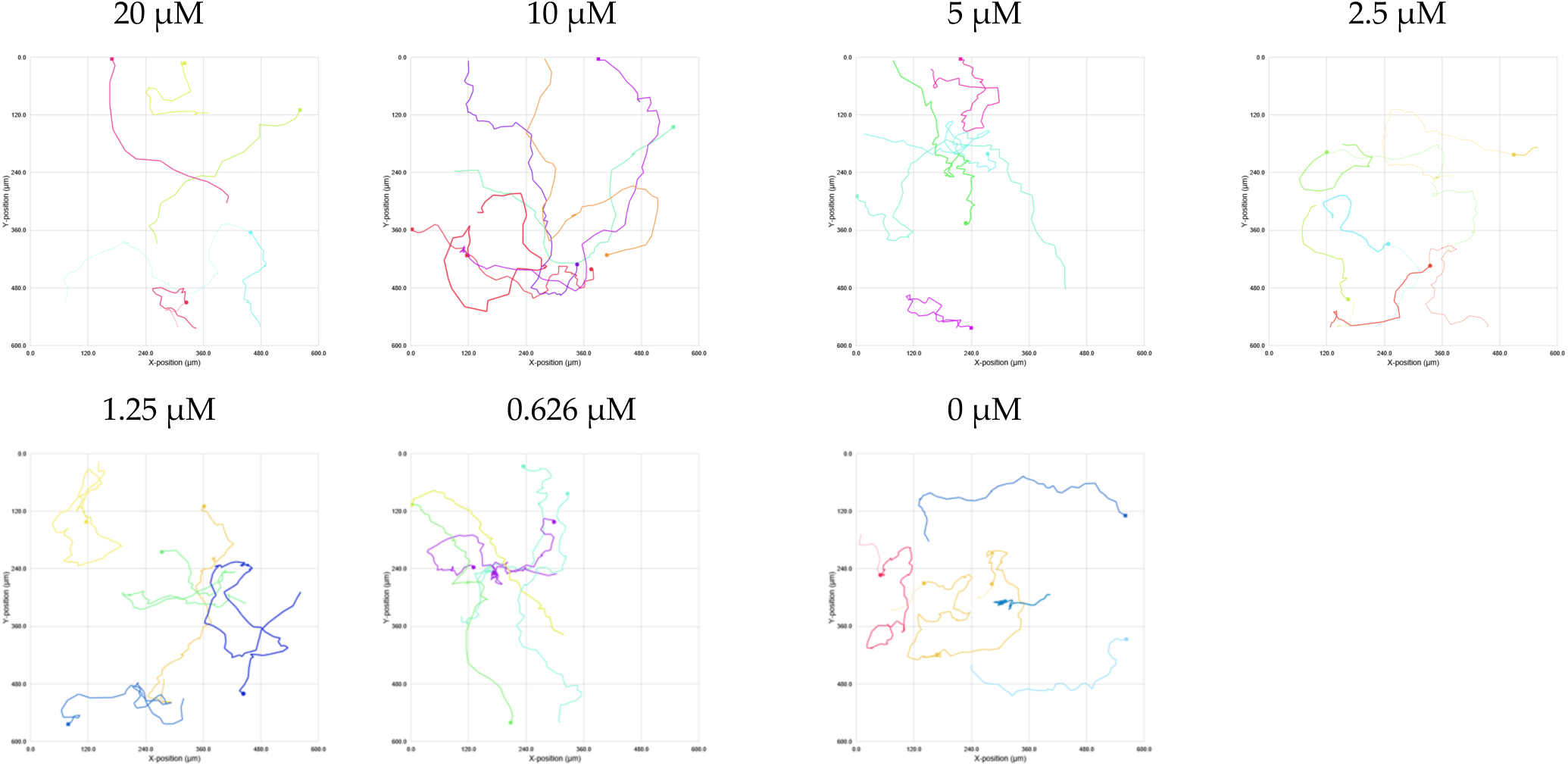

Single cell tracking analysis of *N. perurans* motility following exposure to pentamidine during a 4-hrs analysis windows at 0 h. The tested concentrations ranging from 20 µM to 0.625 µM. Each plot represents the movement of a single amoeba based on its X and Y coordinates captured by HoloMonitor at one randomly selected position within the well during the initial 6 hours of treatment. Lines depict the path each amoeba migrated during this period. Different colours indicate different amoeba being tracked.

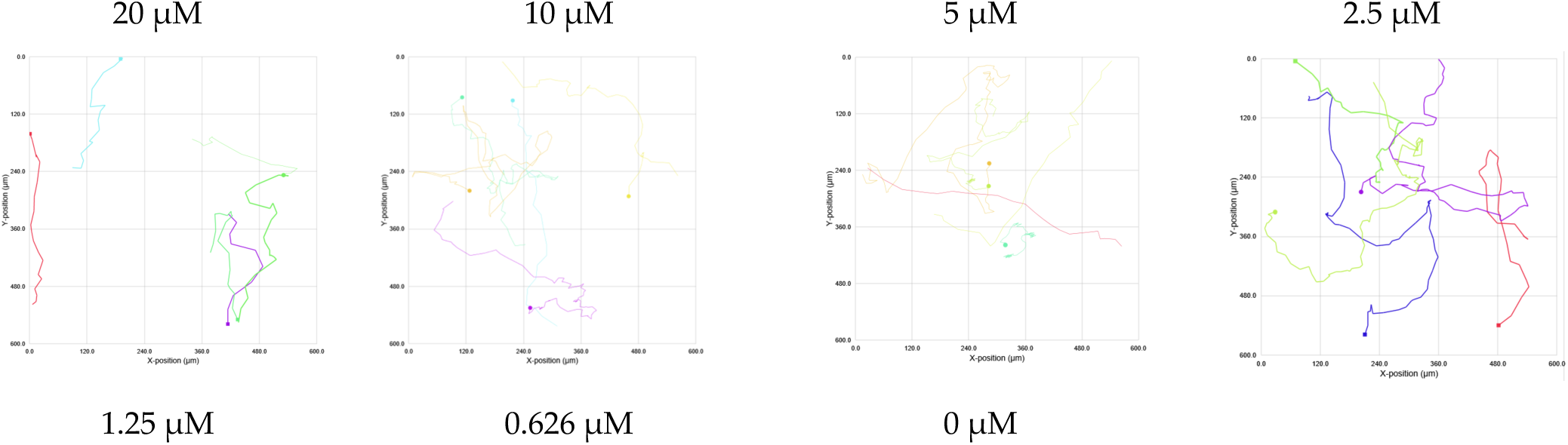
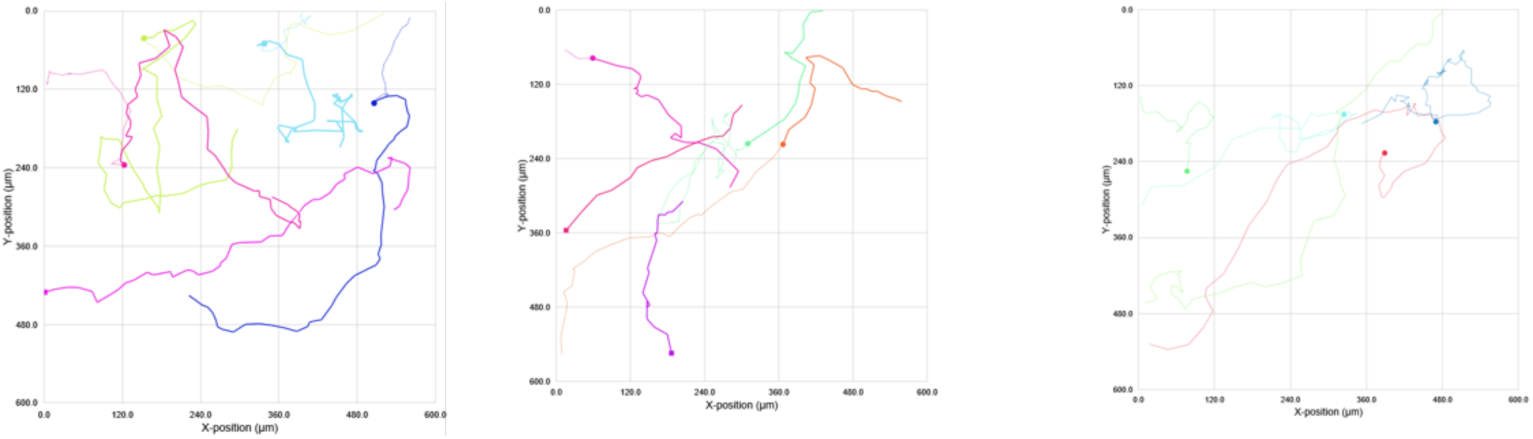

Single cell tracking analysis of *N. perurans* motility following exposure to pentamidine during a 4-hrs analysis windows at 24 h. The tested concentrations ranging from 20 µM to 0.625 µM. Each plot represents the movement of a single amoeba based on its X and Y coordinates captured by HoloMonitor at one randomly selected position within the well during the initial 6 hours of treatment. Lines depict the path each amoeba migrated during this period. Different colours indicate different amoeba being tracked.

### S4 Fluorescence microscopy images of in vitro grown Neoparamoeba perurans following treatment with eight drug classes of trypanocidals at different concentrations

**Figure S4.**
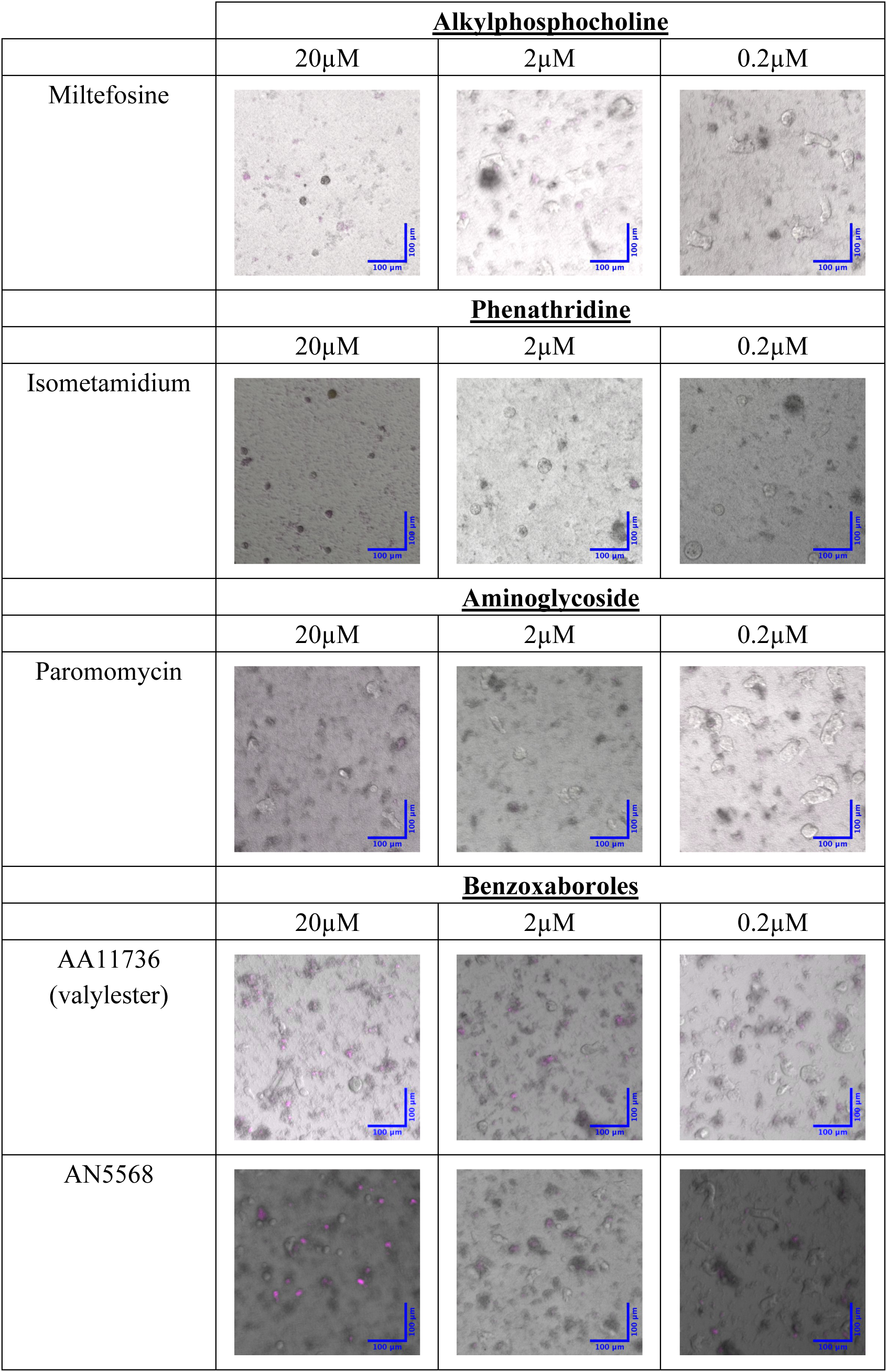
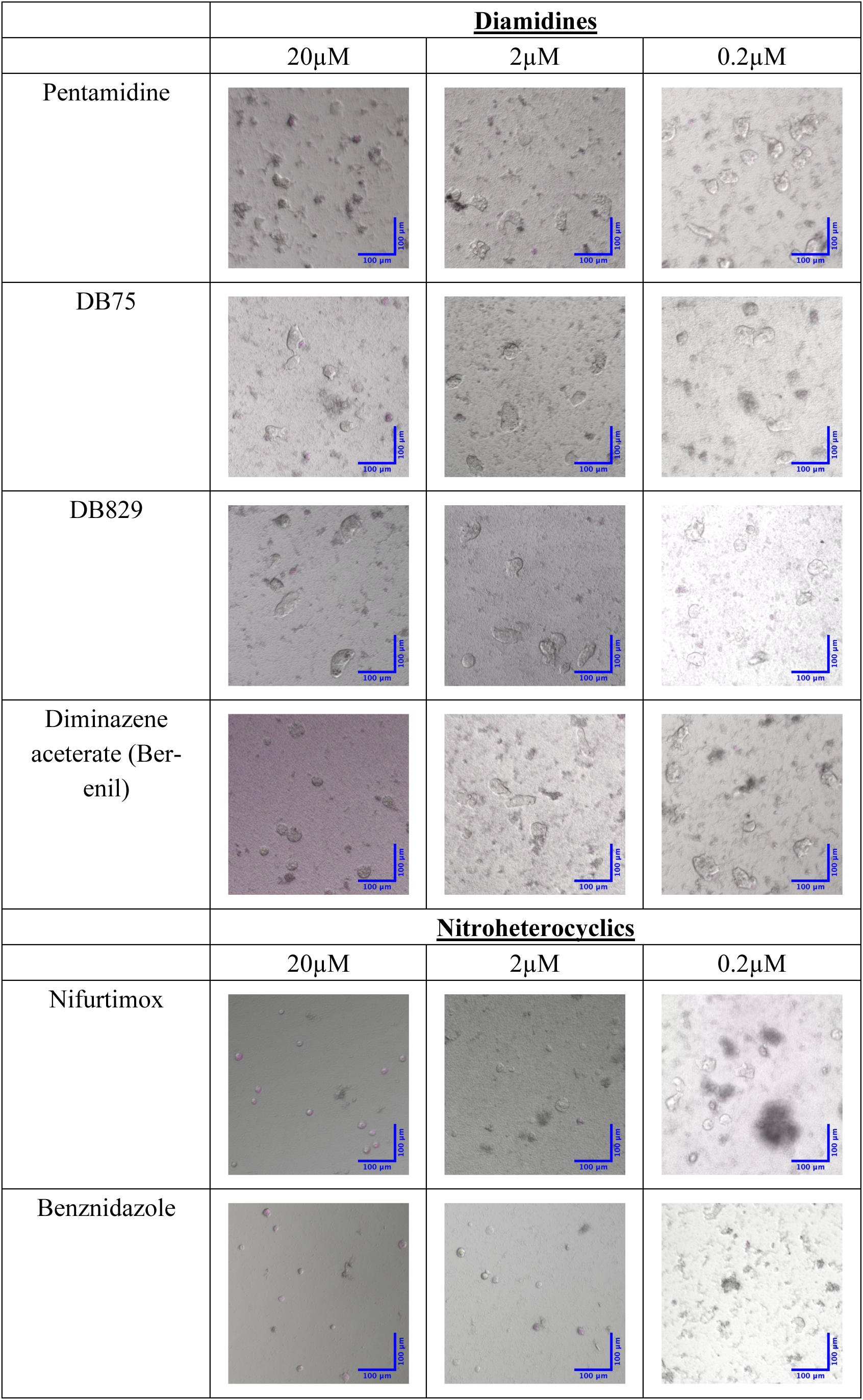
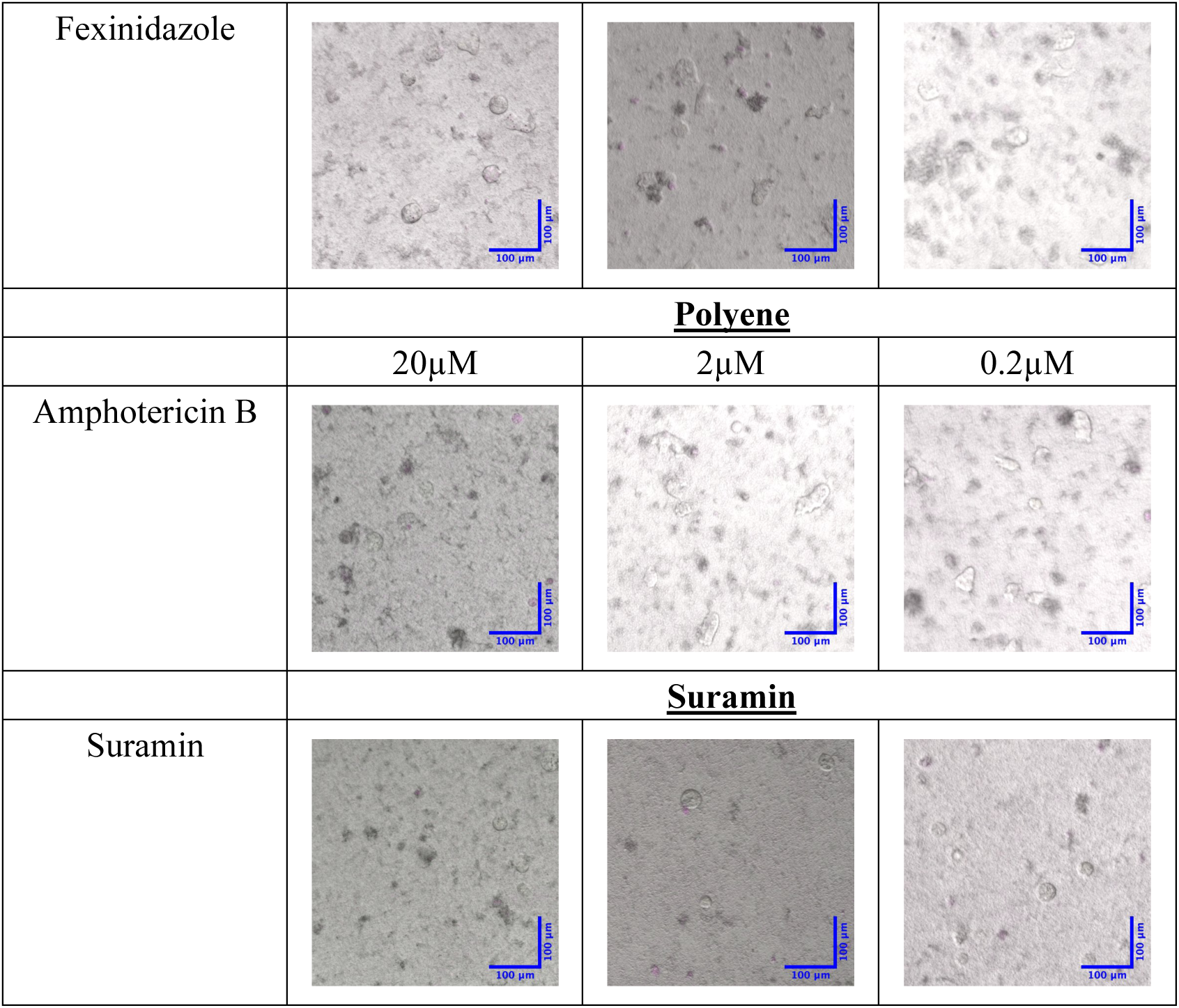
Fluorescence microscopy images of in vitro grown Neoparamoeba perurans following treatment with eight drug classes of trypanocidals at the concentrations of 20 µM, 2 µM, and 0.2 µM after 72 hours. Drug treated amoebae were stained with 20 µM of propidium iodide for 30 mins. Images were generated by merging the red fluorescent channel with the brightfield channel using a 10X objective. Propidium iodide stains dead cells red, while viable cells remain unstained.

### S5 Video recordings captured (with hyperlink) using the Holomonitor system to assess potential drug-induced changes in *Neoparmaoeba perurans* motility and morphology

**Table S5.**
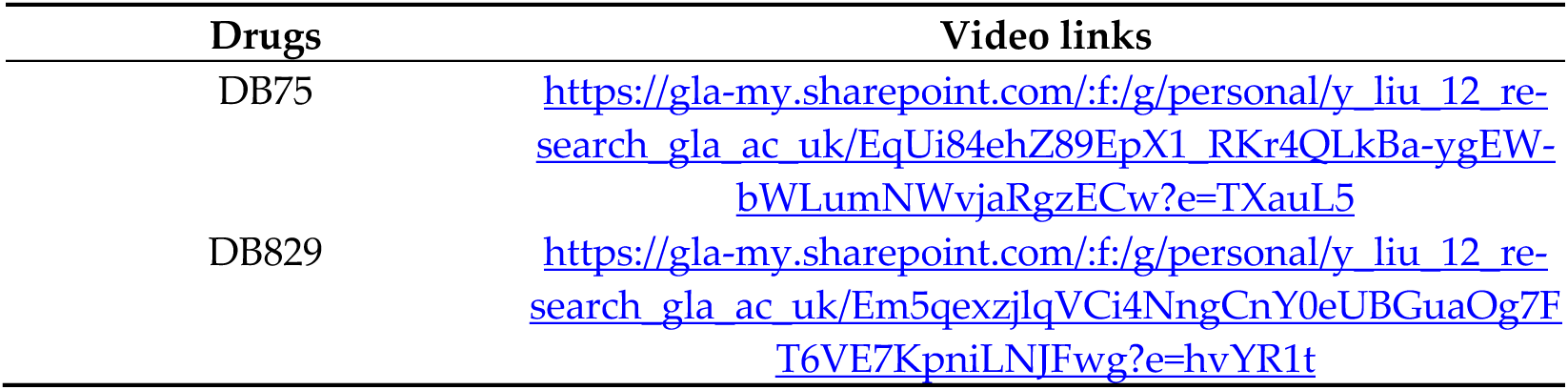
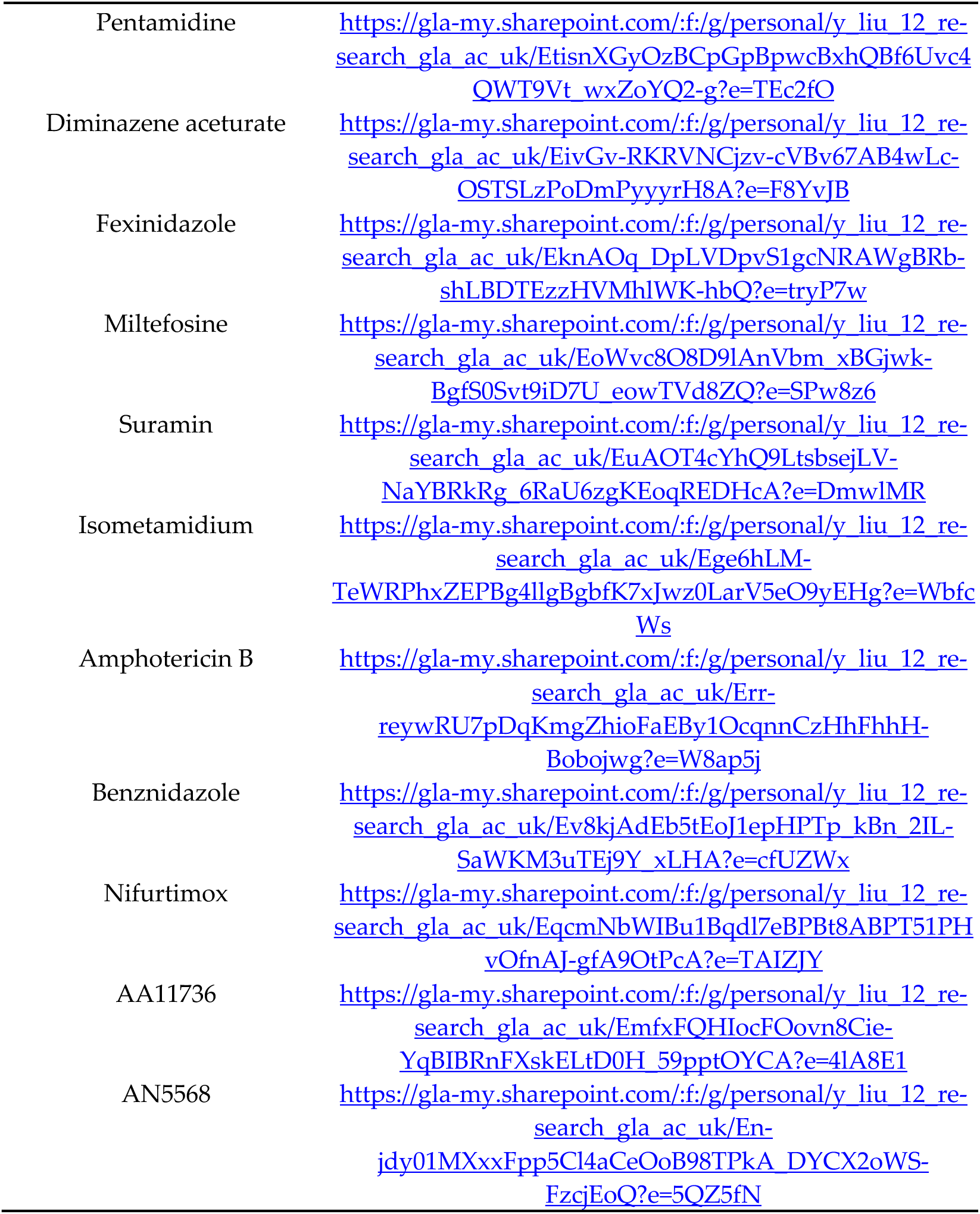
Video recordings captured (with hyperlink) using the Holomonitor system to assess potential drug-induced changes in *Neoparmaoeba perurans* motility and morphology over a 72 hour period following treatment with 13 different trypanocidal drugs. Each drug was tested at a concentration range of 20 µM to 0.625 µM. A drug is considered ineffective if no treatment-induced changes in motility or morphology were observed at any tested concentration, including the highest concentration of 20 µM.

## Notes

### Competing Interest Statement

The authors have declared no competing interest.

