## Supplementary Materials for "Repurposing drugs to treat Amoebic Gill Disease in Atlantic Salmon"

Article

Received: date

Revised: date

Accepted: date

Published: date

**Citation:** To be added by editorial staff during production.

**Copyright:** © 2025 by the authors. Submitted for possible open access publication under the terms and conditions of the Creative Commons Attribution (CC BY) license (https://creativecommons.org/licenses/by/4.0/).

1 School of Biodiversity, One Health and Veterinary Medicine, Graham Kerr Building, University of Glasgow, Glasgow, G12 8QQ;
2 Marine Insititute, Ireland
3 University College Cork, Ireland
4 Bantry Marine Research Station,
5 University of Strathclyde
6 Scottish Sea Farms Ltd
7 Universität Heidelberg

Supplementary material 1. S1

**Different concentration of Propidium iodide used and viewed under fluorescent microscope**

To identify the optimum concentration of propidium iodide if it can identify dead amoeba, propidium iodide of different concentration were added to formaldehyde fixed and no-treatment amoeba. As shown in Figure A, no staining occurs in the healthy amoeba but in the fixed amoeba which suggests that we can distinguish dead cells by propidium iodide. Furthermore, the optimal concentration of propidium iodide is 20µM as the fluorescence seems to be faint at lower concentrations but remains similar fluorescence level even in the concentration higher than 20µM. As a result, 20µM of propidium iodide was selected to be used in drug screening against N. perurans.

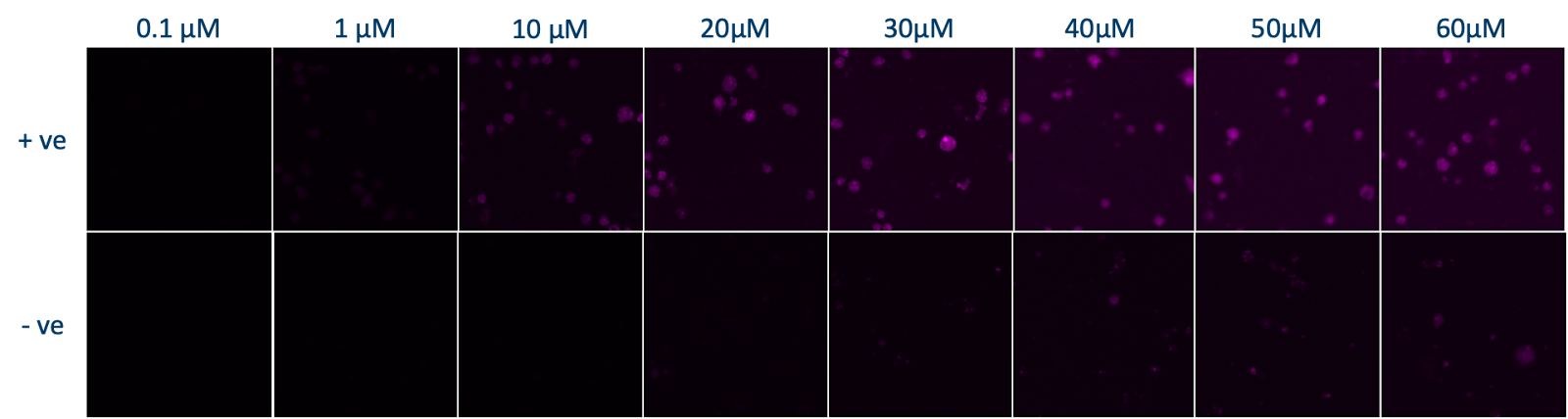

**Figure S1.** Formaldehyde killed Neoparamoeba perurans (+ve) and living Neoaramoeba perurans (-ve) were stained with different concentrations of propidium iodide from 0.1µM to 60µM.

Supplementary material 2. S2

Intracellular bacterial extraction using different methods

To develop effective lysing methods for eliminating amoebae while preserving the bacterial population from the non-axenically cultured *Neoparamoeba perurans*, three different methods were utilised: (1) manually breaking amoebae through rapid passaging with a gauged needle, water bath sonication, and the use of the isobiotec cell homogenizer. Despite attempting 50 passages with the fine gauge needle, it was insufficient in eliminating all amoebae (data not shown). (2) Similarly, 30 minutes of sonication also failed to eradicate all amoeba (data not shown). (3) The isobiotec cell homogeniser was particularly of interest due to its potential to break amoebae at 4 and 6 microns with 20 strokes (Figure A). By using isobiotec cell homogeniser, fluorescence and luminescence activity of the extracted extracellular and intracellular bacteria (hereafter homogenised amoebae) can be used as a background control, by comparing to the activity of non-axenically cultured *N. perurans* (hereafter non-homogenised amoebae), facilitating the estimation of the fluorescence and luminescence level originating from the amoebae themselves when conducting CellTox Green Cytotoxicity assay and CellTitre Glo Cell Viability assay.

| (**A**) | | (**B**) | | |
| --- | --- | --- | --- | --- |
| 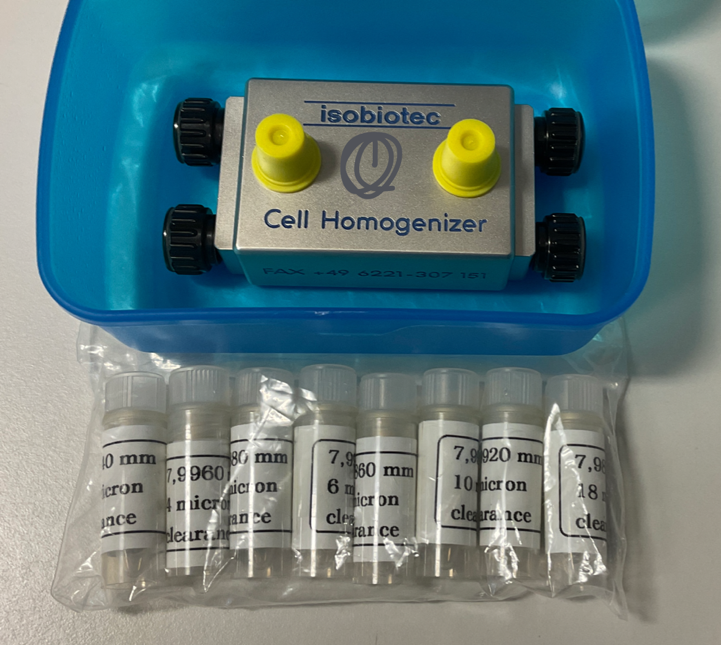 | | 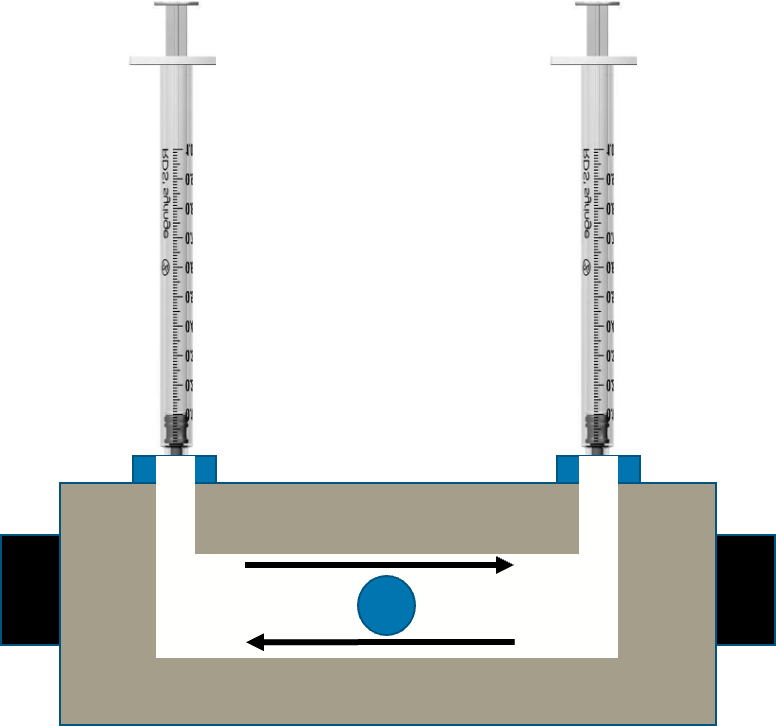 | | |
| (**C**) | | | | |
| Control | 8 microns | | 6 microns | 4 microns |
| 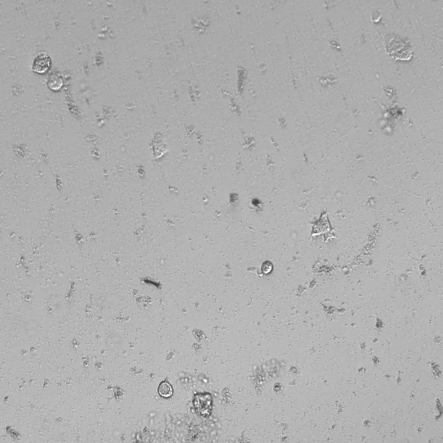 | 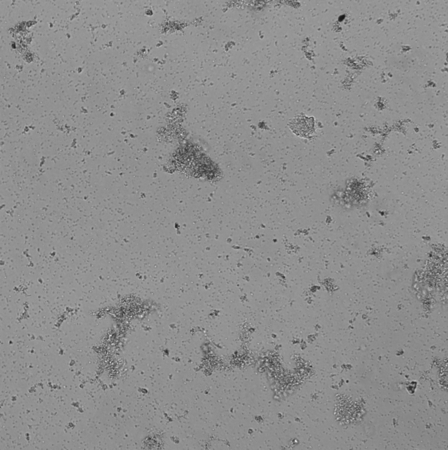 | | 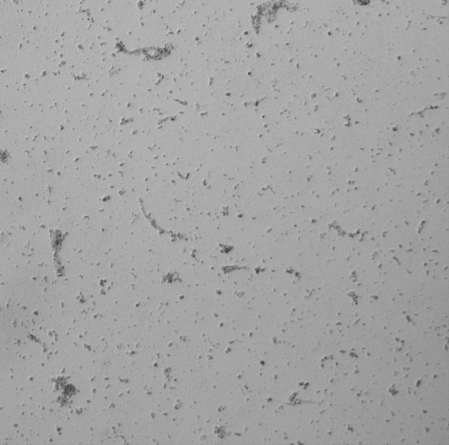 | 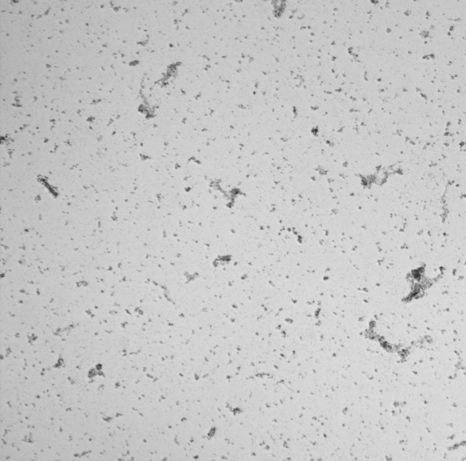 |

**Figure S2.** Preparation of intracellular and extracellular bacteria from *Neoparamoeba perurans* cultures using a Balch cell homogeniser. (A) Photograph of the isobiotec Balch homogeniser. (B) Schematic of the homogeniser chamber. A stainless-steel ball and an amoeba suspension in malt yeast broth were loaded into the chamber, then the sample was repeatedly forced through the narrow annular gap between the ball and the chamber wall to rupture cells and generate a homogenate. Balls of 8 μm, 6 μm and 4 μm diameter were used, and 20 strokes were applied. (C) Representative bright-field micrographs acquired at ×10 on a Leica DMi8 microscope, showing disrupted *N. perurans* cells of approximately 10–20 μm and release of associated bacteria.

Supplementary materials. S3.

Single Cell Tracking Analysis

**Miltefosine**

| **20 µM** | **10 µM** | **5 µM** | **2.5 µM** |
| --- | --- | --- | --- |
| 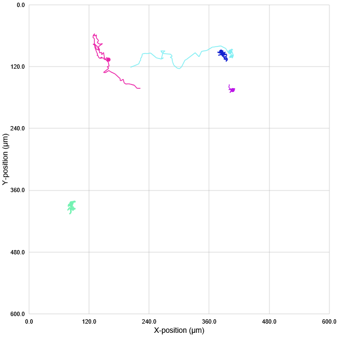 | 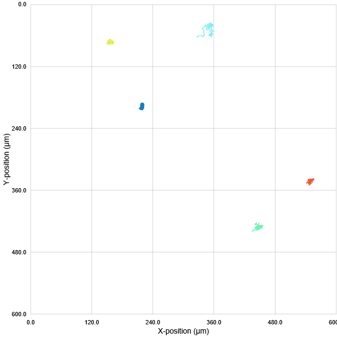 | 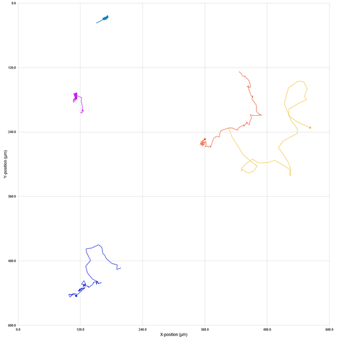 | 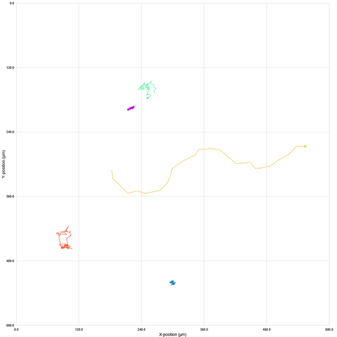 |
| **1.25 µM** | **0.626 µM** | **0 µM** |  |
| 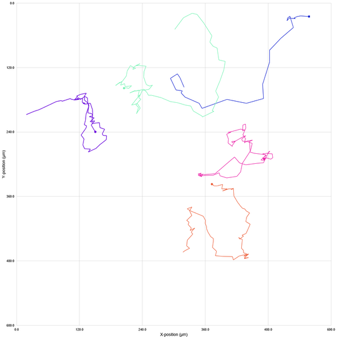 | 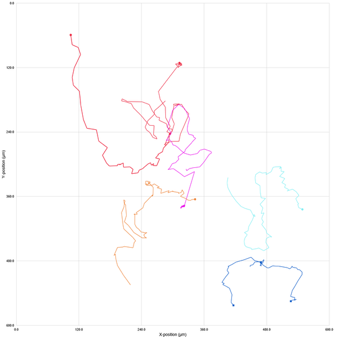 | 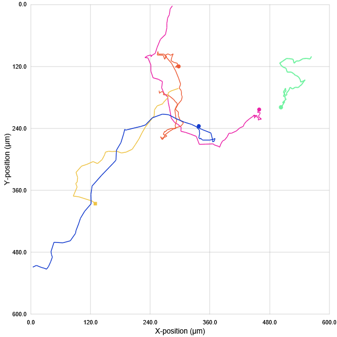 |  |

Single cell tracking analysis of *N. perurans* motility following exposure to miltefosine during a 4-hrs analysis windows at 0 h. The tested concentrations ranging from 20 µM to 0.625 µM. Each plot represents the movement of a single amoeba based on its X and Y coordinates captured by HoloMonitor at one randomly selected position within the well during the initial 6 hours of treatment. Lines depict the path each amoeba migrated during this period. Different colours indicate different amoeba being tracked.

| **20 µM** | **10 µM** | **5 µM** | **2.5 µM** |
| --- | --- | --- | --- |
| 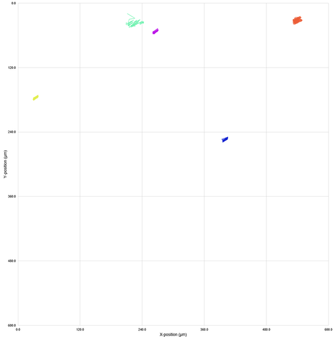 | 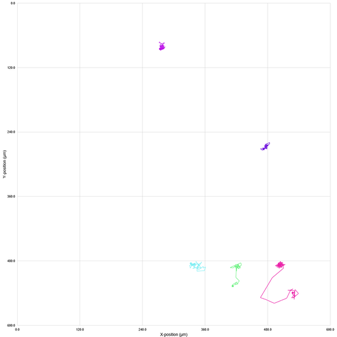 | 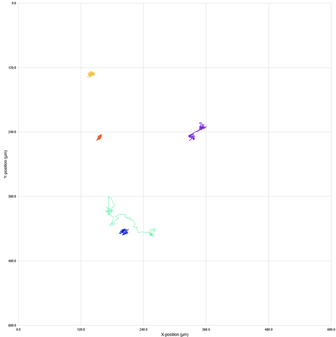 | 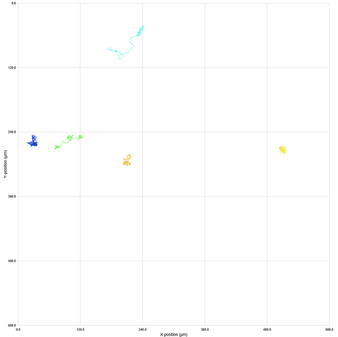 |
| **1.25 µM** | **0.626 µM** | **0 µM** |  |
| 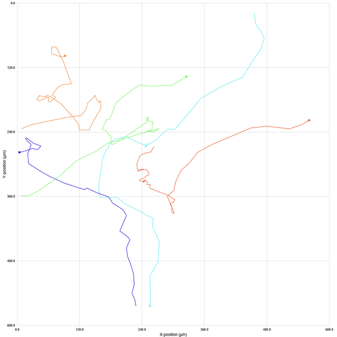 | 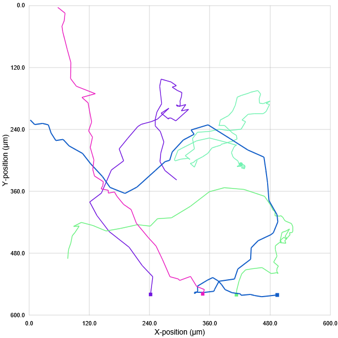 | 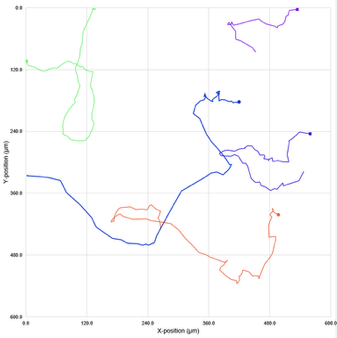 |  |

Single cell tracking analysis of *N. perurans* motility following exposure to miltefosine during a 4-hrs analysis windows at 24 h. The tested concentrations ranging from 20 µM to 0.625 µM. Each plot represents the movement of a single amoeba based on its X and Y coordinates captured by HoloMonitor at one randomly selected position within the well during the initial 6 hours of treatment. Lines depict the path each amoeba migrated during this period. Different colours indicate different amoeba being tracked.

| 20 µM | 10 µM | 5 µM | 2.5 µM |
| --- | --- | --- | --- |
| 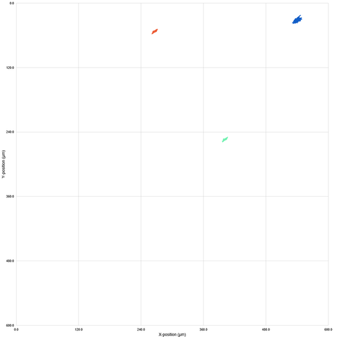 | 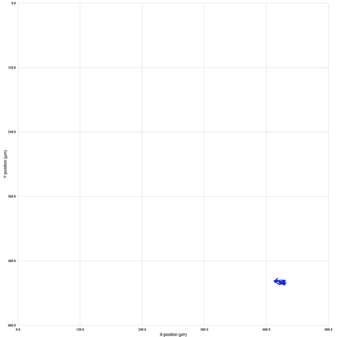 | 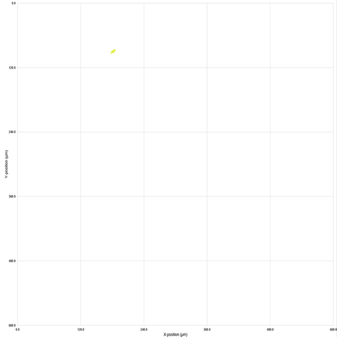 | 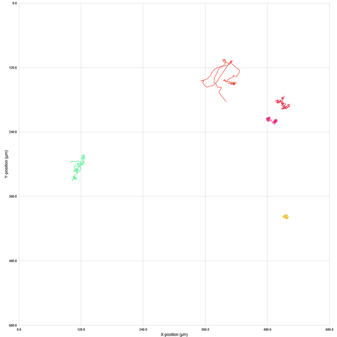 |
| 1.25 µM | 0.626 µM | 0 µM |  |
| 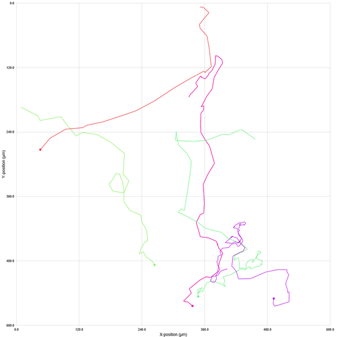 | 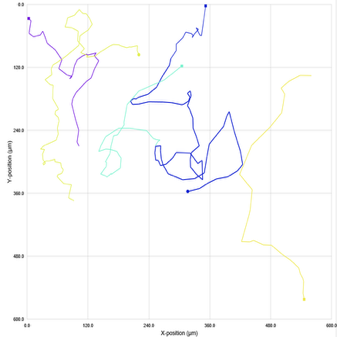 | 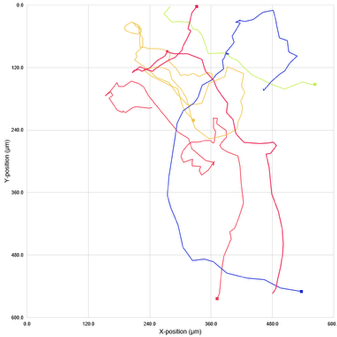 |  |

Single cell tracking analysis of N. perurans motility following exposure to miltefosine during a 4-hrs analysis windows at 48 h. The tested concentrations ranging from 20 µM to 0.625 µM. Each plot represents the movement of a single amoeba based on its X and Y coordinates captured by HoloMonitor at one randomly selected position within the well during the initial 6 hours of treatment. Lines depict the path each amoeba migrated during this period. Different colours indicate different amoeba being tracked.

| 20 µM | 10 µM | 5 µM | 2.5 µM |
| --- | --- | --- | --- |
| 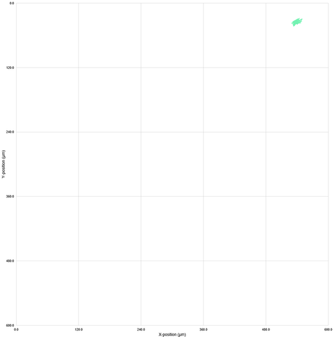 | No amoeba found | 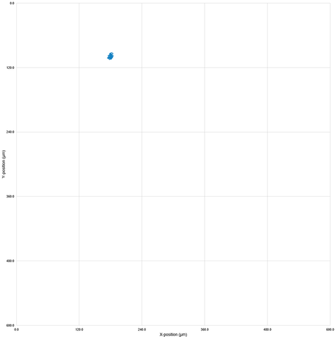 | 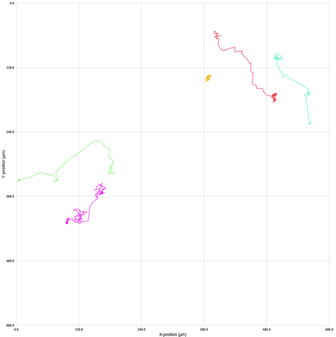 |
| 1.25 µM | 0.626 µM | 0 µM |  |
| 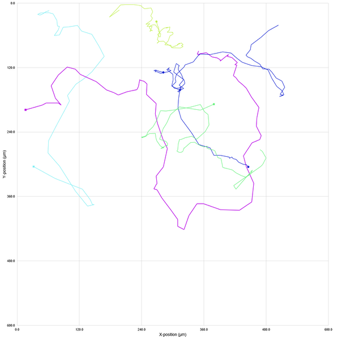 | 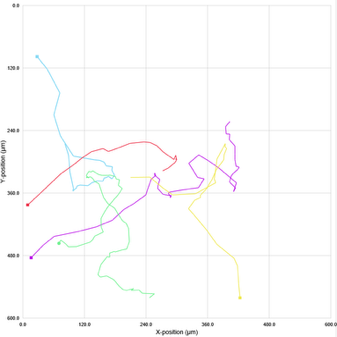 | 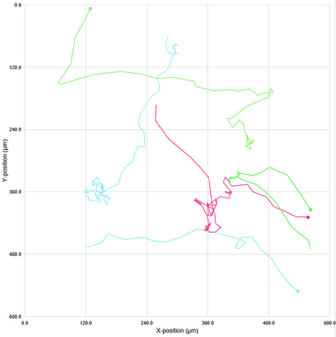 |  |

Single cell tracking analysis of N. perurans motility following exposure to miltefosine during a 4-hrs analysis windows at 72 h. The tested concentrations ranging from 20 µM to 0.625 µM. Each plot represents the movement of a single amoeba based on its X and Y coordinates captured by HoloMonitor at one randomly selected position within the well during the initial 6 hours of treatment. Lines depict the path each amoeba migrated during this period. Different colours indicate different amoeba being tracked.

Isometamidium

| 20 µM | 10 µM | 5 µM | 2.5 µM |
| --- | --- | --- | --- |
| 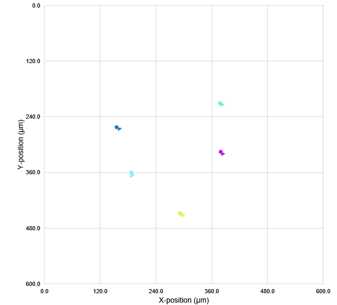 | 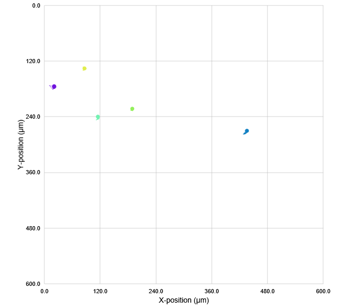 | 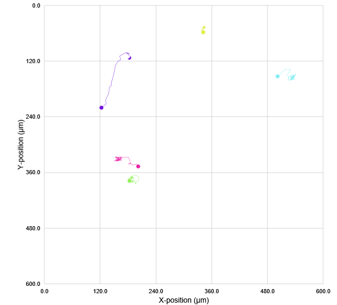 | 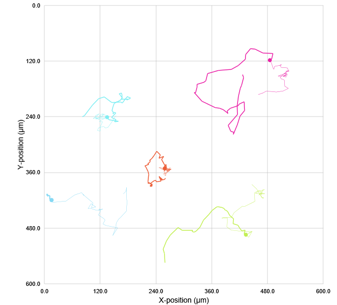 |
| 1.25 µM | 0.626 µM | 0 µM |  |
| 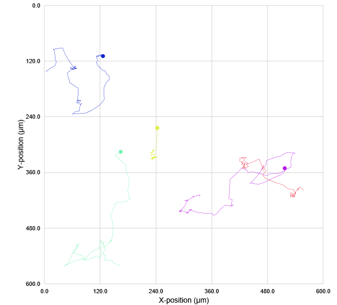 | 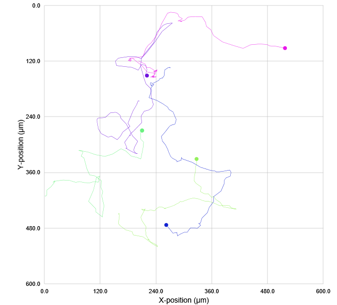 | 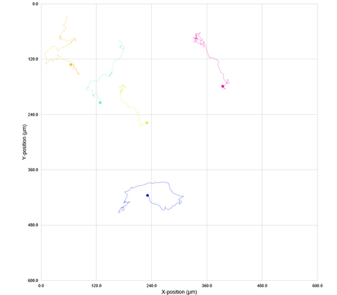 |  |

Single cell tracking analysis of N. perurans motility following exposure to isometamidium during a 4-hrs analysis windows at 0 h. The tested concentrations ranging from 20 µM to 0.625 µM. Each plot represents the movement of a single amoeba based on its X and Y coordinates captured by HoloMonitor at one randomly selected position within the well during the initial 6 hours of treatment. Lines depict the path each amoeba migrated during this period. Different colours indicate different amoeba being tracked.

| 20 µM | 10 µM | 5 µM | 2.5 µM |
| --- | --- | --- | --- |
| 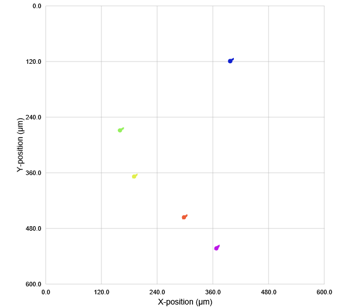 | 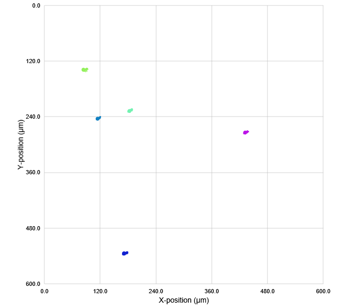 | 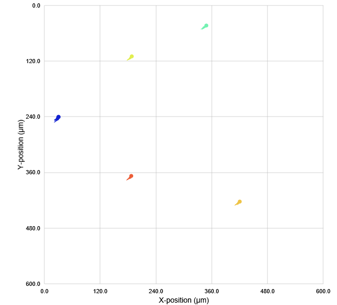 | 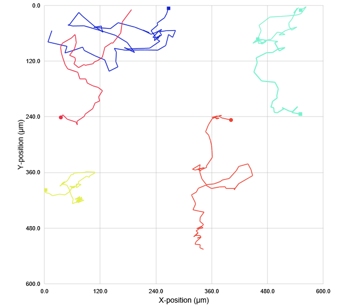 |
| 1.25 µM | 0.626 µM | 0 µM |  |
| 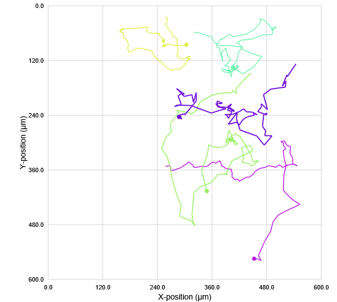 | 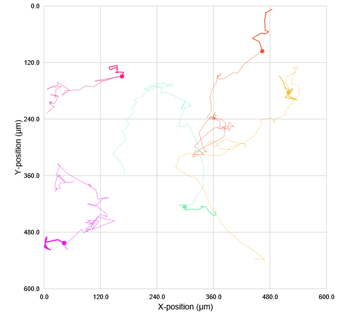 | 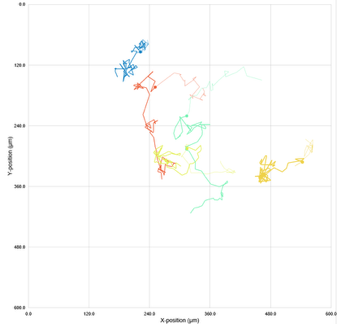 |  |

Single cell tracking analysis of *N. perurans* motility following exposure to isometamidium during a 4-hrs analysis windows at 24 h. The tested concentrations ranging from 20 µM to 0.625 µM. Each plot represents the movement of a single amoeba based on its X and Y coordinates captured by HoloMonitor at one randomly selected position within the well during the initial 6 hours of treatment. Lines depict the path each amoeba migrated during this period. Different colours indicate different amoeba being tracked.

| 20 µM | 10 µM | 5 µM | 2.5 µM |
| --- | --- | --- | --- |
| 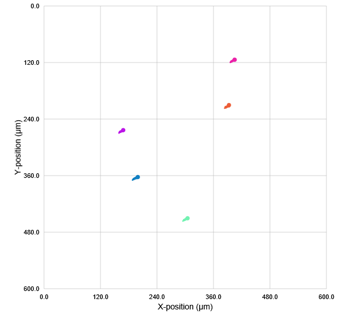 | 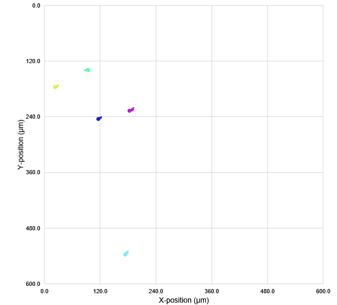 | 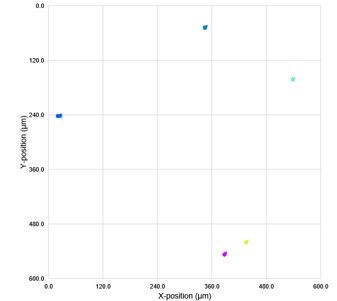 | 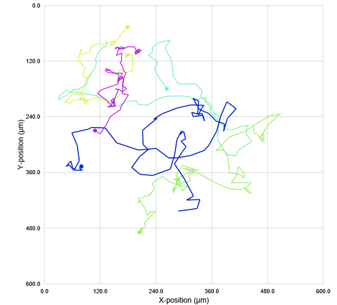 |
| 1.25 µM | 0.626 µM | 0 µM |  |
| 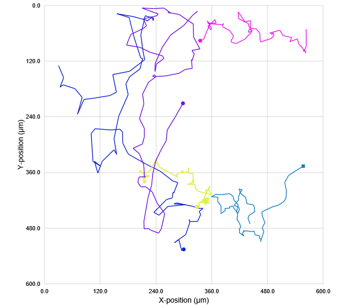 | 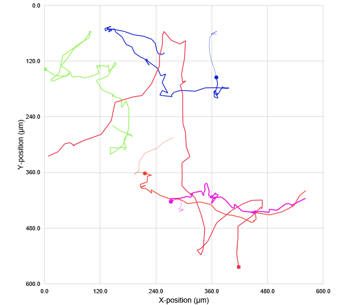 | 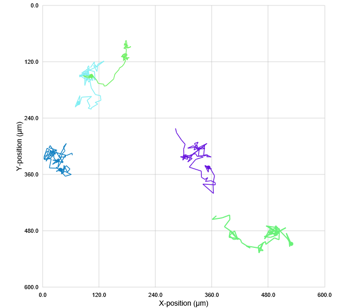 |  |

Single cell tracking analysis of *N. perurans* motility following exposure to isometamidium during a 4-hrs analysis windows at 48 h. The tested concentrations ranging from 20 µM to 0.625 µM. Each plot represents the movement of a single amoeba based on its X and Y coordinates captured by HoloMonitor at one randomly selected position within the well during the initial 6 hours of treatment. Lines depict the path each amoeba migrated during this period. Different colours indicate different amoeba being tracked.

| 20 µM | 10 µM | 5 µM | 2.5 µM |
| --- | --- | --- | --- |
| 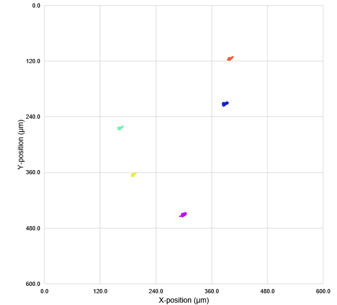 | 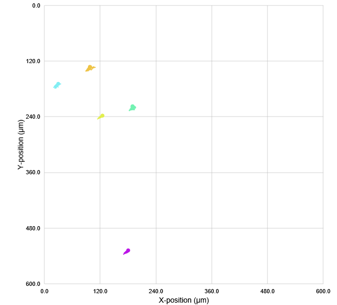 | 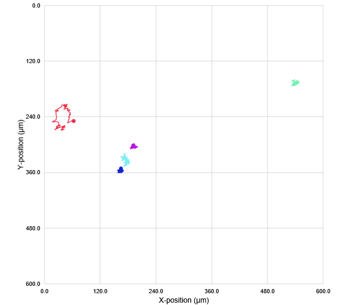 | 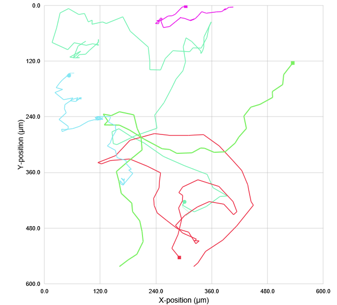 |
| 1.25 µM | 0.626 µM | 0 µM |  |
| 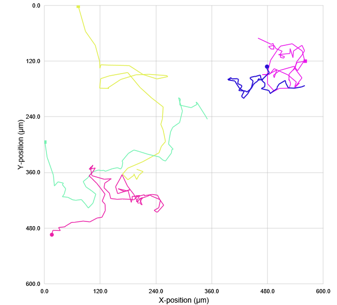 | 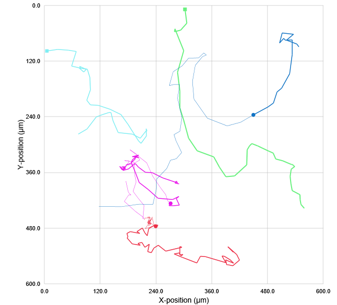 | 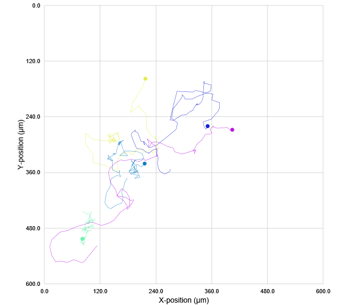 |  |

Single cell tracking analysis of *N. perurans* motility following exposure to isometamidium during a 4-hrs analysis windows at 72 h. The tested concentrations ranging from 20 µM to 0.625 µM. Each plot represents the movement of a single amoeba based on its X and Y coordinates captured by HoloMonitor at one randomly selected position within the well during the initial 6 hours of treatment. Lines depict the path each amoeba migrated during this period. Different colours indicate different amoeba being tracked.

**AN11736**

| 20 µM | 10 µM | 5 µM | 2.5 µM |
| --- | --- | --- | --- |
| 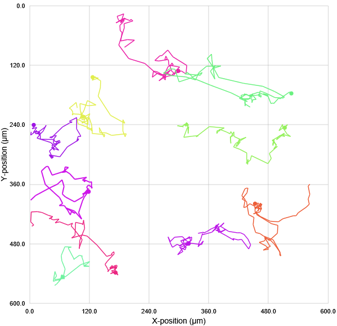 | 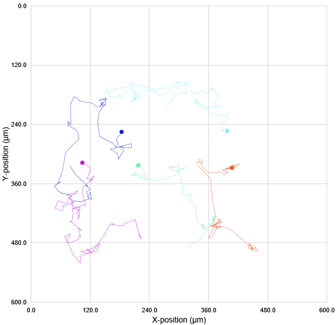 | 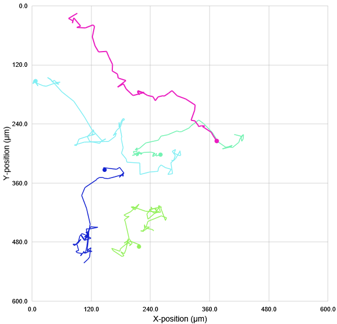 | 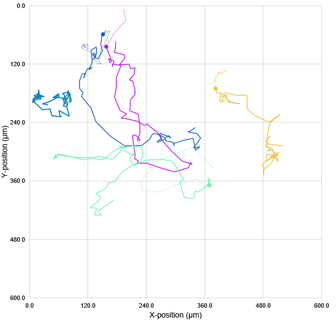 |
| 1.25 µM | 0.626 µM | 0 µM |  |
| 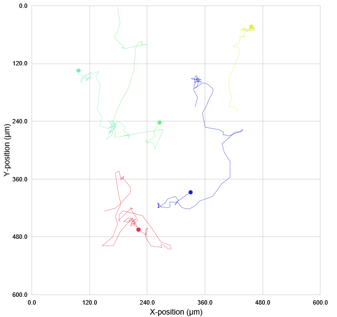 | 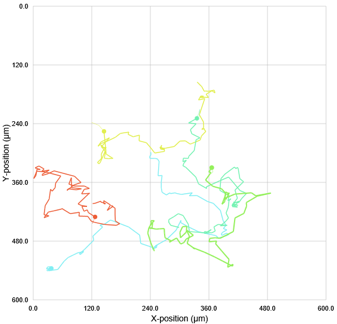 | 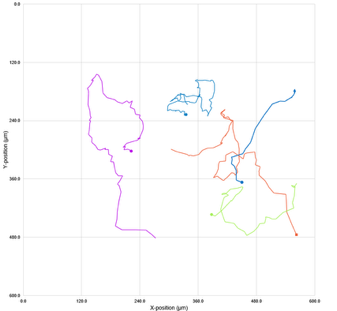 |  |

Single cell tracking analysis of *N. perurans* motility following exposure to AN11736 during a 4-hrs analysis windows at 0 h. The tested concentrations ranging from 20 µM to 0.625 µM.Each plot represents the movement of a single amoeba based on its X and Y coordinates captured by HoloMonitor at one randomly selected position within the well during the initial 6 hours of treatment. Lines depict the path each amoeba migrated during this period. Different colours indicate different amoeba being tracked.

| 20 µM | 10 µM | 5 µM | 2.5 µM |
| --- | --- | --- | --- |
| 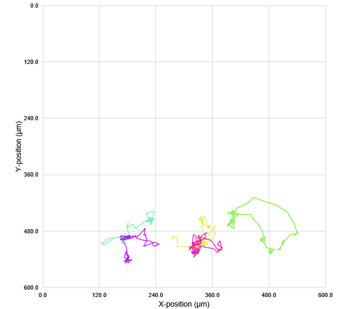 | 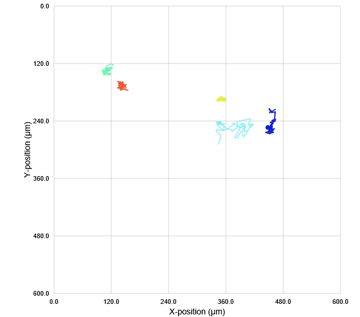 | 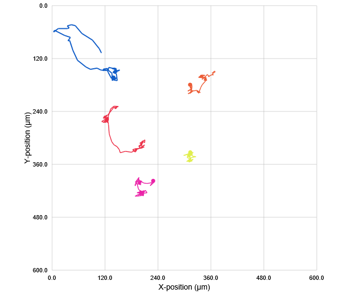 | 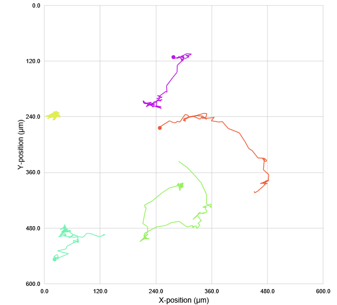 |
| 1.25 µM | 0.626 µM | 0 µM |  |
| 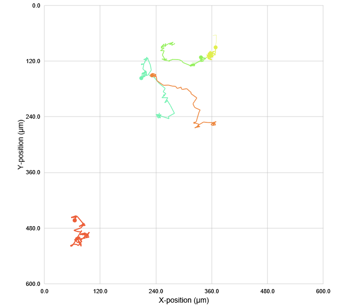 | 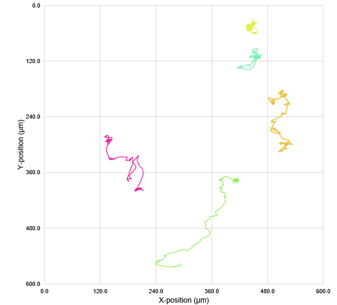 | 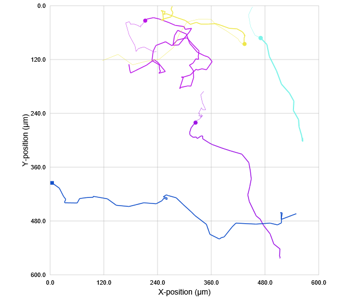 |  |

Single cell tracking analysis of *N. perurans* motility following exposure to AN11736 during a 4-hrs analysis windows at 24 h. The tested concentrations ranging from 20 µM to 0.625 µM.Each plot represents the movement of a single amoeba based on its X and Y coordinates captured by HoloMonitor at one randomly selected position within the well during the initial 6 hours of treatment. Lines depict the path each amoeba migrated during this period. Different colours indicate different amoeba being tracked.

| 20 µM | 10 µM | 5 µM | 2.5 µM |
| --- | --- | --- | --- |
| 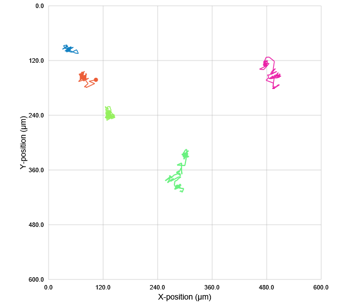 | 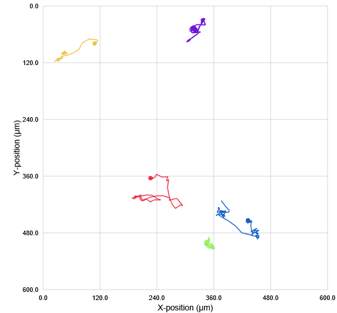 | 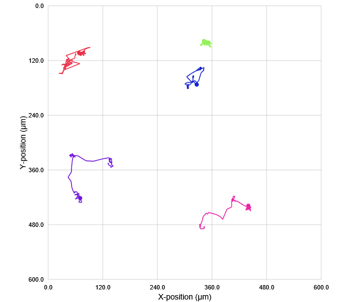 | 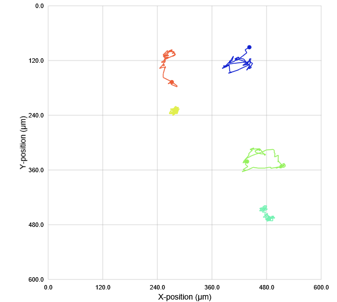 |
| 1.25 µM | 0.626 µM | 0 µM |  |
| 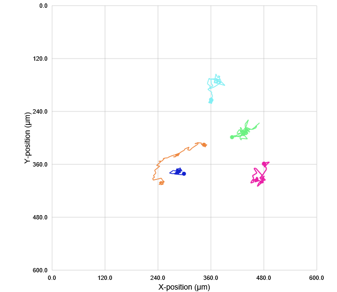 | 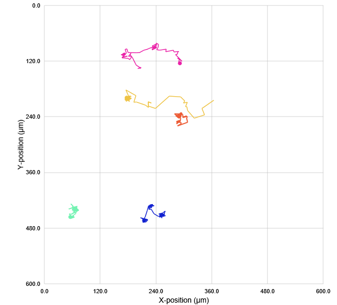 | 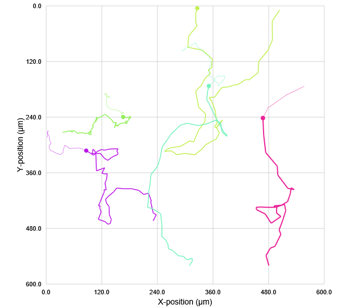 |  |

Single cell tracking analysis of *N. perurans* motility following exposure to AN11736 during a 4-hrs analysis windows at 48 h. The tested concentrations ranging from 20 µM to 0.625 µM. Each plot represents the movement of a single amoeba based on its X and Y coordinates captured by HoloMonitor at one randomly selected position within the well during the initial 6 hours of treatment. Lines depict the path each amoeba migrated during this period. Different colours indicate different amoeba being tracked.

| 20 µM | 10 µM | 5 µM | 2.5 µM |
| --- | --- | --- | --- |
| 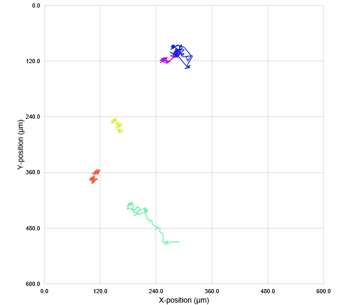 | 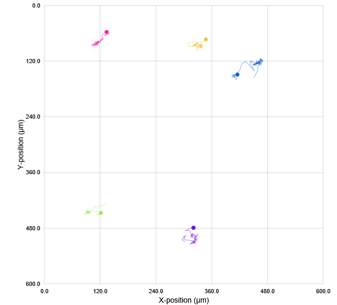 | 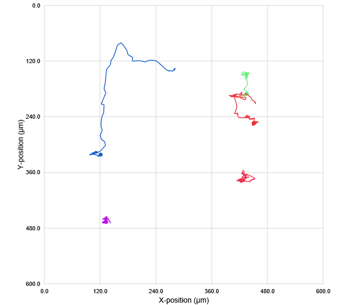 | 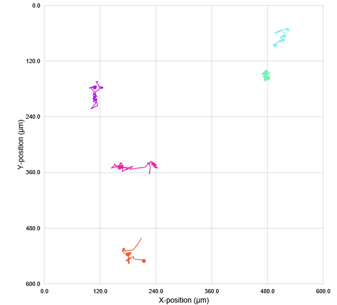 |
| 1.25 µM | 0.626 µM | 0 µM |  |
| 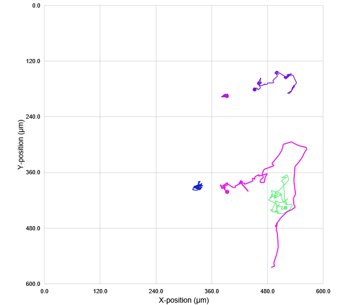 | 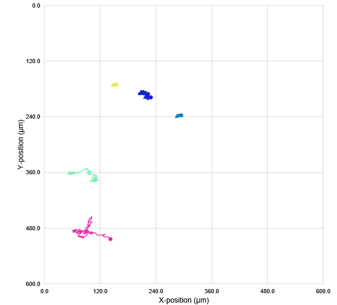 | 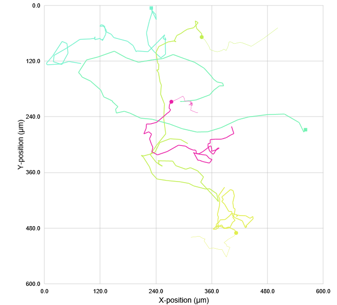 |  |

Single cell tracking analysis of *N. perurans* motility following exposure to AN11736 during a 4-hrs analysis windows at 72 h. The tested concentrations ranging from 20 µM to 0.625 µM. Each plot represents the movement of a single amoeba based on its X and Y coordinates captured by HoloMonitor at one randomly selected position within the well during the initial 6 hours of treatment. Lines depict the path each amoeba migrated during this period. Different colours indicate different amoeba being tracked.

**Pentamidine**

| 20 µM | 10 µM | 5 µM | 2.5 µM |
| --- | --- | --- | --- |
| 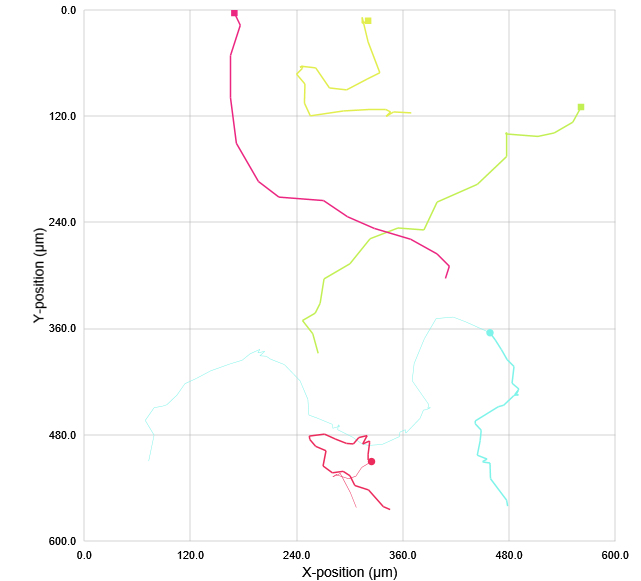 | 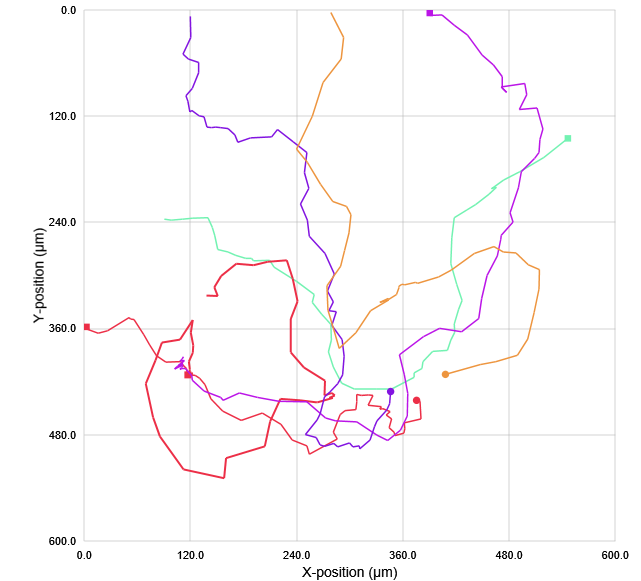 | 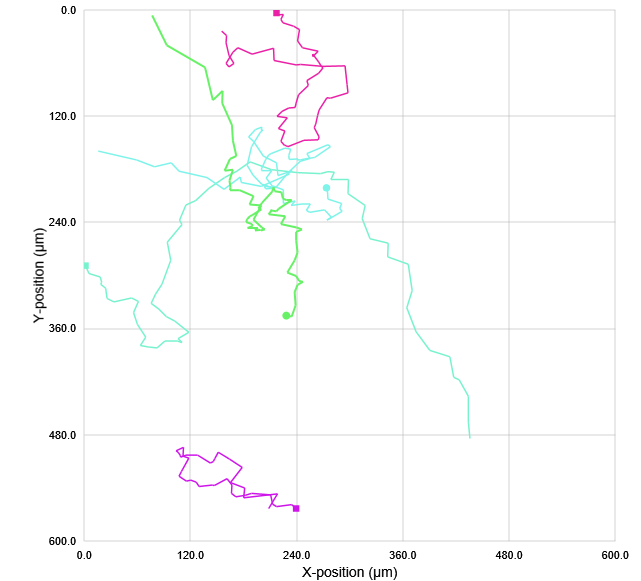 | 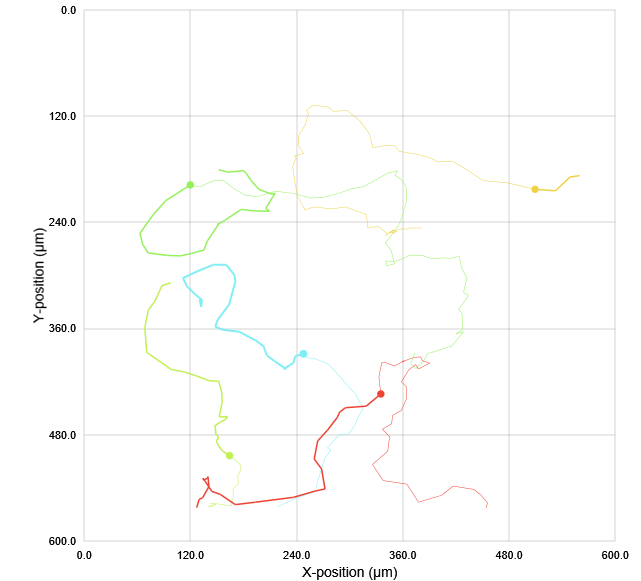 |
| 1.25 µM | 0.626 µM | 0 µM |  |
| 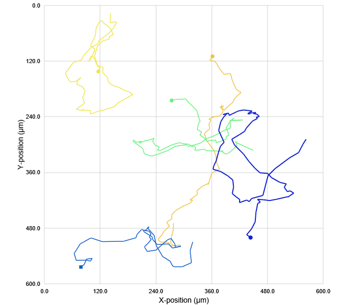 | 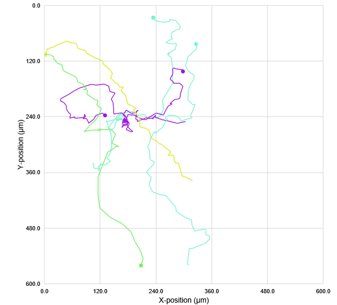 | 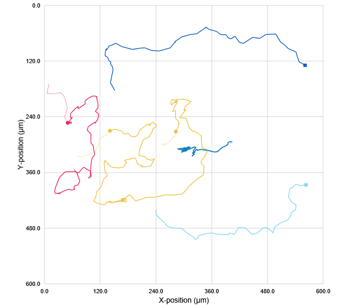 |  |

Single cell tracking analysis of *N. perurans* motility following exposure to pentamidine during a 4-hrs analysis windows at 0 h. The tested concentrations ranging from 20 µM to 0.625 µM. Each plot represents the movement of a single amoeba based on its X and Y coordinates captured by HoloMonitor at one randomly selected position within the well during the initial 6 hours of treatment. Lines depict the path each amoeba migrated during this period. Different colours indicate different amoeba being tracked.

| 20 µM | 10 µM | 5 µM | 2.5 µM |
| --- | --- | --- | --- |
| 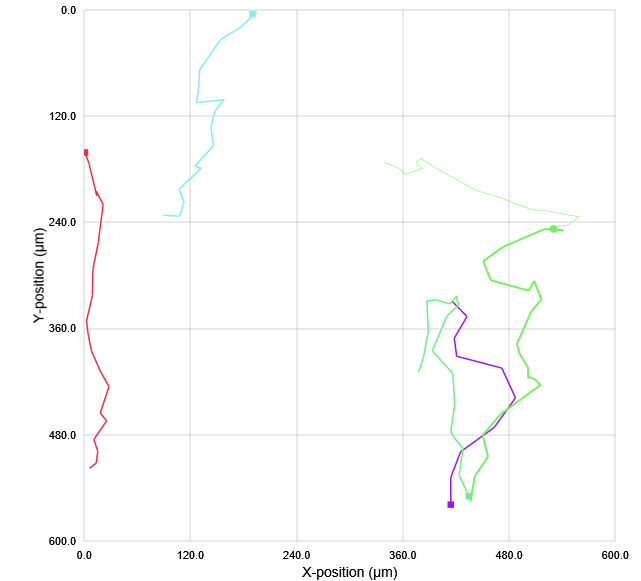 | 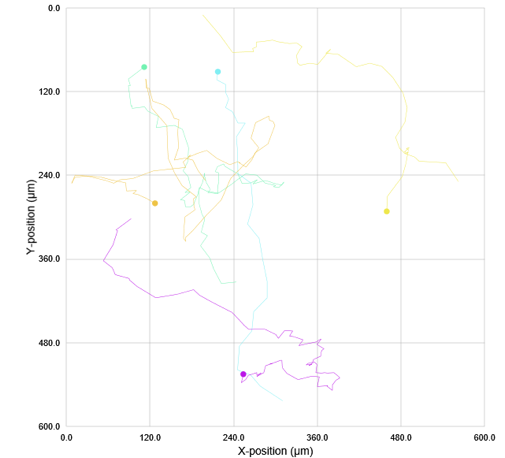 | 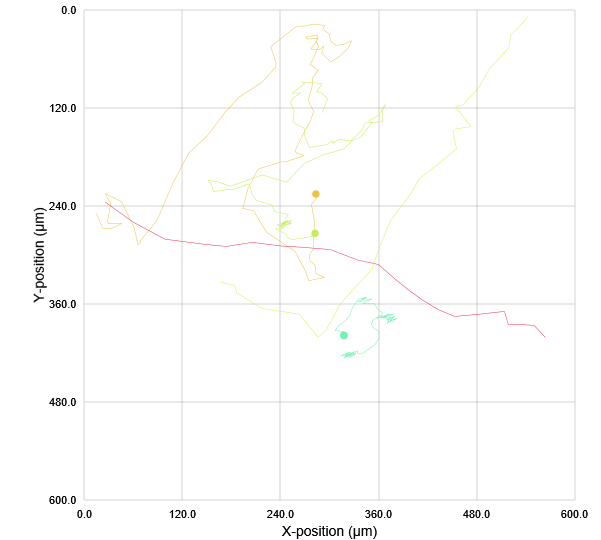 |  |
| 1.25 µM | 0.626 µM | 0 µM |  |

Single cell tracking analysis of *N. perurans* motility following exposure to pentamidine during a 4-hrs analysis windows at 24 h. The tested concentrations ranging from 20 µM to 0.625 µM. Each plot represents the movement of a single amoeba based on its X and Y coordinates captured by HoloMonitor at one randomly selected position within the well during the initial 6 hours of treatment. Lines depict the path each amoeba migrated during this period. Different colours indicate different amoeba being tracked.

Supplementary materials 4, S4

Fluorescence microscopy images of in vitro grown Neoparamoeba perurans following treatment with eight drug classes of trypanocidals at different concentrations.

|  | **Alkylphosphocholine** | | |
| --- | --- | --- | --- |
|  | 20µM | 2µM | 0.2µM |
| Miltefosine |  |  |  |
|  | **Phenathridine** | | |
|  | 20µM | 2µM | 0.2µM |
| Isometamidium |  |  |  |
|  | **Aminoglycoside** | | |
|  | 20µM | 2µM | 0.2µM |
| Paromomycin |  |  |  |
|  | **Benzoxaboroles** | | |
|  | 20µM | 2µM | 0.2µM |
| AA11736 (valylester) |  |  |  |
| AN5568 |  |  |  |

|  | **Diamidines** | | |
| --- | --- | --- | --- |
|  | 20µM | 2µM | 0.2µM |
| Pentamidine |  |  |  |
| DB75 |  |  |  |
| DB829 |  |  |  |
| Diminazene aceterate (Berenil) |  |  |  |
|  | **Nitroheterocyclics** | | |
|  | 20µM | 2µM | 0.2µM |
| Nifurtimox |  |  |  |
| Benznidazole |  |  |  |
| Fexinidazole |  |  |  |
|  | **Polyene** | | |
|  | 20µM | 2µM | 0.2µM |
| Amphotericin B |  |  |  |
|  | **Suramin** | | |
| Suramin |  |  |  |

**Figure S4.** Fluorescence microscopy images of in vitro grown Neoparamoeba perurans following treatment with eight drug classes of trypanocidals at the concentrations of 20 µM, 2 µM, and 0.2 µM after 72 hours. Drug treated amoebae were stained with 20 µM of propidium iodide for 30 mins. Images were generated by merging the red fluorescent channel with the brightfield channel using a 10X objective. Propidium iodide stains dead cells red, while viable cells remain unstained.

Supplementary Material S5

Video recordings captured (with hyperlink) using the Holomonitor system to assess potential drug-induced changes in *Neoparmaoeba perurans* motility and morphology

**Table S5.** Video recordings captured (with hyperlink) using the Holomonitor system to assess potential drug-induced changes in *Neoparmaoeba perurans* motility and morphology over a 72 hour period following treatment with 13 different trypanocidal drugs. Each drug was tested at a concentration range of 20 µM to 0.625 µM. A drug is considered ineffective if no treatment-induced changes in motility or morphology were observed at any tested concentration, including the highest concentration of 20 µM.

| **Drugs** | **Video links** |
| --- | --- |
| DB75 | <https://gla-my.sharepoint.com/:f:/g/personal/y_liu_12_research_gla_ac_uk/EqUi84ehZ89EpX1_RKr4QLkBa-ygEWbWLumNWvjaRgzECw?e=TXauL5> |
| DB829 | <https://gla-my.sharepoint.com/:f:/g/personal/y_liu_12_research_gla_ac_uk/Em5qexzjlqVCi4NngCnY0eUBGuaOg7FT6VE7KpniLNJFwg?e=hvYR1t> |
| Pentamidine | <https://gla-my.sharepoint.com/:f:/g/personal/y_liu_12_research_gla_ac_uk/EtisnXGyOzBCpGpBpwcBxhQBf6Uvc4QWT9Vt_wxZoYQ2-g?e=TEc2fO> |
| Diminazene aceturate | <https://gla-my.sharepoint.com/:f:/g/personal/y_liu_12_research_gla_ac_uk/EivGv-RKRVNCjzv-cVBv67AB4wLcOSTSLzPoDmPyyyrH8A?e=F8YvJB> |
| Fexinidazole | <https://gla-my.sharepoint.com/:f:/g/personal/y_liu_12_research_gla_ac_uk/EknAOq_DpLVDpvS1gcNRAWgBRbshLBDTEzzHVMhlWK-hbQ?e=tryP7w> |
| Miltefosine | <https://gla-my.sharepoint.com/:f:/g/personal/y_liu_12_research_gla_ac_uk/EoWvc8O8D9lAnVbm_xBGjwkBgfS0Svt9iD7U_eowTVd8ZQ?e=SPw8z6> |
| Suramin | <https://gla-my.sharepoint.com/:f:/g/personal/y_liu_12_research_gla_ac_uk/EuAOT4cYhQ9LtsbsejLVNaYBRkRg_6RaU6zgKEoqREDHcA?e=DmwlMR> |
| Isometamidium | <https://gla-my.sharepoint.com/:f:/g/personal/y_liu_12_research_gla_ac_uk/Ege6hLMTeWRPhxZEPBg4llgBgbfK7xJwz0LarV5eO9yEHg?e=WbfcWs> |
| Amphotericin B | <https://gla-my.sharepoint.com/:f:/g/personal/y_liu_12_research_gla_ac_uk/ErrreywRU7pDqKmgZhioFaEBy1OcqnnCzHhFhhHBobojwg?e=W8ap5j> |
| Benznidazole | <https://gla-my.sharepoint.com/:f:/g/personal/y_liu_12_research_gla_ac_uk/Ev8kjAdEb5tEoJ1epHPTp_kBn_2ILSaWKM3uTEj9Y_xLHA?e=cfUZWx> |
| Nifurtimox | <https://gla-my.sharepoint.com/:f:/g/personal/y_liu_12_research_gla_ac_uk/EqcmNbWIBu1Bqdl7eBPBt8ABPT51PHvOfnAJ-gfA9OtPcA?e=TAIZJY> |
| AA11736 | <https://gla-my.sharepoint.com/:f:/g/personal/y_liu_12_research_gla_ac_uk/EmfxFQHIocFOovn8CieYqBIBRnFXskELtD0H_59pptOYCA?e=4lA8E1> |
| AN5568 | <https://gla-my.sharepoint.com/:f:/g/personal/y_liu_12_research_gla_ac_uk/Enjdy01MXxxFpp5Cl4aCeOoB98TPkA_DYCX2oWSFzcjEoQ?e=5QZ5fN> |
